# The evolution of reversible plasticity in stable environments

**DOI:** 10.1101/2024.08.27.609887

**Authors:** Nicole Walasek, Karthik Panchanathan, Willem E. Frankenhuis

**Author notes:** **Conflict of Interest:** The authors declare no conflict of interest. **Author Contributions:** NW and WEF conceived of the study. NW developed the model and analyzed the resulting data. NW wrote the first draft. NW, WEF, and KP revised the draft. **Data Availability Statement:** No data were collected or used for this paper.

## Abstract

*Phenotypic plasticity* – the capacity of a genotype to develop into different phenotypes depending on environmental inputs – is widespread in nature. Although the *construction* of phenotypes is often the focus of research, many animals are also able to *deconstruct* phenotypic adjustments. Organisms routinely use such *reversible plasticity* to adjust to changes in their social or physical environment. For example, various invertebrates are able to deconstruct defensive morphologies previously built to defend against predators. Theory that explores the selection pressures favoring reversibility is scarce. Existing theory has almost exclusively focused on traits that develop instantaneously rather than incrementally, as is common with many morphological traits. Here, we present a model of the evolution of reversible plasticity when organisms develop incrementally. In our model, organisms repeatedly sample cues to infer the environmental state – which varies between generations but is stable across the lifetime – and incrementally adjust their phenotype to match their environment. Organisms have the possibility to deconstruct phenotypic adjustments. We assume two different modes of phenotypic deconstruction: Organisms can either deconstruct phenotypic adjustments incrementally or completely deconstruct all phenotypic adjustments in one time period. We highlight two results. First, while plasticity in construction is typically highest early in ontogeny, the highest levels of plasticity in deconstruction typically occur in mid-ontogeny. Second, contrasting previous models, we find that reversibility evolves frequently in stable environments and in species with shorter lifespans. Our model thus shows that reversibility does not require environmental change. Rather, reversibility may be favored when organisms are uncertain about the environmental state because the environment can change across generations. Our work illustrates the capacity for reversibility in species who experience environmental changes for the first time in their lives.

## Introduction

### Reversible plasticity is common

Phenotypic plasticity – the capacity of a genotype to develop into different phenotypes depending on environmental inputs (Nettle & Bateson, 2015; Stearns, 1989; West-Eberhard, 2003) – is widespread in nature. Although the *construction* of phenotypes is often the focus of research, many animals are also able to *deconstruct* phenotypic adjustments. Organisms routinely use such *reversible plasticity* to adjust to changes in their social or physical environment (Burggren, 2020). For example, the African cichlid fish (*Astatotilapia burtoni*) shows high reversibility both in behavior and morphology in response to their social environment (Maruska & Fernald, 2013): Depending on their dominance rank, male fish either express bright coloring with an active reproductive strategy or drab coloring with suppressed reproduction; males are able to switch between these phenotypes repeatedly across their lifetimes. Another remarkable example of reversible plasticity in response to changes in the social environment are reversible sex changes in gobies (Oyama et al., 2023). In the absence of males several goby species are able to change sex from female to male while retaining their ovarian tissues (Lorenzi et al., 2006; Oyama et al., 2023; Sunobe & Nakazono, 1993). Once a more dominant male appears, these sex-changed males revert back to being female. Reversible plasticity is also commonly observed in response to changes in food availability (Utz et al., 2014). An extreme example is the capacity of marine iguanas (*Amblyrhynchus cristatus*) to shrink body length by 20 percent to avoid starvation following El Niño events and to reverse this shrinkage during La Niña events (Wikelski & Thom, 2000). Traditionally, reversible phenotypic plasticity is equated with the life-long capacity to fully reverse phenotypes (Burggren, 2020; Piersma & Drent, 2003). While the examples above fit this definition, many others do not. This is especially apparent for morphological defenses. For instance, freshwater snails (*Helisoma trivolis*) are able to decrease the thickness of shells previously increased as a defense against predators (Hoverman & Relyea, 2007).

However, only when predators were removed early in ontogeny were snails able to fully reverse induced defenses. In daphnia (*Daphnia barbata*), reversibility of morphological defenses appears to be both trait and predator specific: Daphnia exposed to tadpoles (*Triops cancriformis*) but not to backswimmers (*Notonecta glauca*; an aquatic insect) show reversibility of defenses (Herzog et al., 2016). Notably, reversibility was higher for traits related to tail-spine (spina) length compared to helmets. Taken together these examples illustrate that reversibility comes in different flavors. Some traits are fully reversible while others are only partially reversible. In some cases reversibility occurs across an organism’s entire lifetime while it is limited to shorter developmental windows in others. Why does this variation exist? To date, there exist only a handful of theoretical studies exploring the evolution of reversible plasticity (Piersma & Drent, 2003).

### Lessons learned from modeling the evolution of reversible plasticity

The theoretical framework for understanding the evolution of reversible plasticity remains underdeveloped (Dupont et al., 2024; Piersma & Drent, 2003). The little work that exists mostly focuses on identifying the environmental conditions under which reversibility has a fitness advantage over irreversibility (Botero et al., 2015; Gabriel, 2005; Gabriel et al., 2005; Pfab et al., 2016; Ratikainen & Kokko, 2019; Utz et al., 2014). This research finds that reversible plasticity evolves when environmental conditions change across an organism’s life and the lifespan is long. Importantly, reversibility is only useful if organisms are able to predict changes based on environmental cues. Under such conditions, the benefits of phenotype reversal are thought to outweigh its costs (e.g. physiological costs or tradeoffs with specialization). In contrast, unpredictably changing environments should favor non-plastic strategies, such as conservative bet-hedging (Botero et al., 2015). In stable environments reversibility is assumed to provide no benefits and organisms are expected to evolve irreversible phenotypic plasticity.

Existing models typically assume that organisms can instantaneously develop fully specialized phenotypes; only one model assumes incremental phenotypic development (Pfab et al., 2016). To illustrate the difference, let us revisit the example of marine iguanas adjusting body size based on food availability (Wikelski & Thom, 2000). If body size adjustment happens instantaneously, iguanas should instantaneously achieve the target body size following changes in food availability. If body size adjustment is incremental, iguanas will develop through intermediate body sizes before reaching their target. The model suggests that reversibility only outperforms irreversibility when incrementally developing organisms are able to reach their target phenotypes (Pfab et al., 2016). This is possible when the environment changes smore slowly than it takes organisms to achieve the target phenotype matching their current environment. Here, incremental development constraints the benefits of reversibility.

### Gaps in our understanding of reversible plasticity

The existing body of theoretical studies does not yet fully capture the variety of reversible plasticity in nature. Notably, only one model (Pfab et al., 2016) has explored the evolution of reversibility when development is incremental. However, the fact that morphological traits, which tend to develop incrementally (e.g. defensive morphologies in daphnia), can be reversible highlights the need for more theory in this area. Previous work has illustrated that incremental development can constrain reversibility (Pfab et al., 2016). It is an open question how incremental development constraints the timing of plasticity in phenotype construction and deconstruction across ontogeny. Moreover, existing modeling does not address when we should expect ‘temporary’ instead of life-long reversibility. Or, when we should expect full instead of partial reversibility.

Here, we explore these questions within our modeling framework for studying the evolution and development of sensitive periods (Frankenhuis & Panchanathan, 2011; Panchanathan & Frankenhuis, 2016; Walasek et al., 2022a, 2022b). These models – assuming incremental development – explore when during ontogeny natural selection favors the highest levels of phenotypic plasticity (so-called sensitive periods; reviewed in Fawcett & Frankenhuis, 2015; Frankenhuis & Fraley, 2017; Frankenhuis & Walasek, 2020; Walasek et al., n.d.). We extend this previous work by allowing organisms to deconstruct phenotypic adjustments, resulting in the first model to separately examine plasticity in phenotype construction and deconstruction. In our model, the environment varies between generations but is stable within generations. Newborn organisms are uncertain about the state of their environment. Using Bayesian inference, organisms learn about the state of their environment throughout ontogeny. Across ontogeny, organisms repeatedly choose between phenotype construction and deconstruction. We compare two different modes of deconstruction: Organisms can either incrementally undo phenotypic adjustments or completely discard all of them in one go.

With this model we explore the following research questions. When do sensitive periods for construction and deconstruction evolve? What environmental conditions favor incremental or complete deconstruction? And, how common is reversibility when the environment is stable but organisms are uncertain about its state?

## Model

### Environment and organism

The environment consists of an infinite number of discrete, non-overlapping patches. Each patch can be in one of two states, *E*_0_(e.g. dangerous) or *E*_1_(e.g. safe). Patch states remain stable across generations. Organisms are born and randomly disperse into patches. Organisms do not initially know the state of their patch. However, organisms are equipped with an evolutionary prior distribution over the states, denoted by *P*(*E*_0_) and *E*(*E*_1_), indicating the initial probability of dispersing into either state (Stamps & Frankenhuis, 2016). In line with previous models (Frankenhuis & Panchanathan, 2011; Panchanathan & Frankenhuis, 2016; Walasek et al., 2022a, 2022b), we explore uncertain (0.5), moderately certain (0.7), and highly certain (0.9) priors.

Across ontogeny organisms sample cost-free cues (*C*_0_ or *C*_1_) to learn about the state of their patch. Ontogeny, during which organisms sample cues and make phenotypic decisions can last 5, 10, or 20 time periods. Information obtained from these cues is imperfect because cues do not perfectly predict the actual state of the patch. How well a cue predicts the environmental state depends on the cue reliability. The cue reliability, *P*(*C*_*x*_|*E*_*x*_), corresponds to the probability of observing a cue of a specific type (*x*) conditioned on being in the corresponding environment. The probability of observing the wrong cue then corresponds to *P*(*C*_*y*_|*E*_*x*_) = 1 − *P*(*C*_*x*_|*E*_*x*_). We assume symmetric cue reliabilities, such that *P*(*C*_0_|*E*_0_) = *P*(*C*_1_|*E*_1_). Based on a cue organisms update their current distribution over environmental states to a posterior distribution, *P*(*E*_0_|*C*_*x*_) and *P*(*E*_1_|*C*_*x*_), through Bayesian inference (Dall et al., 2015; McNamara et al., 2006; Mcnamara & Houston, 1980; Trimmer et al., 2011). In line with previous models, we explore low (0.55), moderate (0.75), and high cue reliabilities (0.95).

### Phenotypic development

Each environmental state, *E*_0_ and *E*_1_, is associated with a corresponding optimal phenotype, *P*_0_and *P*_1_. We assume that *P*_0_ and *P*_1_ are two different and independent traits. This means that specializations towards *P*_0_ are independent of specializations towards *P*_1_. For example, *P*_0_ could correspond to developing armor to fight predators and *P*_1_ to developing ornamentation for attracting mates. This model extends a previous model of sensitive period evolution (Panchanathan & Frankenhuis, 2016) by allowing organisms to deconstruct developed phenotypes. In each time period during ontogeny organisms can choose one of five options: (1) incrementally develop towards *P*_0_, (2) incrementally develop towards *P*_1_, (3) deconstruct previously developed phenotypic specializations towards *P*_0_, (4) deconstruct previously developed phenotypic specializations towards *P*_1_, or (5) wait and forgo phenotypic changes. Phenotypic construction and deconstruction is cost-free. However, the fact that ontogeny is constrained creates trade-offs between specialization and plasticity (i.e. adjusting phenotypes based on cues).

We developed two versions of the same model, assuming two different modes of phenotypic deconstruction. In the first version, we assume phenotypes can be *incrementally deconstructed*: During any specific time period, an organism can only deconstruct one phenotypic adjustment from one phenotypic target. In the second model, we assume *complete deconstruction*: During any time period, an organism can choose to discard all previously developed specializations from one phenotypic target. Thus, in both models, organisms can choose one of the following options in each time period: Specialize towards *P*_0_, specialize towards *P*_1_, wait and forgo phenotypic adjustment, deconstruct *P*_0_, and deconstruct *P*_1_. In each time period, organisms choose the phenotypic option that maximizes long-term expected fitness. However, in some cases multiple options may yield the same expected pay-off, resulting in ties. Organisms then choose randomly between tied options.

### Fitness

Organisms accrue fitness at the end of ontogeny based on how well their phenotype matches the actual state of their patch. An organism in *E*_0_ attains the maximum fitness if it is fully specialized towards *P*_0_. Likewise, an organism in *E*_1_ attains the maximum fitness if it is fully specialized towards *P*_1_. We assume that fitness at the end of ontogeny consists of three components: baseline fitness (*π*_0_), rewards for correct phenotypic specializations (*φ*), and penalties for incorrect phenotypic specializations (*ψ*). Thus, an organism who ends ontogeny with zero specializations towards either phenotypic target achieves baseline fitness. In principle, organisms can also achieve fitness below baseline if their fitness penalties are larger than their rewards.

We explore three different mappings between phenotypic adjustments and marginal (i.e. additional to baseline fitness) fitness rewards and penalties: linear, increasing, and diminishing (see ESM 1 for the formulas). The maximally attainable fitness is equivalent across all mappings. With diminishing rewards or penalties, a few specializations have large fitness consequences. With increasing rewards or penalties, many specializations are needed to achieve large fitness consequences. We assume equal weights for rewards from correct specializations and penalties from incorrect specializations.

### Optimal policies

For each combination of prior, cue reliability, and reward-penalty mapping, we compute optimal policies. For every possible state across ontogeny (i.e. sampled cues and current phenotype), the optimal policy provides the phenotypic choice that maximizes expected fitness at the end of ontogeny. We use stochastic dynamic programming with backwards induction, to compute optimal policies (Mangel & Clark, 1988; McNamara & Houston, 1980; see ESM 1 for details).

### Analyses based on the optimal policies

Based on the optimal policies, we simulate populations of organisms from different evolutionary ecologies. Following their developmental trajectories across ontogeny, we can quantify changes in plasticity, compare fitness of different policies, and describe distributions of mature phenotypes.

### Quantifying plasticity

We use the optimal policies to simulate adoption experiments, resembling empirical study paradigms for quantifying plasticity. We simulate an organism that receives cues and develops according to the optimal policy. At some time period *E* during ontogeny, we clone that organism, creating an identical copy with the same phenotypic state and cues sampled. Now, we separate original and clone. From that point onwards the clone will receive opposite cues from the original. Thus, whenever the original samples *C*_0_ the clone samples *C*_1_ and vice versa. We then measure plasticity as the phenotypic distance between original and clone at the end of ontogeny. Large phenotypic distances between original and clone indicate high levels of plasticity. We simulate 10.000 pairs to account for the stochasticity in sequences of cues. Plasticity then corresponds to the average phenotypic distance across these 10.000 pairs of clones. We perform such an adoption experiment for each time period in ontogeny. Thus, plasticity at time period 5 corresponds to the average phenotypic distance across 10.000 simulated pairs of clones separated in that time period.

For each pair of clones, we compute three phenotypic ‘distance’ measures: total phenotypic distance (‘total plasticity’), distance in construction (‘plasticity in construction’), and distance in deconstruction (‘plasticity in deconstruction’). Our separate distance measures provide insights into when during ontogeny cues have the largest impact on construction and deconstruction and how this plasticity shapes total phenotypic development.

Total phenotypic distance corresponds to the Euclidean distance between the number of specializations towards either phenotypic target. Distance in construction corresponds to the Euclidean distance between the number of time periods spent constructing the phenotypic targets. Likewise, distance in deconstruction corresponds to the Euclidean distance between the number of time periods spent deconstructing. We normalize all phenotypic distance measures to range between 0 and 1 by dividing each measure by the maximally attainable distance. Total phenotypic distance closely maps onto empirical measures of plasticity. Empiricists routinely measure plasticity as the phenotypic difference between individuals in a control and treatment group at the end of the observation period (Stamps & Luttbeg, 2022). We can think of this plasticity as the consequence of plasticity in construction and deconstruction. To quantify plasticity in construction or deconstruction alone, empiricists would need to continuously monitor individuals across ontogeny.

We additionally explore alternative paradigms for quantifying plasticity and their results. In our ‘base paradigm’, as described above, separated clones experience opposite cues until the end of ontogeny. Plasticity corresponds to the phenotypic distance at the end of ontogeny. In line with previous work (Walasek et al., 2022a), we vary this base paradigm along three dimensions: treatment, exposure duration, and measurement time. In the base paradigm the clone receives reciprocal, opposite cues from the original. We explore two less extreme treatments in which the clone receives cues representative of the opposite patch or uninformative cues (a form of informational deprivation). In the base paradigm, original and clone remain separated until the end of ontogeny. We additionally explore a scenario where clones are separated temporarily. Lastly, in the base paradigm we always measure phenotypic distances at the end of ontogeny. With temporary separation, we can either measure phenotypic distance right after the exposure period or at the end of ontogeny, after original and clone have been reunited for some time. Different combinations of treatment, exposure duration, and measurement time cover a broader range of empirical study paradigms.

### Fitness differences and distributions of mature phenotypes

Contrasting (typically plastic) optimal policies—capable of incremental or complete deconstruction—with a (typically plastic) policy without deconstruction and non-plastic strategies reveals the extent to which deconstruction provides a fitness advantage in different environments. To this end, we compare the fitness of the optimal policy (allowing for deconstruction) to the fitness attained by a policy without deconstruction, and two non-plastic strategies. For each combination of cue reliability, prior, and reward/penalty mapping, we compute fitness of an optimal policy derived from a model with incremental, irreversible construction (Panchanathan & Frankenhuis, 2016) and fitness of a pure generalist and a pure specialist strategy. Pure generalists always specialize halfway towards either phenotypic target. Pure specialists fully specialize towards the phenotypic target that matches the most likely environmental state according to the prior distribution. In case of uncertain priors (0.5), half the population fully specializes towards *P*_0_ and the other half towards *P*_1_.

Lastly, we use the optimal policies to simulate distributions of mature phenotypes. Based on these distributions we can quantify the extent to which deconstruction evolved for any combination of prior, cue reliability, and reward/penalty mapping. Moreover, the distribution of mature phenotypes allows us to see whether a population of organisms tends to develop more specialized or generalist phenotypes and how much organisms tend to deconstruct (partial vs full) depending on environmental conditions. Our code is written in Python 3.10 and available on GitHub (https://github.com/anonymousAccountForPublication/reversible_plasticity).

## Results

We focus on two main research questions. First, under what environmental conditions does reversibility modeled as deconstruction evolve? Second, when during ontogeny do cues have the largest impact on construction and deconstruction? In answering each question, we highlight differences between incremental and complete deconstruction.

### Under what environmental conditions does reversibility evolve?

Incremental and complete deconstruction evolve frequently in different evolutionary ecologies (Figures 1 and 2). The prevalence of deconstruction tends to decrease as priors become more certain (further away from 0.5). When priors are uncertain (0.5), deconstruction decreases with increasing cue reliability. More certain priors and higher cue reliabilities reduce organisms’ uncertainty about their environment, allowing them to invest more heavily into construction compared to deconstruction. When priors are informative (0.7 and 0.9), deconstruction can be highest with moderate cue reliability (0.75). Here, cues are sufficiently informative for organisms to invest into specialization. At the same time, these cues leave room for uncertainty, leading organisms to revise their estimates throughout ontogeny and potentially deconstruct specializations. We observe little deconstruction with increasing penalties (ESM 2). Organisms focus on specialization because a few incorrect adjustments carry relatively small fitness costs.

**Figure 1:**
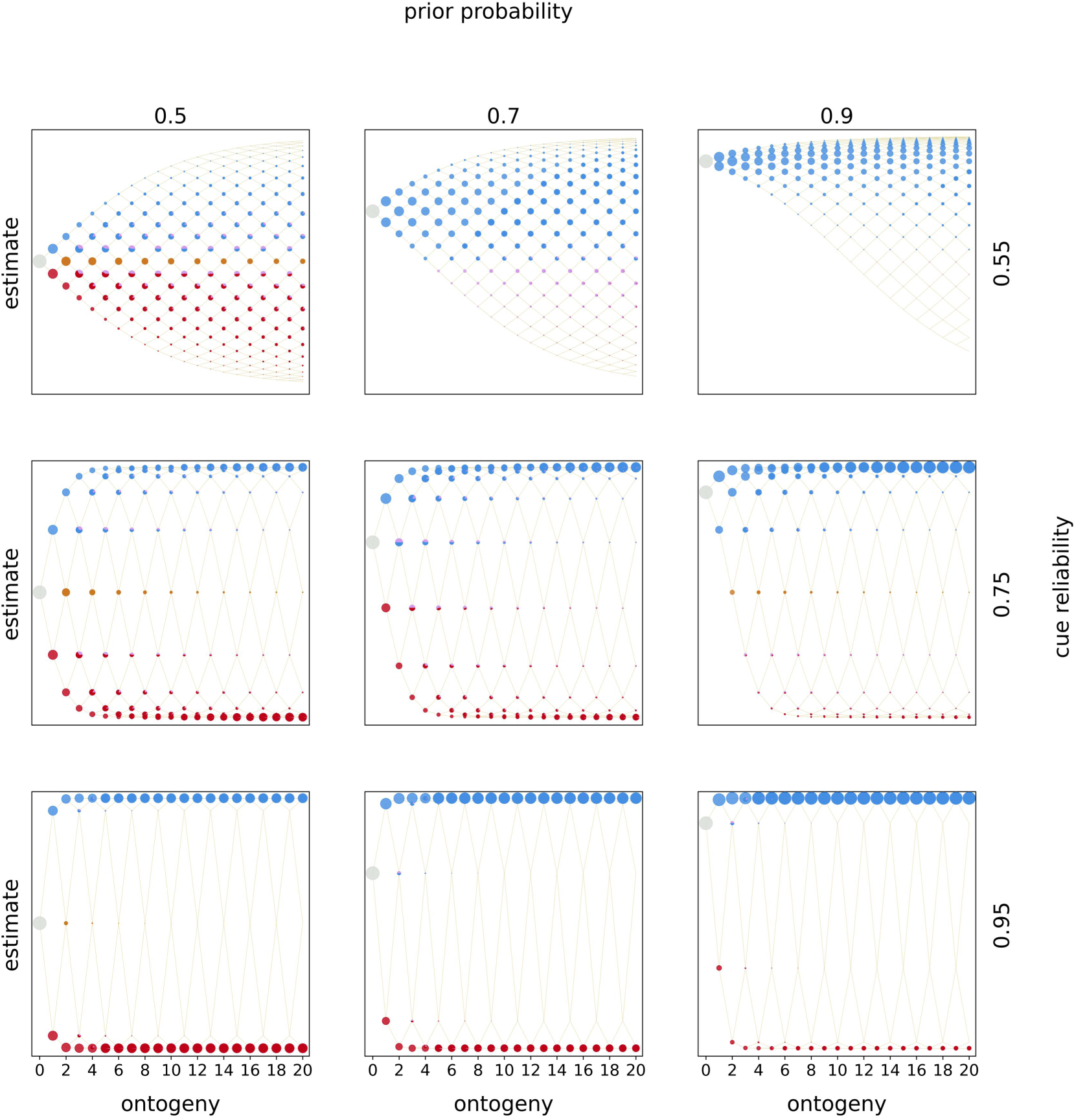
Optimal policies. Optimal policies are shown for a model with incremental deconstruction and linear rewards and penalties. Rows indicate the prior estimate of being in *E*_1_and columns indicate the cue reliability. Within each panel, the horizontal axis denotes ontogeny and the vertical axis the posterior estimates of being in *E*_1_. The entire population starts ontogeny with zero cues sampled and the prior estimate indicated by the row (indicated by the grey circle). In each time period organisms sample a cue (either *C*_0_or *C*_1_), update their estimate, and make a phenotypic decision (colored circles). Beige lines indicate developmental trajectories through this decision space, with lines branching upwards indicating the sampling of *C*_1_ and lines branching downwards indicating the sampling of *C*_*0*_. Colors denote the optimal, fitness-maximizing phenotypic choice in each state. Pies indicate cases in which organisms with the same posterior estimates make different phenotypic decisions. The area of a circle (pie piece) is proportional the probability of reaching that particular state. Colors indicate the following phenotypic decisions: Black corresponds to waiting, red to constructing *P*_0_and light red to deconstructing *P*_1_, blue to constructing *P*_1_and light blue to deconstructing *P*_0_, purple to ties without deconstruction and light purple to ties with deconstruction, and lastly ochre corresponds to a tie between all options. We show optimal policies for all other reward-penalty mappings in the ESM 2 (Figures S2.1-S2.9 for incremental deconstruction and Figures S2.10-S2.18 for complete deconstruction). In the ESM same optimal policies are shown with a more fine grained distinction between different types of ties.

**Figure 2:**
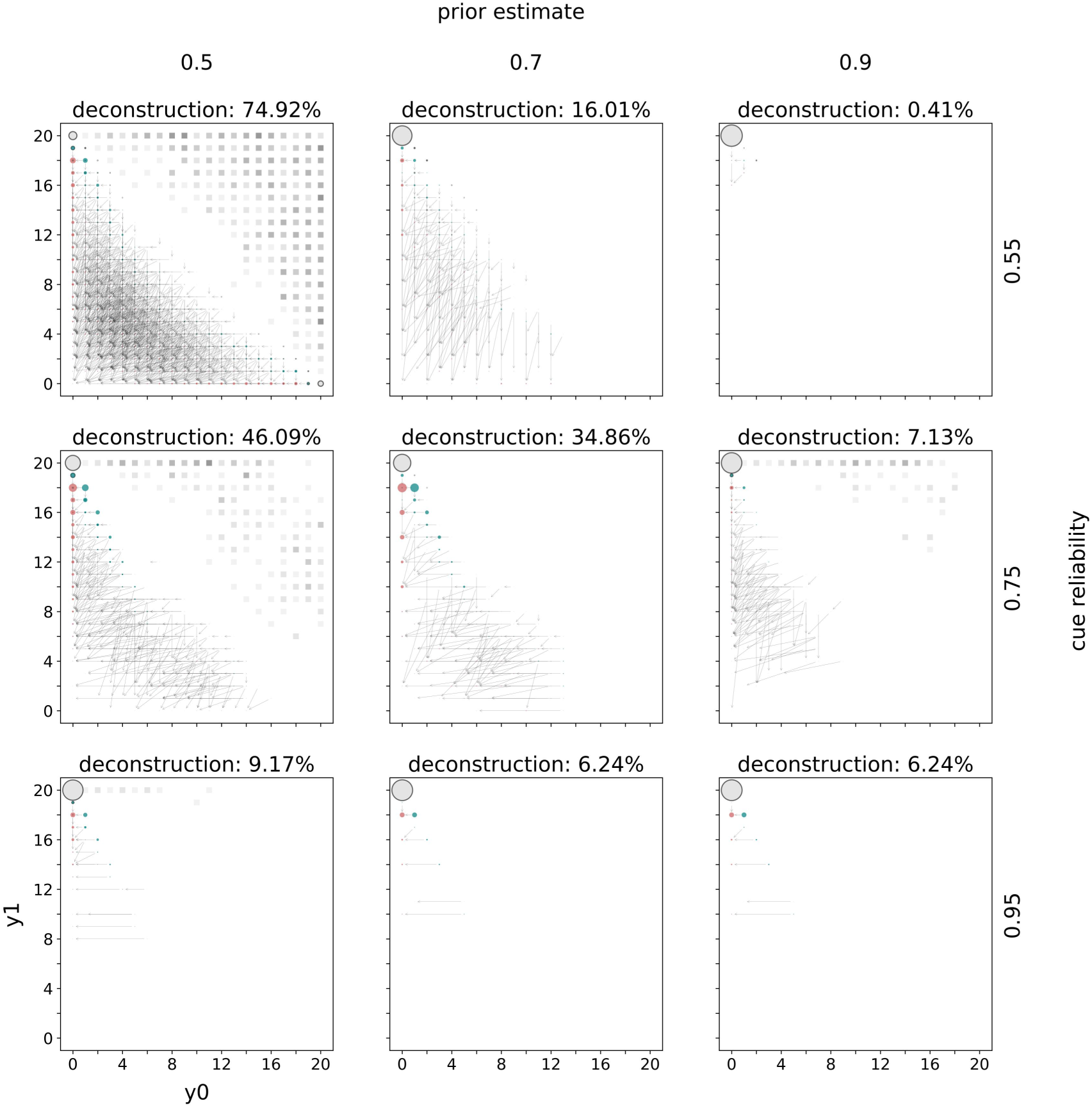
Distributions of mature phenotypes. Distributions of mature phenotypes are shown for a model with incremental deconstruction and linear rewards and penalties. Rows indicate the prior estimate of being in *E*_1_and columns indicate the cue reliability. The populations of mature phenotypes have been simulated in *E*_1_. The title of each panel indicates the percentage of mature phenotypes that have deconstructed at some point during ontogeny. Within each panel the horizontal axis indicates the number of specializations towards *E*_0_ and the vertical axis towards *E*_1_. The lower triangle indicates how much mature phenotypes have constructed (teal circles) and what their phenotype looked like after deconstruction (red circles). Grey arrows connect phenotypes before (teal) and after (red) deconstruction. Grey circles with a black outline belong to mature phenotypes that never deconstructed. The area of a circle is proportional to the number mature organisms with this phenotype. The upper triangle indicates waiting. For each mature phenotype (after deconstruction) below the diagonal the corresponding square above the diagonal highlights the amount of waiting. The color intensity is proportional to the amount of waiting. Black squares indicate phenotypes that waited all of ontogeny (i.e. 20 time periods) and white squares phenotypes that never waited. We show distributions of mature phenotypes for all other reward-penalty mappings in the ESM 2 (Figures S2.19-S2.27 for incremental deconstruction and Figures S2.28-S2.36 for complete deconstruction).

How organisms deconstruct depends on environmental conditions. With increasingly higher cue reliabilities, organisms tend to only deconstruct incorrect phenotypic specializations (across rows in Figure 2 and ESM 2 Figure S2.28). With lower cue reliabilities, organisms deconstruct in either direction because cues are less informative about the actual environment.

In some cases, deconstruction favors mature organisms with zero phenotypic specializations. These mature phenotypes result from a mix of deconstruction and waiting. They are favored when few incorrect specializations lead to high fitness costs and few correct specializations only yield small rewards (linear/diminishing, increasing/diminishing, linear/diminishing reward/penalties; ESM 2). Thus, organisms mature with zero specializations to avoid penalties from incorrect specializations. The proportion of phenotypes with zero specializations is largest for uncertain priors (0.5) and low cue reliability (0.55). Under these conditions, organisms sometimes even choose to deconstruct a phenotype that only has correct specializations. This pattern is driven by those organisms who remain highly uncertain about the state of their environment until the end of ontogeny.

Compared to other plastic and non-plastic strategies deconstruction only conveys a small fitness benefit (Figure 3). With linear rewards and penalties, a model without deconstruction performs as well as a model with incremental deconstruction. Complete deconstruction conveys a small fitness benefit over these strategies. This benefit is largest, when cues are not highly reliable (0.55 or 0.75) and priors are not highly certain (0.5 or 0.7). These conditions produce a small proportion of organisms who are ‘moderately’ certain about the true environmental state towards the end of ontogeny (Figure 1). When deconstruction is complete, these organisms are able to discard all specializations for the other environmental state (ESM 2 Figure S2.28). In contrast, incremental deconstruction is gradual and time-consuming, such that some organisms will only be able to partially deconstruct these specializations (Figure 2). Under some conditions both incremental and complete deconstruction outperform a policy without deconstruction. This happens when penalties for a few incorrect specializations are higher than rewards for a few correct specializations (linear/diminishing, increasing/diminishing, and increasing/linear reward/penalties; ESM 2).

**Figure 3:**
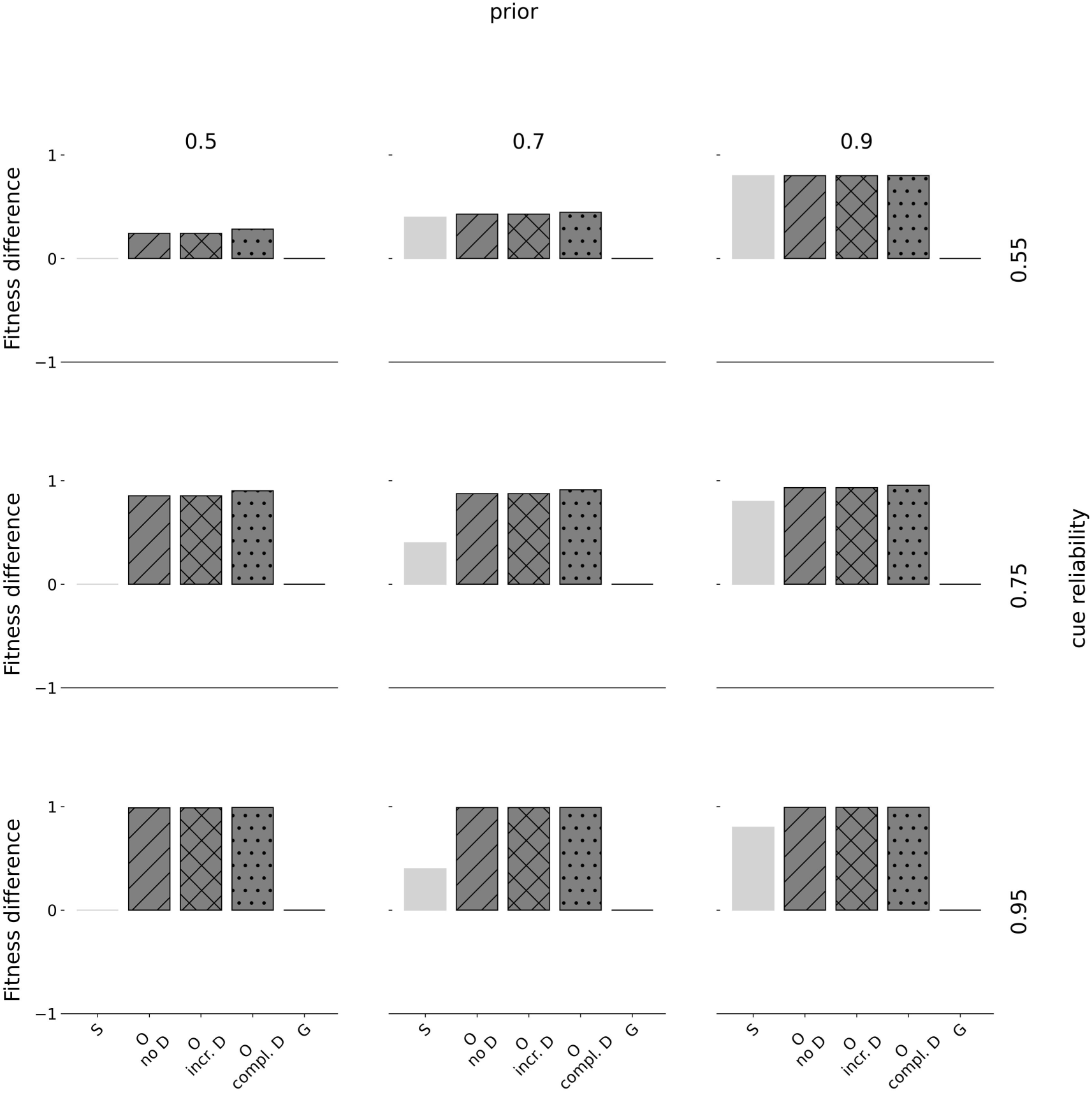
Fitness differences. Fitness differences are shown for linear rewards and penalties. Rows indicate the prior estimate of being in *E*_1_and columns indicate the cue reliability. Within each panel, the horizontal axis denotes the type of strategy where ‘S’ corresponds to a pure specialist strategy, ‘O’ to an optimal policy, and ‘G’ to a pure generalist strategy. The horizontal axis denotes fitness differences from baseline (corresponding to 0), normalized to range between −1 and 1. We show fitness of three different optimal policies: without deconstruction (‘no D’), with incremental deconstruction (‘incr. D’), and complete deconstruction (‘complete D’). Specialists always fully specialize according to the prior distribution. When priors are uncertain (0.5), half the population fully specializes towards *P*_0_ and the other one towards *P*_1_. Generalists always specialize halfway towards either phenotypic target. We show fitness differences for all other reward-penalty mappings in the ESM 2 Figures S2.37-S2.45.

Here, the relatively harsh penalties induce a selective advantage for deconstruction. Notably, policies with complete deconstruction still outperform those with incremental deconstruction. Lastly, we explored to what extent the evolution of deconstruction depends on the duration of ontogeny. When cues a are low in reliability (0.55), the amount of deconstruction (both incremental and complete) is qualitatively similar across all durations of ontogeny (ESM 3). However, when cues are moderately (0.75) or highly (0.95) reliable, shorter durations of ontogeny (5 or 10 time periods) favor larger and more constant amounts of deconstruction across priors compared to a long duration of ontogeny (20 time periods). When there is less time to sample cues, a larger proportion of organisms remains uncertain across ontogeny despite more reliable cues. These organisms are more likely to deconstruct.

### Timing of sensitive periods: When during ontogeny do cues have the largest impact

#### When during ontogeny do cues have the largest impact on construction

Our results indicate early-ontogeny sensitive periods for construction (Figure 4). Plasticity in construction is highest at the onset of ontogeny and declines gradually across ontogeny. The only exception occurs with highly certain priors (0.9) and low cue reliability (0.55): These conditions select against any plasticity; organisms ignore uninformative cues and fully specialize based on their prior. Interestingly, patterns of plasticity in construction (dotted lines with stars) are highly similar with incremental (teal lines) and complete deconstruction (yellow lines). Moreover, these patterns of plasticity in construction resemble those resulting from a model without deconstruction (Panchanathan & Frankenhuis, 2016). These results suggest an independence between the evolution of sensitive periods for construction and deconstruction.

**Figure 4:**
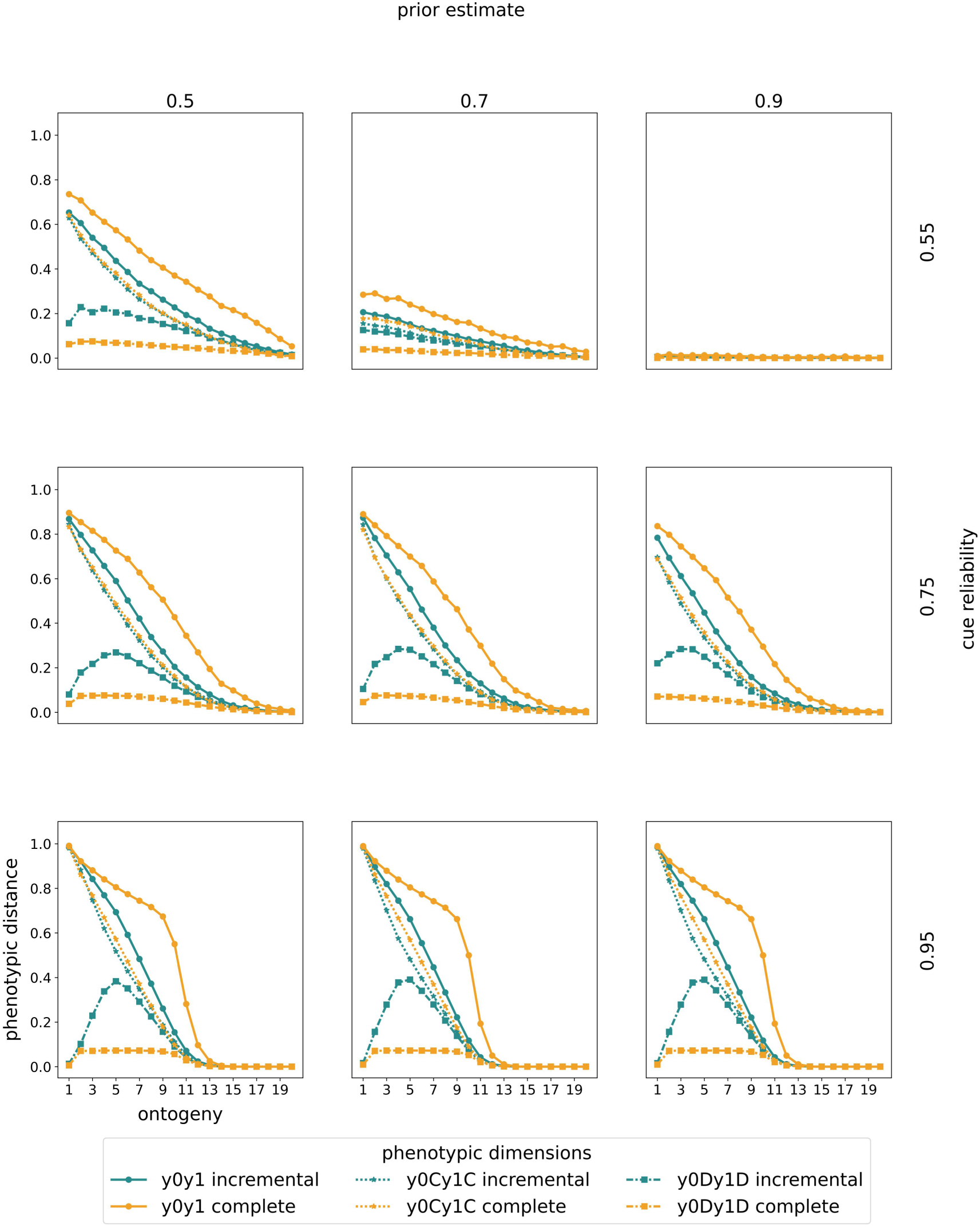
Changes in plasticity. Changes in plasticity are shown for linear rewards and penalties. Rows indicate the prior estimate of being in *E*_1_and columns indicate the cue reliability. Within each panel, the horizontal axis denotes ontogeny. The vertical axis denotes the normalized average phenotypic distance (our plasticity measure) across 10.000 pairs of simulated clones measured at the end of ontogeny. A specific point on any of the curves corresponds to a simulation experiment in which clones have been separated at the time point indicated by the horizontal axis. We show results from a model with incremental deconstruction in teal and with complete deconstruction in yellow. For each mode of deconstruction, we present three types of plasticity curves. First, we show plasticity in construction (dotted lines and stars) as the Euclidean distance between the number of time points spent constructing. Second, we show plasticity in deconstruction (dash-dotted lines and squares) as the Euclidean distance between the number of time points spent deconstruction. Third, we show total plasticity (solid lines and circles) as the Euclidean distance between the number of specialization steps towards either target (after accounting for deconstruction). We normalize phenotypic distance in construction and total phenotypes by dividing by the maximally possible Euclidean distance, corresponding to 2 ∗ √*T* = 20. The normalization constant for deconstruction is 2 ∗ √*T*/2 = 10. We show changes in plasticity for all other reward-penalty mappings in ESM 2, Figures S2.46-S2.54.

#### When during ontogeny do cues have the largest impact on deconstruction

Our results indicate mid-ontogeny sensitive periods for deconstruction when cues are informative (Figure 4). With moderately or highly reliable cues (0.75 or 0.95), plasticity in deconstruction (dash-dotted lines and squares) first increases before decreasing: Organisms show the largest differences in deconstruction when they are separated after they have begun specializing towards one environmental state. The higher the cue reliability, the more organisms invest into specialization early in ontogeny, resulting in higher peaks in plasticity in deconstruction. In contrast to incremental deconstruction, complete deconstruction favors shallow peaks in plasticity. With complete deconstruction, small differences in the number of time steps spent deconstructing can have drastic consequences for phenotypic development. As a consequence, patterns of total plasticity (yellow solid lines and circles) deviate notably from plasticity in construction when deconstruction is complete. Overall, these results are qualitatively similar across all other reward/penalty mappings (ESM 2, Figures S2.46-S2.54).

We observe early-ontogeny sensitive periods in deconstruction when cues are low in reliability (0.55; with the exception of highly certain priors). Here, the window of plasticity in deconstruction covers a wider range of ontogeny compared to the windows emerging from higher cue reliabilities. With low cue reliabilities, a larger proportion of the population is uncertain about their environment (Figure 1). These organisms tends to accrue specializations towards both phenotypic targets across all of ontogeny. When some of these organisms achieve more certain posteriors throughout ontogeny, they will deconstruct specializations towards the less likely state. The earlier organisms are separated, the more they deconstruct. Patterns of sensitive periods in construction and deconstruction are qualitatively similar across different durations of ontogeny (5, 10, and 20 time periods; ESM 3).

#### Different paradigms for quantifying plasticity

Lastly, we explored the effects of different treatments, separation durations, and measurement times on patterns of plasticity (ESM 4). Manipulating the extent to which cues between separated organisms differ (‘treatment’) does not change results qualitatively (ESM 4, Figures S4.1-S4.3). We also compared plasticity following temporary separation measured right after separation and at the end of ontogeny (ESM 4, Figures S4.4-S4.5). Phenotypic differences following temporary exposure are often not visible right after the separation but only at the end of ontogeny. This implies a delay between organisms experiencing diverging cues and resulting phenotypic differences. This pattern applies especially to deconstruction: Organisms who are temporarily exposed to diverging cues, do not alter phenotypes during that time. However, these organisms deconstruct specializations made during this time before reaching adulthood. Here, the effect of cues on phenotype development is often largest towards the end of ontogeny. Taken together, this shows that we may not be able to observe effects of temporary separation without long-term measurements of phenotypic differences.

## Discussion

Our model studies the evolution of reversible phenotypic plasticity when organisms develop incrementally in a stable environment. During ontogeny organisms sample cues to the environmental state and adjust their phenotypes accordingly. We operationalized reversibility as the ability to deconstruct phenotypic adjustments, assuming two modes of deconstruction: Organisms can either deconstruct one phenotypic adjustment at a time (‘incremental’) or completely discard all adjustments of one type (‘complete’). In the discussion, we highlight four insights from our model, as well as limitations and future directions.

### Deconstruction can evolve in stable environments

Previous theory argues that reversibility evolves in the context of changing environments (Botero et al., 2015; Gabriel, 2005; Gabriel et al., 2005; Pfab et al., 2016; Ratikainen & Kokko, 2019; Utz et al., 2014). Our model shows that reversibility – modeled as deconstruction – evolves even in stable environments when organisms are adapted to changing environments across but not within generations. Our findings highlight the capacity for reversibility in species who experience environmental changes for the first time in their lives. Existing experimental studies of reversible plasticity typically expose animals to changing environmental conditions across development (e.g. Hoverman & Relyea, 2007).

Extending such experiments across generations would allow us to test under what conditions animals can evolve the capacity for reversibility. Such experimental evolution studies could manipulate whether organisms experience environmental change only across or also within generations. These studies would provide insight into whether species can develop reversible plasticity to combat environmental changes previously absent within generations.

### Shorter ontogeny increases the frequency of deconstruction

Our model shows that shorter durations of ontogeny select for reversibility across a wider range of environmental conditions. All else equal, shorter durations of ontogeny imply shorter lifespans of organisms. Traditionally, reversibility was thought to primarily benefit long-lived species and evolve alongside longevity (Hoffmann & Bridle, 2021; Ratikainen & Kokko, 2019). Notably, this work assumes that organisms can make phenotypic adjustments throughout the lifespan. Our model suggests that the ability to deconstruct may not be limited to long-lived organisms, when organisms can only make phenotypic adjustments during a fixed window of ontogeny. Reversible anti-predatory defenses in daphnia and snails fit this finding (Herzog et al., 2016; Hoverman & Relyea, 2007). Although typically only found in long-lived vertebrates, one study even observed reversible brain plasticity in insects: Sexually reproducing ants (*Harpegnathos saltator*) demonstrate reversible changes in brain volume when switching between forager and reproductive worker phenotypes in adulthood (Penick et al., 2021). These examples highlight that reversibility may evolve commonly across species with different life-histories.

### The fitness advantage of deconstruction

Complete deconstruction provides a larger fitness benefit compared to incremental deconstruction. This finding is line with previous models of reversible plasticity (Gabriel, 2005; Gabriel et al., 2005; Pfab et al., 2016; Utz et al., 2014). Those models find that the fitness benefit of reversible plasticity is largest with short response lags, which correspond to the time between inferring an environmental change and reversing a phenotype. Although we did not model response lags as such, our two modes of deconstruction are analogous, representing two extremes. When deconstruction is incremental reversing phenotypic adjustments takes time, whereas reversal is immediate with complete deconstruction.

Incremental deconstruction can thus be seen as a less efficient form of reversibility; it makes miscalibration more costly as it takes longer to undo wrong specializations and often only results in partial reversibility. It seems likely that incremental deconstruction should only evolve when organisms are constrained in their physiological development. Empirical patterns appear to align with results from models. We previously discussed a study in daphnia (*Daphnia barbata*) which showed higher reversibility for adjusting tail-spine (spina) length compared to helmets (Herzog et al., 2016). The lower reversibility of helmets is assumed to be the result of higher physiological constraints on adjusting helmets compared to spina (Utz et al., 2014). The fact that reversibility is more commonly observed in easily adjustable behavioral traits compared to morphological ones is also in line with this pattern (Burggren, 2020).

The cost of reversibility is not the only factor shaping variation in reversible plasticity: *Daphnia barbata* also showed differences in reversibility in the same trait (Herzog et al., 2016). Daphnia exposed to tadpoles (*Triops cancriformis*) but not to backswimmers (*Notonecta glauca*; an aquatic insect) show reversibility of defenses. Costs of deconstruction cannot explain these differences, as the same trait is being deconstructed in both cases. Based on our modeling results, such predator-specific differences in reversibility could be related to the relative fitness penalties of incorrect specializations. We find that deconstruction provides the largest fitness benefits in harsh environments in which only a few incorrect phenotypic adjustments result in large fitness costs. Thus, we may expect differences in reversibility when incorrect adjustments in the presence of one type of predator result in larger fitness penalties compared to another type. The authors of the daphnia study do not provide information about the relative fitness costs of incorrect specializations in the presence of either predator. However, they note that the removal of predator kairomones – a chemical substance emitted by organisms – is less informative in the case of an aquatic insect with the ability to fly (*Notonecta glauca*), compared to a water-bound tadpole (*Triops cancriformis*). The absence of kairomones does not guarantee the absence of flying insect predators. The authors suggest that the difference in cue reliabilities may explain why reversibility is higher in response to the removal of tadpole kairomones. Our model shows that reversibility increases with cue reliability. This explanation does not, however, exclude the possibility that organisms may exhibit increased levels of reversibility in response to the more costly predator. Future work could test this hypothesis in daphnia or some other short-lived species using experimental evolution techniques (English & Barreaux, 2020).

### Patterns of sensitive periods in construction and deconstruction

Our model shows that sensitive periods for deconstruction typically peak during the middle of ontogeny. When cues are informative, plasticity increases at first and then decreases – eventually disappearing, a process resulting in a critical period (Knudsen, 2004). During later stages of ontogeny, organisms have likely already specialized to some degree.

When these organisms receive informative cues indicating a different environment, they should focus on deconstructing prior specializations. When cues are less informative, plasticity in deconstruction tends to be higher at the onset of ontogeny, though the window for plasticity is wider, sometimes spanning most of ontogeny. These results align with existing findings about the evolution of reversible and irreversible plasticity. Reversible plasticity, traditionally defined as the life-long ability to adjust phenotypes, should be favored in changing environments (Botero et al., 2015; McDermott & Safran, 2021; Ratikainen & Kokko, 2019). Changing environments create uncertainty about the state of the world. In our model, the window during which organisms can deconstruct phenotypic adjustments remains wide open when organisms are uncertain about their environment. Irreversible plasticity, commonly defined as the ability to make phenotypic adjustments during a delineated time window in ontogeny should be favored in stable environments (Dupont et al., 2024).

Compared to changing environmental conditions, stable environments are easier to infer, resulting in reduced uncertainty. In our model, delineated temporal windows of reversibility result when organisms can reduce uncertainty about the environment state. Our model shows that the distinction between reversible and irreversible plasticity does not require environmental change. Instead, the distinction between reversible and irreversible plasticity depends on an organism’s uncertainty about the environmental state.

There exist other models of sensitive period evolution in which organisms can reverse phenotypes across their entire lifespan (Fischer et al., 2014; Stamps & Krishnan, 2014, 2017). These models assume instantaneous development, such that organisms can express any phenotype at any time. Across sensitive period models that do (Fischer et al., 2014; Stamps & Krishnan, 2014, 2017) and do not (English et al., 2016; Frankenhuis & Panchanathan, 2011; Panchanathan & Frankenhuis, 2016; Walasek et al., 2022a, 2022b) allow for reversibility, patterns of sensitive periods are remarkably robust. In those models, as in our model, plasticity tends to be highest early in ontogeny before continuously declining. The robustness of this result across a range of models making different assumptions suggests high selection pressures on the evolution of early-life sensitive periods across traits and species. However, understanding whether these patterns result purely from construction or from a combination of construction and deconstruction provides insights about organisms’ development.

### Limitations and future directions

While our model provides novel insights into the evolution of reversible plasticity in stable environments, we see value in exploring fluctuating environments within our modeling framework. Fluctuating environments are often viewed as a requirement for the evolution of reversible plasticity. All else equal, we expect larger fitness benefits of reversibility when environmental conditions can change. A previous model of sensitive period evolution has found that rapidly changing environmental conditions favor sensitive periods towards the end of ontogeny (Walasek et al., 2022b). The reason is that organisms rely on the most recent information to predict their adult environment. However, if organisms were able to deconstruct phenotypic adjustments, plasticity in construction could be higher earlier in ontogeny and organisms might instead use late-ontogeny cues for deconstruction. As a result, patterns of sensitive periods in fluctuating environments may be less robust across models with and without deconstruction. This would imply an interdependence between the evolution of sensitive periods for construction and deconstruction in changing environments.

Studying reversibility across ontogeny in changing environments can also help us make sense of empirical phenomena. For example, there are humans born with severely impaired vision (congenital cataracts) which can be restored later in life (Bruns & Röder, 2023; Sourav et al., 2024). Early in life these individuals develop compensatory abilities in other sensory systems (e.g. hearing). It is an open question under what conditions human brains retain or ‘deconstruct’ these specializations after vision is restored. Theoretical modeling may help us understand under what environmental conditions natural selection favors the deconstruction of initial specializations after an environmental change.

Future models exploring changing environments could be further extended by allowing organisms to adjust phenotypes throughout their entire lifetimes. By varying lifespan, modelers can then explore to what extent patterns of reversibility depend on longevity when organisms develop incrementally. Previous modeling work, assuming instantaneous development, suggests that longevity and reversibility co-evolve in changing environments (Ratikainen & Kokko, 2019). It is an open question whether we should expect the same result when organisms develop incrementally.

Research on reversible plasticity is lacking both empirically and theoretically. Reversibility is difficult to study empirically. To identify the existence or absence of reversibility, an organism would ideally be monitored throughout its entire life (Burggren, 2020; Stamps & Luttbeg, 2022). Theory outlining the conditions which favor reversibility can help focus empirical research efforts on particular species, traits, or developmental stages.

Our model provides novel insights into when we may expect reversibility in organisms adapted to stable environments. Our model also highlights the benefits of widening the definition of reversibility (i.e. life-long and fully reversible plasticity) to also include temporary and partial reversibility. As noted by others (Burggren, 2020; Herzog et al., 2016), adopting a broader definition of reversibility can capture interesting variation across species and traits. With it we can move towards a theoretical framework that can explain the full breadth of reversibility in nature.

## Supporting information

ESM

ESM

## Notes

**Funding:** This work used the Dutch national e-infrastructure with the support of the SURF Cooperative using grant no. EINF-6641. WEF’s contributions have been supported by the Dutch Research Council (V1. Vidi.195.130) and the James S. McDonnell Foundation (https://doi.org/10.37717/220020502).

### Competing Interest Statement

The authors have declared no competing interest.

## References

Botero, C. A., Weissing, F. J., Wright, J., & Rubenstein, D. R. (2015). Evolutionary tipping points in the capacity to adapt to environmental change. Proceedings of the National Academy of Sciences of the United States of America, 112(1), 184–189. 10.1073/pnas.1408589111

Bruns, P., & Röder, B. (2023). Development and experience-dependence of multisensory spatial processing. Trends in Cognitive Sciences, S136466132300102X. 10.1016/j.tics.2023.04.012

Burggren, W. W. (2020). Phenotypic switching resulting from developmental plasticity: fixed or reversible? Frontiers in Physiology, 10(January), 1–13. 10.3389/fphys.2019.01634

Dall, S. R. X., McNamara, J. M., & Leimar, O. (2015). Genes as cues: Phenotypic integration of genetic and epigenetic information from a Darwinian perspective. Trends in Ecology and Evolution, 30(6), 327–333. 10.1016/j.tree.2015.04.002

Dupont, L., Thierry, M., Zinger, L., Legrand, D., & Jacob, S. (2024). Beyond reaction norms: The temporal dynamics of phenotypic plasticity. Trends in Ecology & Evolution, 39(1), 41–51. 10.1016/j.tree.2023.08.014

English, S., & Barreaux, A. M. (2020). The evolution of sensitive periods in development: Insights from insects. Current Opinion in Behavioral Sciences, 36, 71–78. 10.1016/j.cobeha.2020.07.009

English, S., Fawcett, T. W., Higginson, A. D., Trimmer, P. C., & Uller, T. (2016). Adaptive use of information during growth can explain long-term effects of early life experiences. American Naturalist, 187(5), 620–632. 10.1086/685644

Fawcett, T. W., & Frankenhuis, W. E. (2015). Adaptive explanations for sensitive windows in development. Frontiers in Zoology, 12(Suppl 1), S3. 10.1186/1742-9994-12-S1-S3

Fischer, B., van Doorn, G. S., Dieckmann, U., & Taborsky, B. (2014). The evolution of age-dependent plasticity. The American Naturalist, 183(1), 108–125. 10.1086/674008

Frankenhuis, W. E., & Fraley, R. C. (2017). What do evolutionary models teach us about sensitive periods in psychological development? European Psychologist, 22(3), 141–150. 10.1027/1016-9040/a000265

Frankenhuis, W. E., & Panchanathan, K. (2011). Balancing sampling and specialization: An adaptationist model of incremental development. Proceedings of the Royal Society B: Biological Sciences, 278(1724), 3558–3565. 10.1098/rspb.2011.0055

Frankenhuis, W. E., & Walasek, N. (2020). Modeling the evolution of sensitive periods. Developmental Cognitive Neuroscience, 41, 100715. 10.1016/j.dcn.2019.100715

Gabriel, W. (2005). How stress selects for reversible phenotypic plasticity. Journal of Evolutionary Biology, 18(4), 873–883. 10.1111/j.1420-9101.2005.00959.x

Gabriel, W., Luttbeg, B., Sih, A., & Tollrian, R. (2005). Environmental tolerance, heterogeneity, and the evolution of reversible plastic responses. The American Naturalist, 166(3), 339–353. 10.1086/432558

Herzog, Q., Tittgen, C., & Laforsch, C. (2016). Predator-specific reversibility of morphological defenses in Daphnia barbata. Journal of Plankton Research, 38(4), 771–780. 10.1093/plankt/fbw045

Hoffmann, A. A., & Bridle, J. (2021). The dangers of irreversibility in an age of increased uncertainty: Revisiting plasticity in invertebrates. Oikos. 10.1111/oik.08715

Hoverman, J. T., & Relyea, R. A. (2007). How flexible is phenotypic plasticity? Developmental windows for trait induction and reversal. Ecology, 88(3), 693–705. 10.1890/05-1697

Knudsen, E. I. (2004). Sensitive periods in the development of the brain and behavior. Journal of Cognitive Neuroscience, 16(8), 1412–1425. 10.1162/0898929042304796

Lorenzi, V., Earley, R. L., & Grober, M. S. (2006). Preventing behavioural interactions with a male facilitates sex change in female bluebanded gobies, Lythrypnus dalli. Behavioral Ecology and Sociobiology, 59(6), 715–722. 10.1007/s00265-005-0101-0

Mangel, M., & Clark, C. W. (1988). Dynamic modeling in behavioral ecology. Princeton University Press.

Maruska, K. P., & Fernald, R. D. (2013). Social regulation of male reproductive plasticity in an African cichlid fish. Integrative and Comparative Biology, 53(6), 938–950. 10.1093/icb/ict017

McDermott, M. T., & Safran, R. J. (2021). Sensitive periods during the development and expression of vertebrate sexual signals: A systematic review. Ecology and Evolution, 11(21), 14416–14432. 10.1002/ece3.8203

McNamara, J. M., Green, R. F., & Olsson, O. (2006). Bayes’ theorem and its applications in animal behaviour. Oikos, 112(2), 243–251. 10.1111/j.0030-1299.2006.14228.x

Mcnamara, J. M., & Houston, A. (1980). The application of statistical decision theory to animal behaviour. Journal of Theoretical Biology, 85(4), 673–690.

Nettle, D., & Bateson, M. (2015). Adaptive developmental plasticity: What is it, how can we recognize it and when can it evolve? Proceedings of the Royal Society B: Biological Sciences, 282(1812), 20151005. 10.1098/rspb.2015.1005

Oyama, T., Sonoyama, T., Kasai, M., Sakai, Y., & Sunobe, T. (2023). Bidirectional sex change and plasticity of gonadal phases in the goby Lubricogobius exiguus. Journal of Fish Biology, 102(5), 1079–1087. 10.1111/jfb.15363

Panchanathan, K., & Frankenhuis, W. E. (2016). The evolution of sensitive periods in a model of incremental development. Proceedings of the Royal Society B: Biological Sciences, 283(1823). 10.1098/rspb.2015.2439

Penick, C. A., Ghaninia, M., Haight, K. L., Opachaloemphan, C., Yan, H., Reinberg, D., & Liebig, J. (2021). Reversible plasticity in brain size, behaviour and physiology characterizes caste transitions in a socially flexible ant (*Harpegnathos saltator*). Proceedings of the Royal Society B: Biological Sciences, 288(1948), rspb.2021.0141, 20210141. 10.1098/rspb.2021.0141

Pfab, F., Gabriel, W., & Utz, M. (2016). Reversible phenotypic plasticity with continuous adaptation. Journal of Mathematical Biology, 72(1–2), 435–466. 10.1007/s00285-015-0890-3

Piersma, T., & Drent, J. (2003). Phenotypic flexibility and the evolution of organismal design. Trends in Ecology & Evolution, 18(5), 228–233. 10.1016/S0169-5347(03)00036-3

Ratikainen, I. I., & Kokko, H. (2019). The coevolution of lifespan and reversible plasticity. Nature Communications, 10(1), 1–7. 10.1038/s41467-019-08502-9

Sourav, S., Kekunnaya, R., Bottari, D., Shareef, I., Pitchaimuthu, K., & Röder, B. (2024). Sound suppresses earliest visual cortical processing after sight recovery in congenitally blind humans. Communications Biology, 7(1), 1–14. 10.1038/s42003-023-05749-3

Stamps, J. A., & Frankenhuis, W. E. (2016). Bayesian models of development. Trends in Ecology & Evolution, 31(4), 260–268. 10.1016/j.tree.2016.01.012

Stamps, J. A., & Krishnan, V. V. (2014). Combining information from ancestors and personal experiences to predict individual differences in developmental trajectories. The American Naturalist, 184(5), 647–657. 10.1086/678116

Stamps, J. A., & Krishnan, V. V. (2017). Age-dependent changes in behavioural plasticity: Insights from Bayesian models of development. Animal Behaviour, 126, 53–67. 10.1016/j.anbehav.2017.01.013

Stamps, J. A., & Luttbeg, B. (2022). Sensitive period diversity: Insights from evolutionary models. The Quarterly Review of Biology, 97(4), 243–295. 10.1086/722637

Stearns, S. C. (1989). The evolutionary significance of phenotypic plasticity. Bioscience, 39(7), 436–445.

Sunobe, T., & Nakazono, A. (1993). Sex change in both directions by alteration of social dominance in trimma okinawae (Pisces: Gobiidae). Ethology, 94(4), 339–345. 10.1111/j.1439-0310.1993.tb00450.x

Trimmer, P. C., Houston, A. I., Marshall, J. A. R., Mendl, M. T., Paul, E. S., & McNamara, J. M. (2011). Decision-making under uncertainty: Biases and Bayesians. Animal Cognition, 14(4), 465–476. 10.1007/s10071-011-0387-4

Utz, M., Jeschke, J. M., Loeschcke, V., & Gabriel, W. (2014). Phenotypic plasticity with instantaneous but delayed switches. Journal of Theoretical Biology, 340, 60–72. 10.1016/j.jtbi.2013.08.038

Walasek, N., Frankenhuis, W. E., & Panchanathan, K. (2022a). An evolutionary model of sensitive periods when the reliability of cues varies across ontogeny. Behavioral Ecology, 33(1), 101–114. 10.1093/beheco/arab113

Walasek, N., Frankenhuis, W. E., & Panchanathan, K. (2022b). Sensitive periods, but not critical periods, evolve in a fluctuating environment: A model of incremental development. Proceedings of the Royal Society B: Biological Sciences, 289(1969), 20212623. 10.1098/rspb.2021.2623

Walasek, N., Panchanathan, K., & Frankenhuis, W. E. (in press). The evolution of sensitive periods beyond early ontogeny: Bridging theory and data. Functional Ecology. 10.1111/1365-2435.14615

West-Eberhard, M. J. (2003). Developmental plasticity and evolution. Oxford University Press.

Wikelski, M., & Thom, C. (2000). Marine iguanas shrink to survive El Niño. Nature, 403(6765), 37–38. 10.1038/47396

