## Supplementary material for "The evolution of reversible plasticity in stable environments": ESM

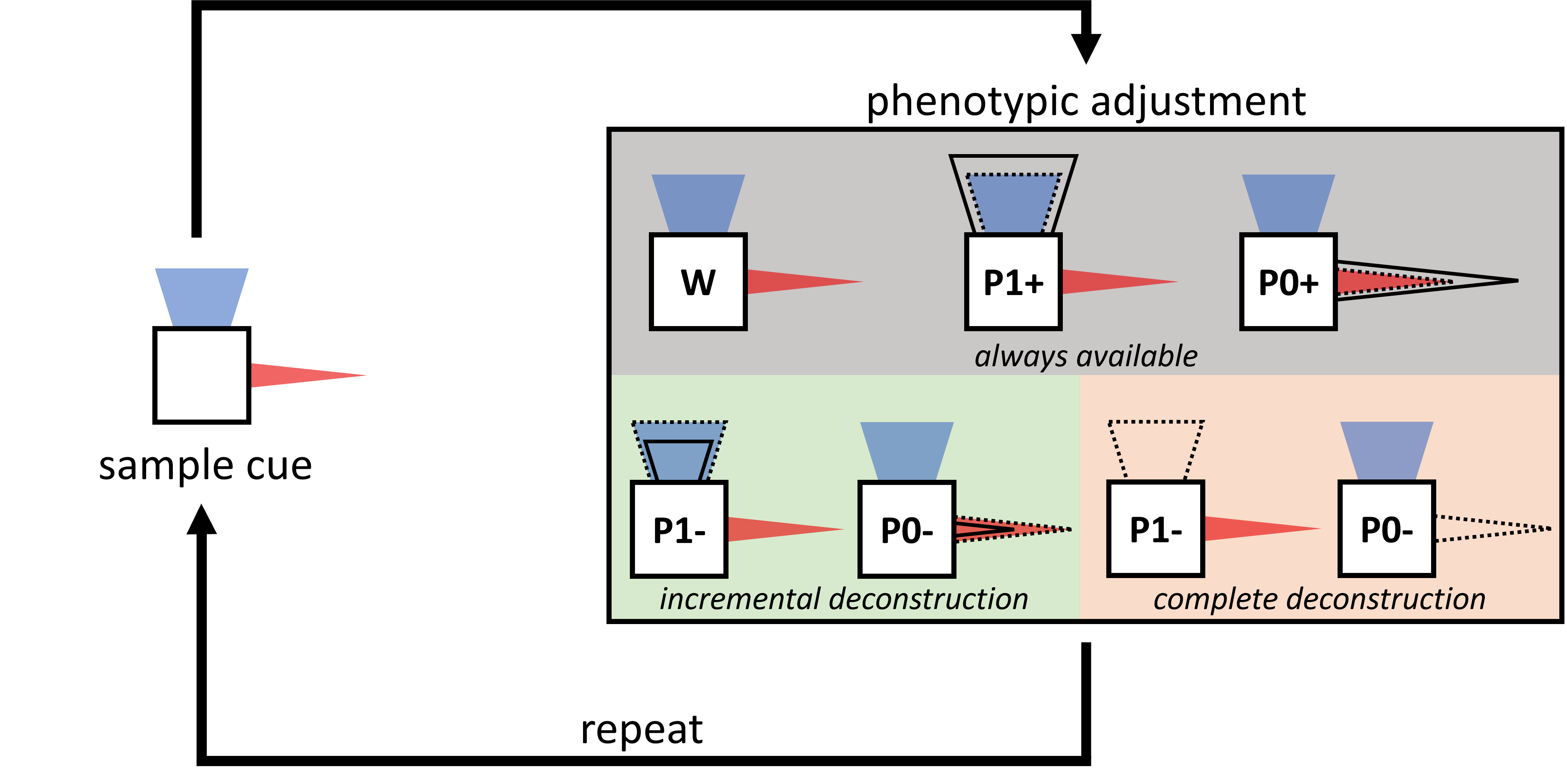

*Figure S1.1:* Overview of possible developmental decisions. We explore two models with different modes of deconstruction: incremental (green box) and complete (peach box). In either model, at any time period *t* during ontogeny, organisms can choose one of five phenotypic options. Organisms can wait and forgo phenotypic adjustments or incrementally specialize towards either phenotypic target (red cone, $P_{0}$ or blue helmet, $P_{1}$ ). These options are always available in both models (grey box). When deconstruction is incremental, organisms also choose to undo one phenotypic adjustment from either phenotypic target. When deconstruction is abrupt, organisms can choose to fully discard all phenotypic adjustments from either phenotypic target.

b) Decisions under uncertainty

Organisms use Bayesian inference to update their initial prior estimate of the environmental state based on the sampled cues.

| Parameters for Bayesian inference | Explanation |
| --- | --- |
| $P(E_{0})$ | Prior probability of $E_{0}$ |
| $P(E_{1})$ | Prior probability of $E_{1}$ |
| $P({C_{0}\vert E}_{0})$ | Cue reliability; conditional probability of receiving $C_{0}$in $E_{0}$ |
| $P({C_{1}\vert E}_{1})$ | Cue reliability; conditional probability of receiving $C_{1}$in $E_{1}$ |
| $P(E_{0}\vert D_{t})$ | Posterior probability of $E_{0}$ after having sampled $D_{t}$ |
| $P(E_{1}\vert D_{t})$ | Posterior probability of $E_{1}$ after having sampled $D_{t}$ |
| $P(D_{t}\vert E_{0})$ | Probability of observing the current cue set in $E_{0}.$ We use the binomial distribution to compute this probability. |
| $P(D_{t}\vert E_{1})$ | Probability of observing the current cue set in $E_{1}.$ We use the binomial distribution to compute this probability. |

According to the laws of probability it holds that:

$$P\left( E_{0} \right)+P\left( E_{1} \right)=1$$

$$P\left( E_{0}|D_{t} \right)+P\left( E_{1}|D_{t} \right)=1$$

$$P\left( {C_{0}|E}_{0} \right)+P\left( {C_{1}|E}_{0} \right)=1$$

$$P\left( {C_{1}|E}_{1} \right)+P\left( {C_{0}|E}_{1} \right)=1$$

Further, we assume that $P\left( {C_{0}|E}_{0} \right)= P\left( {C_{1}|E}_{1} \right).$

Based on this, we compute the posterior probabilities $P\left( E_{0}|D_{t} \right)$ and $P\left( E_{1}|D_{t} \right)$ according to:

$$P\left( E_{0}|D_{t} \right) = \frac{P\left( D_{t}|E_{0} \right)*P(E_{0})}{P\left( D_{t}|E_{0} \right)*P\left( E_{0} \right)+ P\left( D_{t}|E_{1} \right)*P(E_{1})}$$

$$P\left( E_{1}|D_{t} \right) = \frac{P\left( D_{t}|E_{1} \right)*P(E_{1})}{P\left( D_{t}|E_{0} \right)*P\left( E_{0} \right)+ P\left( D_{t}|E_{1} \right)*P(E_{1})}$$

c) Fitness functions

We denote the mature phenotype at the end of ontogeny by$Y_{mat}=\left( y_{0},y_{1}, T_{ont} \right)$.

| Functions and constants | Explanation |
| --- | --- |
| $\phi\left( Y_{mat} \right)$ | Expected, fitness reward at the end of ontogeny |
| $\psi\left( Y_{mat} \right)$ | Expected, fitness penalty at the end of ontogeny |
| $\pi(Y_{mat})$ | Expected fitness at the end of ontogeny |
| $\pi_{0}$ | Baseline fitness |
| $f\left( y \right)$ | Mapping between phenotypic increments and fitness rewards (or penalties) |

Fitness consequences of phenotypic decisions are not accrued throughout ontogeny but only at the end of ontogeny. The fitness difference from baseline corresponds to the total rewards for correct specializations minus penalties from incorrect specializations, where each correct increment results in a marginal gain and each incorrect increment results in a marginal penalty. We studied three mappings between correct (or incorrect) phenotypic development and fitness rewards (or penalties).

Suppose a mature organism is in the following state at the end of ontogeny ($D_{T_{ont}}, y_{0},y_{1}, y_{w}, T_{ont}$) having sampled the sequence of cues$D_{T_{ont}}$ and developed the mature phenotype $Y_{mat}=\{y_{0},y_{1}, T_{ont}\}.$ Its posterior estimates $P\left( E_{0,T_{ont}} | D_{T_{ont}} \right)$ and $P\left( E_{1, T_{ont}} | D_{T_{ont}} \right)$ at the end of ontogeny reflect the probabilities of being in either environmental state at the end of ontogeny.

To compute rewards and penalties, we need to compute the expectation across both environmental states, weighted by how likely each state is as indicated by the posterior estimates. We denote the mapping from phenotypic increments to rewards and penalties by $f\left( y \right)$, where $y$ can refer to both $y_{0}$ and $y_{1}$, and derive the following expressions for expected rewards and penalties at the end of ontogeny:

$$\phi\left( Y_{mat} \right)= P\left( E_{0,T_{ont}} | D_{T_{ont}} \right)\bullet f\left( y_{0} \right)+ P\left( E_{1,T_{ont}} | D_{T_{ont}} \right)\bullet f\left( y_{1} \right)$$

$$\psi\left( Y_{mat} \right)=-\left( P\left( E_{0, T_{ont}} | D_{T_{ont}} \right)\bullet f\left( y_{1} \right)+ P\left( E_{1,T_{ont}} | D_{T_{ont}} \right)\bullet f\left( y_{0} \right) \right)$$

Expected fitness $\pi\left( Y_{mat} \right)$ corresponds to the sum of expected rewards and penalties, in addition to the baseline fitness:

$\pi\left( Y_{mat} \right)= \pi_{0}+ \phi\left( Y_{mat} \right)+ \psi\left( Y_{mat} \right)$.

Lastly, we present the three functional mappings between the realized phenotype and fitness rewards and penalties:

| Returns on fitness - $f\left( y \right)$ | Formula | Parameter settings to ensure that maximal rewards and penalties correspond to $T_{ont}$ |
| --- | --- | --- |
| linear | $f\left( y \right)=y$ | - |
| diminishing | $f\left( y \right)=\alpha(1-e^{-\beta y})$ | $\beta=0.2, \alpha=\frac{T_{ont}}{{1-e}^{-\beta(T_{ont})}}$ |
| increasing | $f\left( y \right)=\alpha(e^{\beta y}-1)$ | $\beta=0.2, \alpha=\frac{T_{ont}}{e^{\beta\left( T_{ont} \right) - 1}}$ |

d) Optimal decisions

In each time period, a developing organism can choose one of five options: increment one step on $P_{0}$, increment one step on $P_{1},$ decrement one step on $P_{0}$, decrement one step on $P_{1},$or wait and forgo specialization. It chooses the option with the highest expected fitness at the end of ontogeny$.$ In the event of a tie between two or all of the options the organism chooses amongst the current alternatives with equal probability.

$F(D_{t},y_{0C},y_{1C},y_{0D},y_{1D}, y_{0},y_{1},y_{w}, t,T_{ont})$ denotes the maximum expected fitness that can be attained across adulthood as a result of decisions made between $t \mathrm{and}$ $T_{ont}$, when the organism’s current state after the last cue sampled is $(y_{0C},y_{1C},y_{0D},y_{1D}, y_{0},y_{1},y_{w})$ and the organisms chooses option $a$, so that:

$$F\left( D_{t},y_{0C},y_{1C},y_{0D},y_{1D}, y_{0},y_{1},y_{w}, t,T_{ont} \right)=\max_{a\epsilon\{0C,1C,0D,1D,w\}} F_{a}, \mathrm{where}$$

$$F_{0C}= F\left( D_{t+1},y_{0C}+1,y_{1C},y_{0D},y_{1D}, y_{0},y_{1}, y_{w},t+1,T_{ont} \right),$$

$$F_{1C}= F\left( D_{t+1},y_{0C},y_{1C}+1,y_{0D},y_{1D}, y_{0},y_{1},y_{w}, t+1,T_{ont} \right),$$

$$F_{0D}= F\left( D_{t+1},y_{0C},y_{1C},y_{0D}+1,y_{1D}, y_{0}-x,y_{1},y_{w}, t+1,T_{ont} \right),$$

$$F_{1D}= F\left( D_{t+1},y_{0C},y_{1C},y_{0D},y_{1D}+1, y_{0},y_{1}-x,y_{w}, t+1,T_{ont} \right),$$

$$F_{w}= F\left( D_{t+1},y_{0C},y_{1C},y_{0D},y_{1D}, y_{0},y_{1},y_{w}+1,t+1,T_{ont} \right).$$

Note, the value of $x$ depends on the mode of deconstruction. When deconstruction is incremental it corresponds to 1. When deconstruction is complete it corresponds to the current value of $y_{0}$ or $y_{1}$. Moreover, organisms can only choose to decrement one step on $P_{0}$ if $y_{0}>1.$Likewise, organisms can only choose to decrement one step on $P_{1}$ if $y_{1}>1$

We apply backwards induction to solve the dynamic programming equation $F\left( D_{t},y_{0C},y_{1C},y_{0D},y_{1D}, y_{0},y_{1},y_{w}, t,T_{ont} \right)$ for all $t.$ We start with $t=T_{ont}$:

$$F\left( D_{T_{ont}},y_{0C},y_{1C},y_{0D},y_{1D}, y_{0},y_{1},y_{w}, T_{ont},T_{ont} \right)= \pi(Y_{mat}).$$

After calculating expected fitness at the end of ontogeny we continue by decrementing $t$. For each $t<T_{ont}$ we compute the $a$, which maximizes $F\left( D_{t+1},y_{0C},y_{1C},y_{0D},y_{1D}, y_{0},y_{1},y_{w}, t+1,T_{ont} \right)$ in time period $t$.

ESM 2 - Main plots for all penalty and reward functions

**Optimal Policies (incremental deconstruction)**

*Linear rewards and linear penalties*

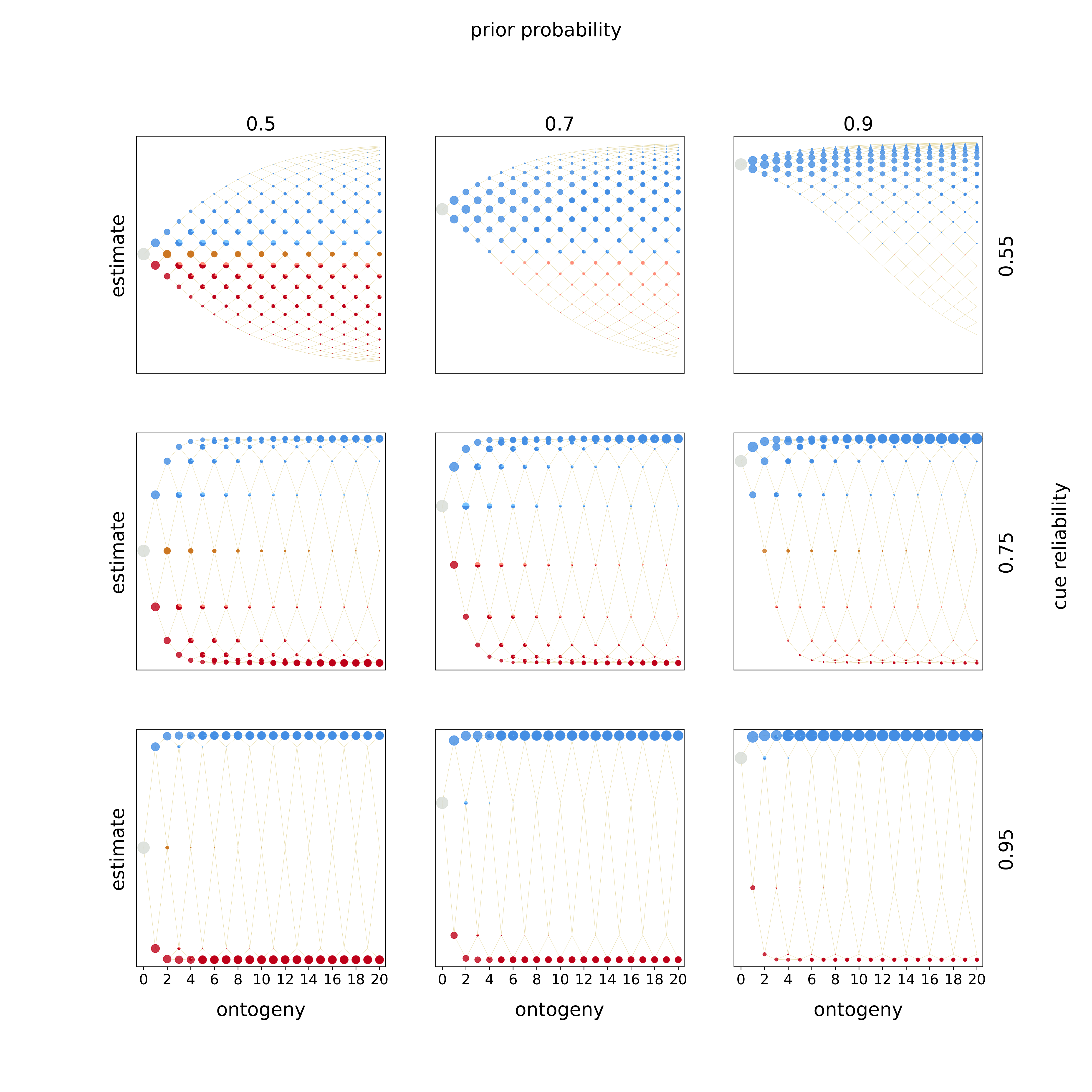

*Figure S2.1:* Optimal policies. Optimal policies are shown for a model with incremental deconstruction and linear rewards and penalties. Rows indicate the prior estimate of being in $E_{1}$ and columns indicate the cue reliability. Within each panel, the horizontal axis denotes ontogeny and the vertical axis the posterior estimates of being in $E_{1}$. The entire population starts ontogeny with zero cues sampled and the prior estimate indicated by the row (indicated by the grey circle). In each time period organisms sample a cue (either $C_{0}$ or $C_{1}$), update their estimate, and make a phenotypic decision (colored circles). Beige lines indicate developmental trajectories through this decision space, with lines branching upwards indicating the sampling of $C_{1}$ and lines branching downwards indicating the sampling of $C_{o}$. Colors denote the optimal, fitness-maximizing phenotypic choice in each state. Pies indicate cases in which organisms with the same posterior estimates make different phenotypic decisions. The area of a circle (pie piece) is proportional the probability of reaching that particular state. Colors indicate the following phenotypic decisions: Black corresponds to waiting, red to constructing $P_{0}$, blue to constructing $P_{1}$, purple to deconstructing $P_{0}$, green to deconstructing $P_{1}$, light red to a tie between constructing $P_{0}$ and deconstructing $P_{1}$, light blue to a tie between constructing $P_{1}$ and deconstructing $P_{0}$, brown to a tie between constructing either phenotypic target, yellow to a tie between deconstructing either target, grey to a tie between construction and waiting, dark grey to a tie between deconstruction and waiting, and lastly ochre to a tie between all options.

*Linear rewards and increasing penalties*

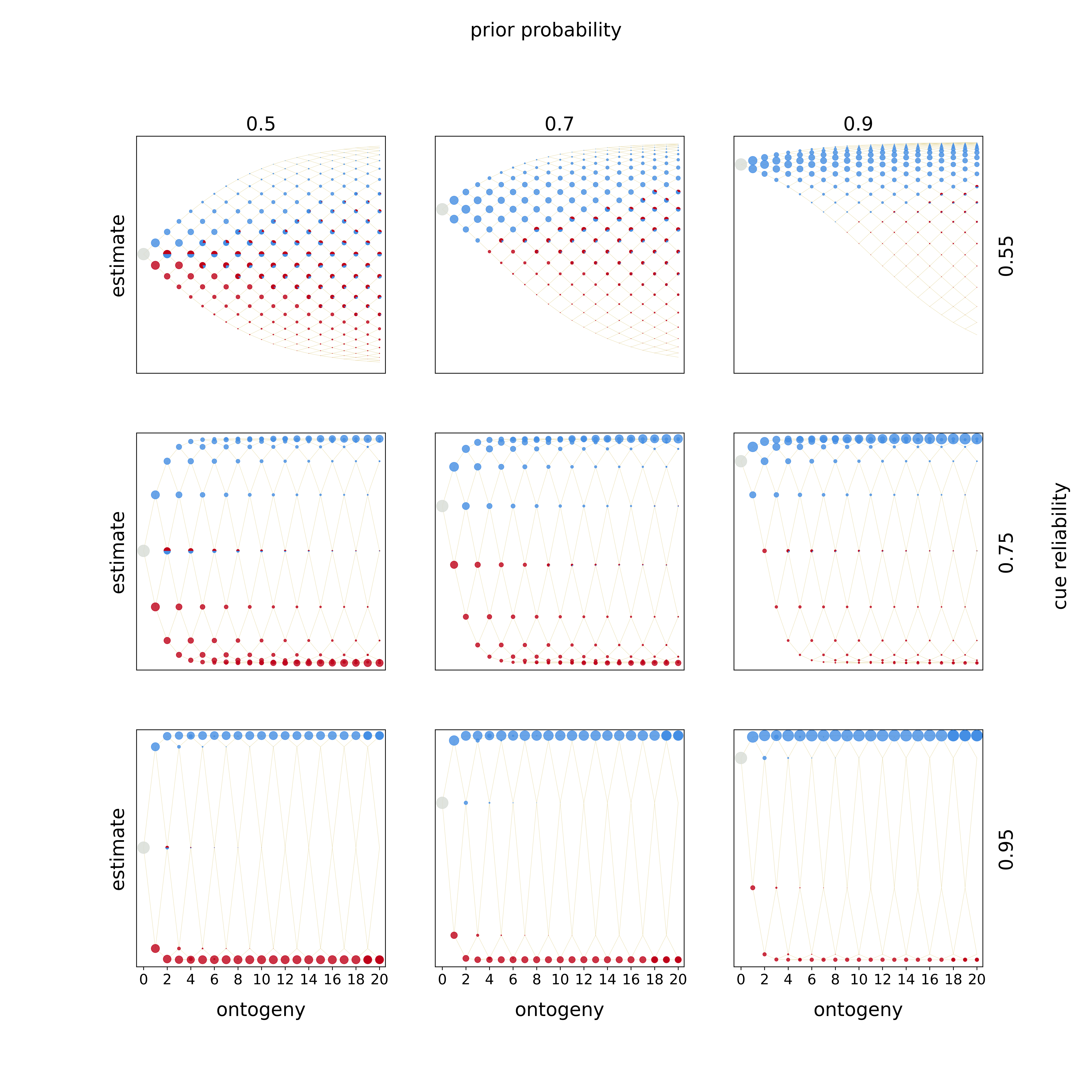

*Figure S2.2:* Optimal policies. Optimal policies are shown for a model with incremental deconstruction and linear rewards and increasing penalties. Rows indicate the prior estimate of being in $E_{1}$ and columns indicate the cue reliability. Within each panel, the horizontal axis denotes ontogeny and the vertical axis the posterior estimates of being in $E_{1}$. The entire population starts ontogeny with zero cues sampled and the prior estimate indicated by the row (indicated by the grey circle). In each time period organisms sample a cue (either $C_{0}$ or $C_{1}$), update their estimate, and make a phenotypic decision (colored circles). Beige lines indicate developmental trajectories through this decision space, with lines branching upwards indicating the sampling of $C_{1}$ and lines branching downwards indicating the sampling of $C_{o}$. Colors denote the optimal, fitness-maximizing phenotypic choice in each state. Pies indicate cases in which organisms with the same posterior estimates make different phenotypic decisions. The area of a circle (pie piece) is proportional the probability of reaching that particular state. Colors indicate the following phenotypic decisions: Black corresponds to waiting, red to constructing $P_{0}$, blue to constructing $P_{1}$, purple to deconstructing $P_{0}$, green to deconstructing $P_{1}$, light red to a tie between constructing $P_{0}$ and deconstructing $P_{1}$, light blue to a tie between constructing $P_{1}$ and deconstructing $P_{0}$, brown to a tie between constructing either phenotypic target, yellow to a tie between deconstructing either target, grey to a tie between construction and waiting, dark grey to a tie between deconstruction and waiting, and lastly ochre to a tie between all options.

*Linear rewards and diminishing penalties*

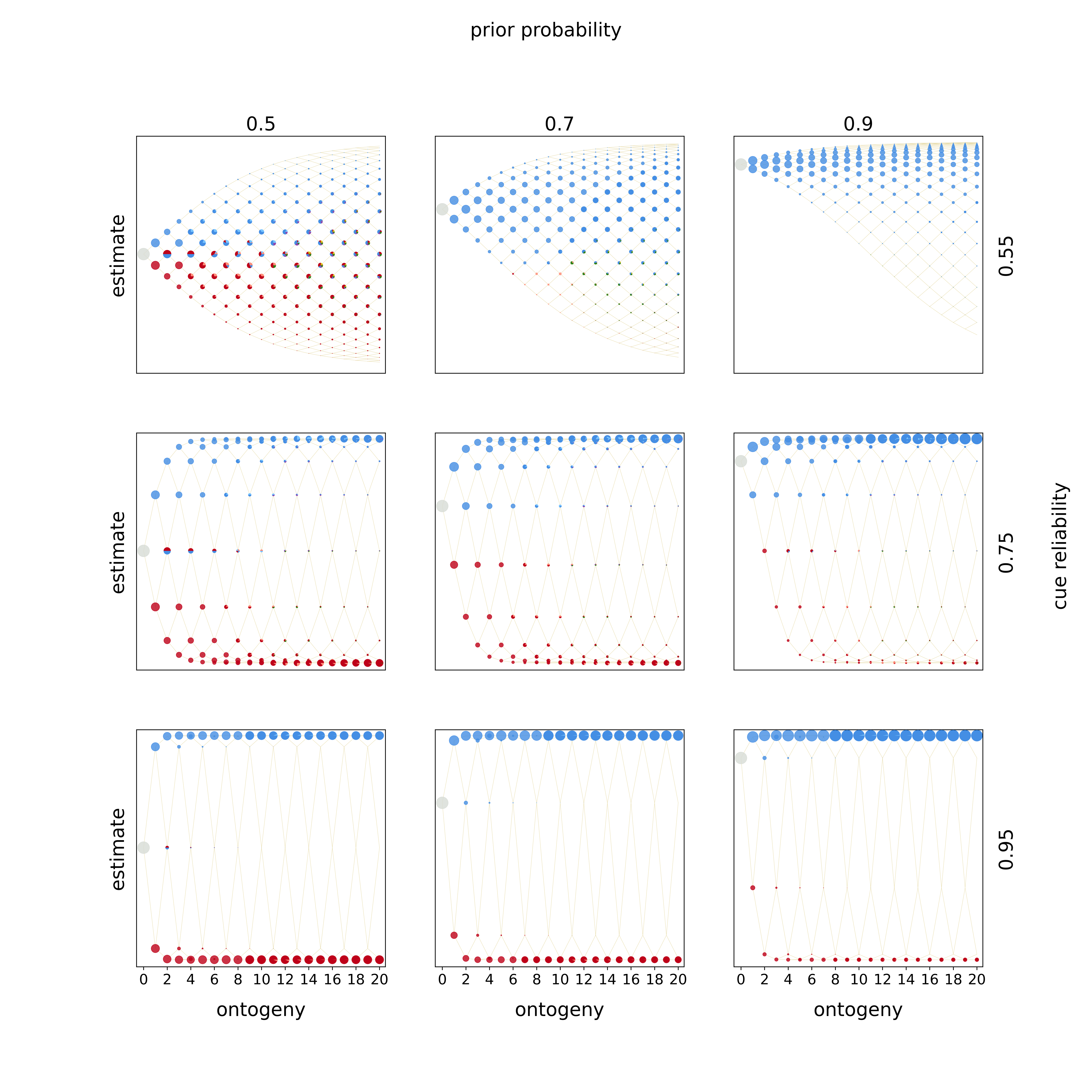

*Figure S2.3:* Optimal policies. Optimal policies are shown for a model with incremental deconstruction and linear rewards and diminishing penalties. Rows indicate the prior estimate of being in $E_{1}$ and columns indicate the cue reliability. Within each panel, the horizontal axis denotes ontogeny and the vertical axis the posterior estimates of being in $E_{1}$. The entire population starts ontogeny with zero cues sampled and the prior estimate indicated by the row (indicated by the grey circle). In each time period organisms sample a cue (either $C_{0}$ or $C_{1}$), update their estimate, and make a phenotypic decision (colored circles). Beige lines indicate developmental trajectories through this decision space, with lines branching upwards indicating the sampling of $C_{1}$ and lines branching downwards indicating the sampling of $C_{o}$. Colors denote the optimal, fitness-maximizing phenotypic choice in each state. Pies indicate cases in which organisms with the same posterior estimates make different phenotypic decisions. The area of a circle (pie piece) is proportional the probability of reaching that particular state. Colors indicate the following phenotypic decisions: Black corresponds to waiting, red to constructing $P_{0}$, blue to constructing $P_{1}$, purple to deconstructing $P_{0}$, green to deconstructing $P_{1}$, light red to a tie between constructing $P_{0}$ and deconstructing $P_{1}$, light blue to a tie between constructing $P_{1}$ and deconstructing $P_{0}$, brown to a tie between constructing either phenotypic target, yellow to a tie between deconstructing either target, grey to a tie between construction and waiting, dark grey to a tie between deconstruction and waiting, and lastly ochre to a tie between all options.

*Increasing rewards and linear penalties*

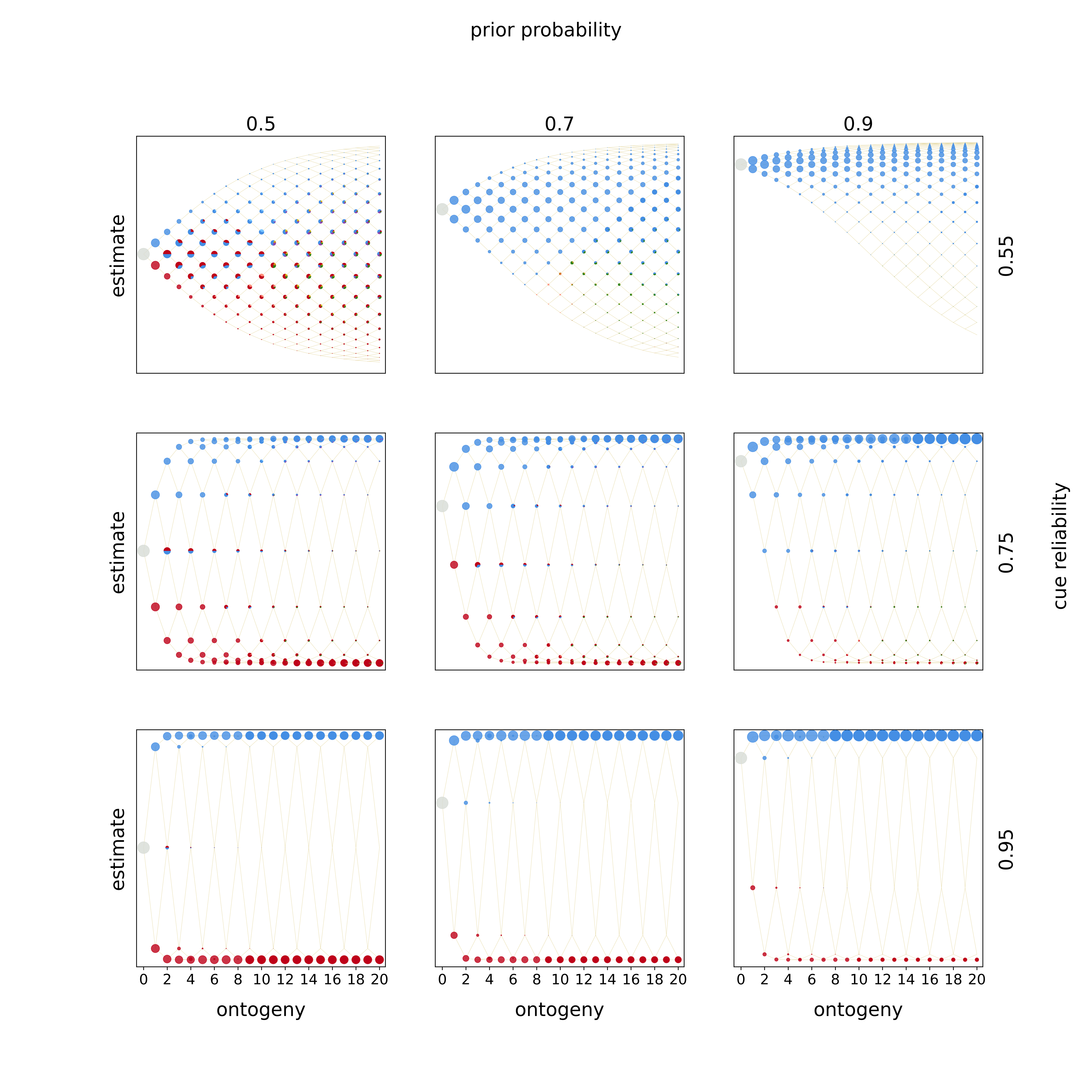

*Figure S2.4:* Optimal policies. Optimal policies are shown for a model with incremental deconstruction and increasing rewards and linear penalties. Rows indicate the prior estimate of being in $E_{1}$ and columns indicate the cue reliability. Within each panel, the horizontal axis denotes ontogeny and the vertical axis the posterior estimates of being in $E_{1}$. The entire population starts ontogeny with zero cues sampled and the prior estimate indicated by the row (indicated by the grey circle). In each time period organisms sample a cue (either $C_{0}$ or $C_{1}$), update their estimate, and make a phenotypic decision (colored circles). Beige lines indicate developmental trajectories through this decision space, with lines branching upwards indicating the sampling of $C_{1}$ and lines branching downwards indicating the sampling of $C_{o}$. Colors denote the optimal, fitness-maximizing phenotypic choice in each state. Pies indicate cases in which organisms with the same posterior estimates make different phenotypic decisions. The area of a circle (pie piece) is proportional the probability of reaching that particular state. Colors indicate the following phenotypic decisions: Black corresponds to waiting, red to constructing $P_{0}$, blue to constructing $P_{1}$, purple to deconstructing $P_{0}$, green to deconstructing $P_{1}$, light red to a tie between constructing $P_{0}$ and deconstructing $P_{1}$, light blue to a tie between constructing $P_{1}$ and deconstructing $P_{0}$, brown to a tie between constructing either phenotypic target, yellow to a tie between deconstructing either target, grey to a tie between construction and waiting, dark grey to a tie between deconstruction and waiting, and lastly ochre to a tie between all options.

*Increasing rewards and increasing penalties*

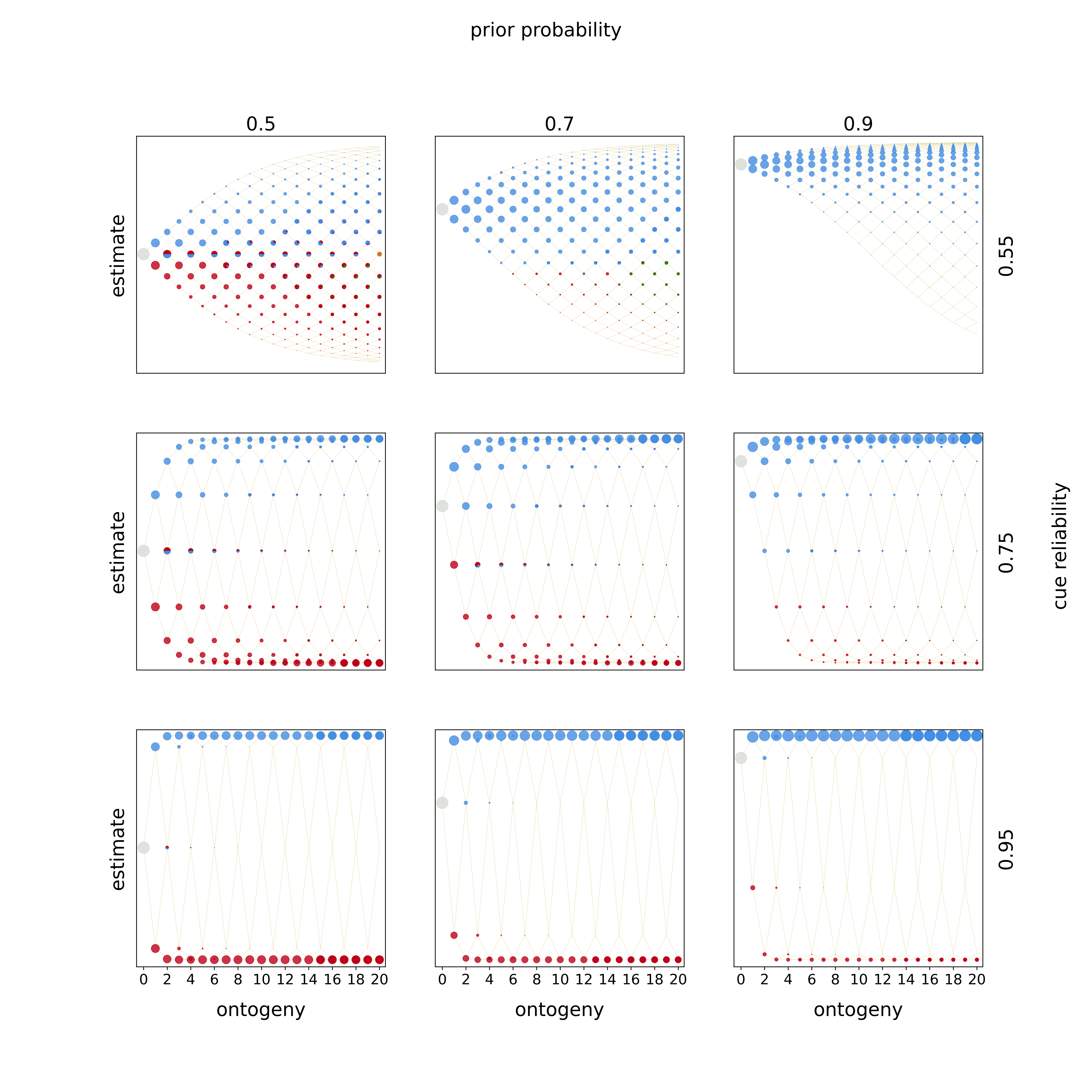

*Figure S2.5:* Optimal policies. Optimal policies are shown for a model with incremental deconstruction and increasing rewards and increasing penalties. Rows indicate the prior estimate of being in $E_{1}$ and columns indicate the cue reliability. Within each panel, the horizontal axis denotes ontogeny and the vertical axis the posterior estimates of being in $E_{1}$. The entire population starts ontogeny with zero cues sampled and the prior estimate indicated by the row (indicated by the grey circle). In each time period organisms sample a cue (either $C_{0}$ or $C_{1}$), update their estimate, and make a phenotypic decision (colored circles). Beige lines indicate developmental trajectories through this decision space, with lines branching upwards indicating the sampling of $C_{1}$ and lines branching downwards indicating the sampling of $C_{o}$. Colors denote the optimal, fitness-maximizing phenotypic choice in each state. Pies indicate cases in which organisms with the same posterior estimates make different phenotypic decisions. The area of a circle (pie piece) is proportional the probability of reaching that particular state. Colors indicate the following phenotypic decisions: Black corresponds to waiting, red to constructing $P_{0}$, blue to constructing $P_{1}$, purple to deconstructing $P_{0}$, green to deconstructing $P_{1}$, light red to a tie between constructing $P_{0}$ and deconstructing $P_{1}$, light blue to a tie between constructing $P_{1}$ and deconstructing $P_{0}$, brown to a tie between constructing either phenotypic target, yellow to a tie between deconstructing either target, grey to a tie between construction and waiting, dark grey to a tie between deconstruction and waiting, and lastly ochre to a tie between all options.

*Increasing rewards and diminishing penalties*

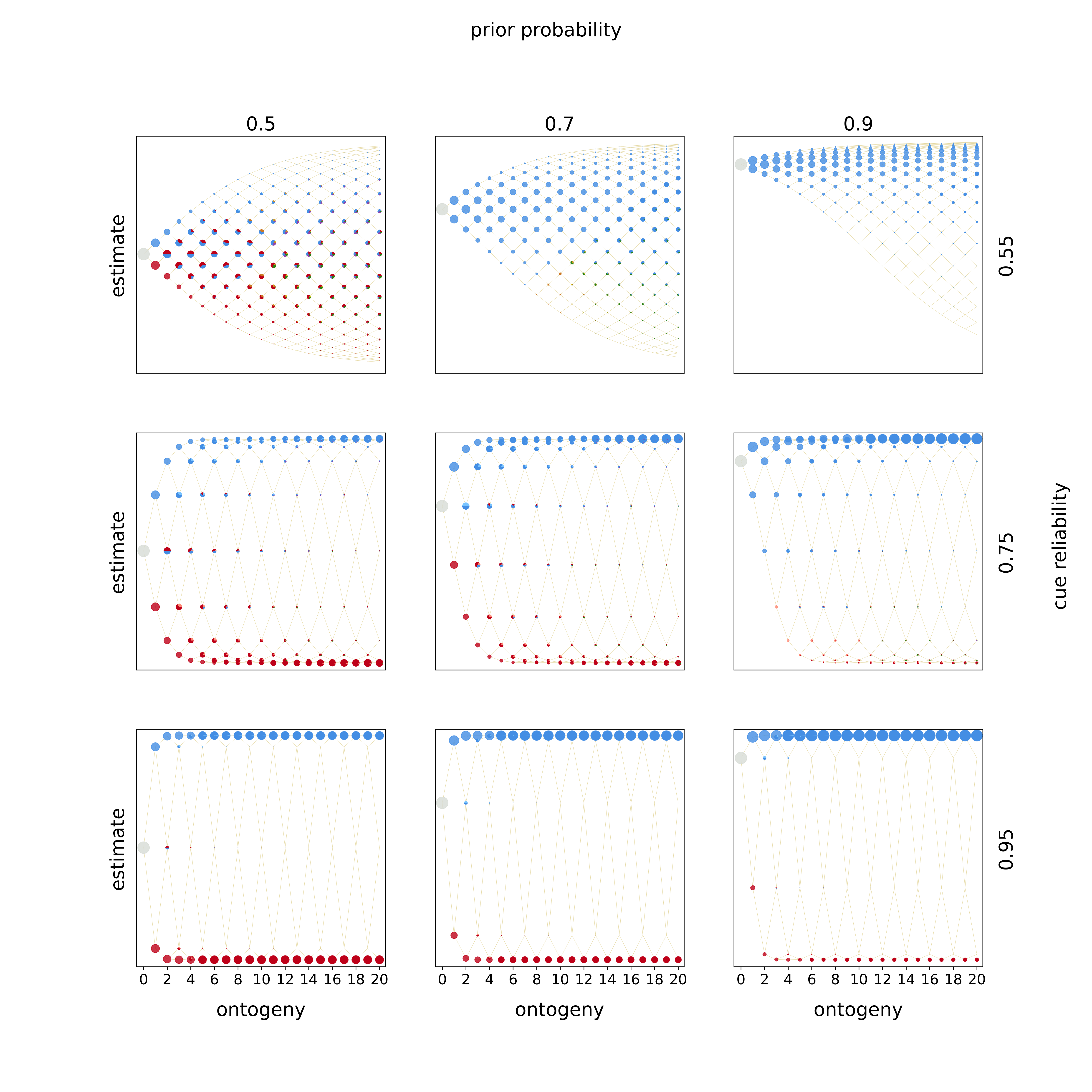

*Figure S2.6:* Optimal policies. Optimal policies are shown for a model with incremental deconstruction and increasing rewards and diminishing penalties. Rows indicate the prior estimate of being in $E_{1}$ and columns indicate the cue reliability. Within each panel, the horizontal axis denotes ontogeny and the vertical axis the posterior estimates of being in $E_{1}$. The entire population starts ontogeny with zero cues sampled and the prior estimate indicated by the row (indicated by the grey circle). In each time period organisms sample a cue (either $C_{0}$ or $C_{1}$), update their estimate, and make a phenotypic decision (colored circles). Beige lines indicate developmental trajectories through this decision space, with lines branching upwards indicating the sampling of $C_{1}$ and lines branching downwards indicating the sampling of $C_{o}$. Colors denote the optimal, fitness-maximizing phenotypic choice in each state. Pies indicate cases in which organisms with the same posterior estimates make different phenotypic decisions. The area of a circle (pie piece) is proportional the probability of reaching that particular state. Colors indicate the following phenotypic decisions: Black corresponds to waiting, red to constructing $P_{0}$, blue to constructing $P_{1}$, purple to deconstructing $P_{0}$, green to deconstructing $P_{1}$, light red to a tie between constructing $P_{0}$ and deconstructing $P_{1}$, light blue to a tie between constructing $P_{1}$ and deconstructing $P_{0}$, brown to a tie between constructing either phenotypic target, yellow to a tie between deconstructing either target, grey to a tie between construction and waiting, dark grey to a tie between deconstruction and waiting, and lastly ochre to a tie between all options.

*Diminishing rewards and linear penalties*

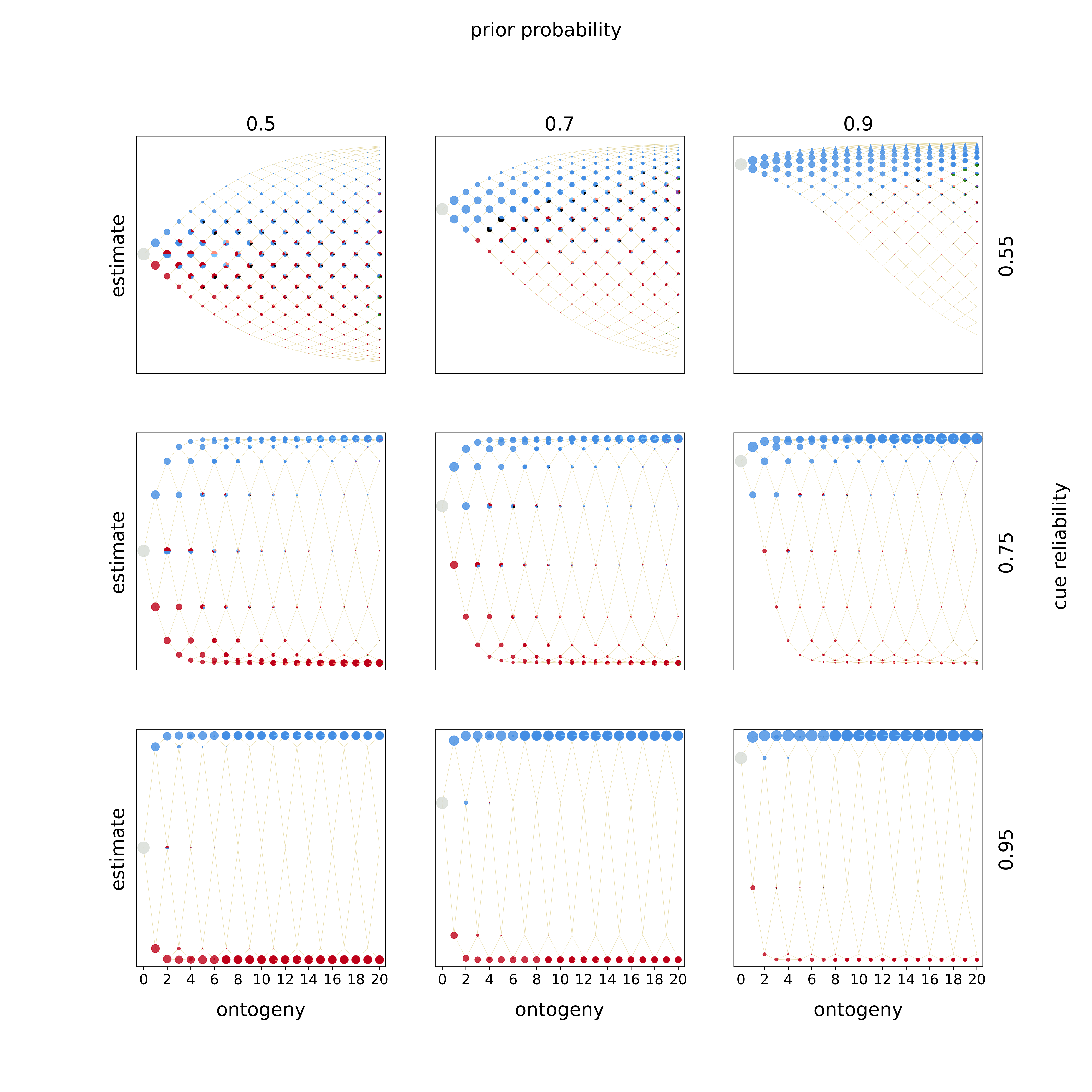

*Figure S2.7:* Optimal policies. Optimal policies are shown for a model with incremental deconstruction and diminishing rewards and linear penalties. Rows indicate the prior estimate of being in $E_{1}$ and columns indicate the cue reliability. Within each panel, the horizontal axis denotes ontogeny and the vertical axis the posterior estimates of being in $E_{1}$. The entire population starts ontogeny with zero cues sampled and the prior estimate indicated by the row (indicated by the grey circle). In each time period organisms sample a cue (either $C_{0}$ or $C_{1}$), update their estimate, and make a phenotypic decision (colored circles). Beige lines indicate developmental trajectories through this decision space, with lines branching upwards indicating the sampling of $C_{1}$ and lines branching downwards indicating the sampling of $C_{o}$. Colors denote the optimal, fitness-maximizing phenotypic choice in each state. Pies indicate cases in which organisms with the same posterior estimates make different phenotypic decisions. The area of a circle (pie piece) is proportional the probability of reaching that particular state. Colors indicate the following phenotypic decisions: Black corresponds to waiting, red to constructing $P_{0}$, blue to constructing $P_{1}$, purple to deconstructing $P_{0}$, green to deconstructing $P_{1}$, light red to a tie between constructing $P_{0}$ and deconstructing $P_{1}$, light blue to a tie between constructing $P_{1}$ and deconstructing $P_{0}$, brown to a tie between constructing either phenotypic target, yellow to a tie between deconstructing either target, grey to a tie between construction and waiting, dark grey to a tie between deconstruction and waiting, and lastly ochre to a tie between all options.

*Diminishing rewards and increasing penalties*

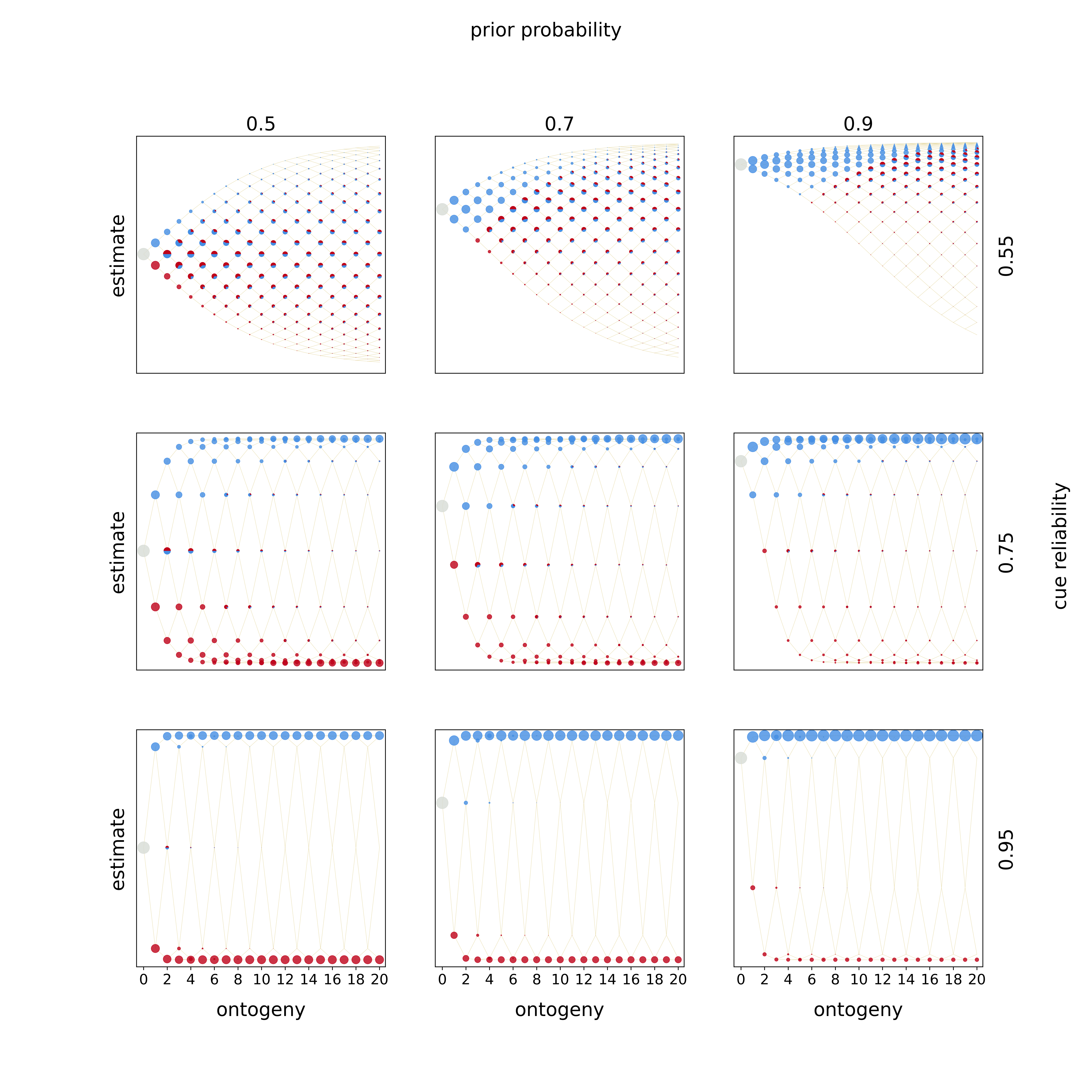

*Figure S2.8:* Optimal policies. Optimal policies are shown for a model with incremental deconstruction and diminishing rewards and increasing penalties. Rows indicate the prior estimate of being in $E_{1}$ and columns indicate the cue reliability. Within each panel, the horizontal axis denotes ontogeny and the vertical axis the posterior estimates of being in $E_{1}$. The entire population starts ontogeny with zero cues sampled and the prior estimate indicated by the row (indicated by the grey circle). In each time period organisms sample a cue (either $C_{0}$ or $C_{1}$), update their estimate, and make a phenotypic decision (colored circles). Beige lines indicate developmental trajectories through this decision space, with lines branching upwards indicating the sampling of $C_{1}$ and lines branching downwards indicating the sampling of $C_{o}$. Colors denote the optimal, fitness-maximizing phenotypic choice in each state. Pies indicate cases in which organisms with the same posterior estimates make different phenotypic decisions. The area of a circle (pie piece) is proportional the probability of reaching that particular state. Colors indicate the following phenotypic decisions: Black corresponds to waiting, red to constructing $P_{0}$, blue to constructing $P_{1}$, purple to deconstructing $P_{0}$, green to deconstructing $P_{1}$, light red to a tie between constructing $P_{0}$ and deconstructing $P_{1}$, light blue to a tie between constructing $P_{1}$ and deconstructing $P_{0}$, brown to a tie between constructing either phenotypic target, yellow to a tie between deconstructing either target, grey to a tie between construction and waiting, dark grey to a tie between deconstruction and waiting, and lastly ochre to a tie between all options.

*Diminishing rewards and diminishing penalties*

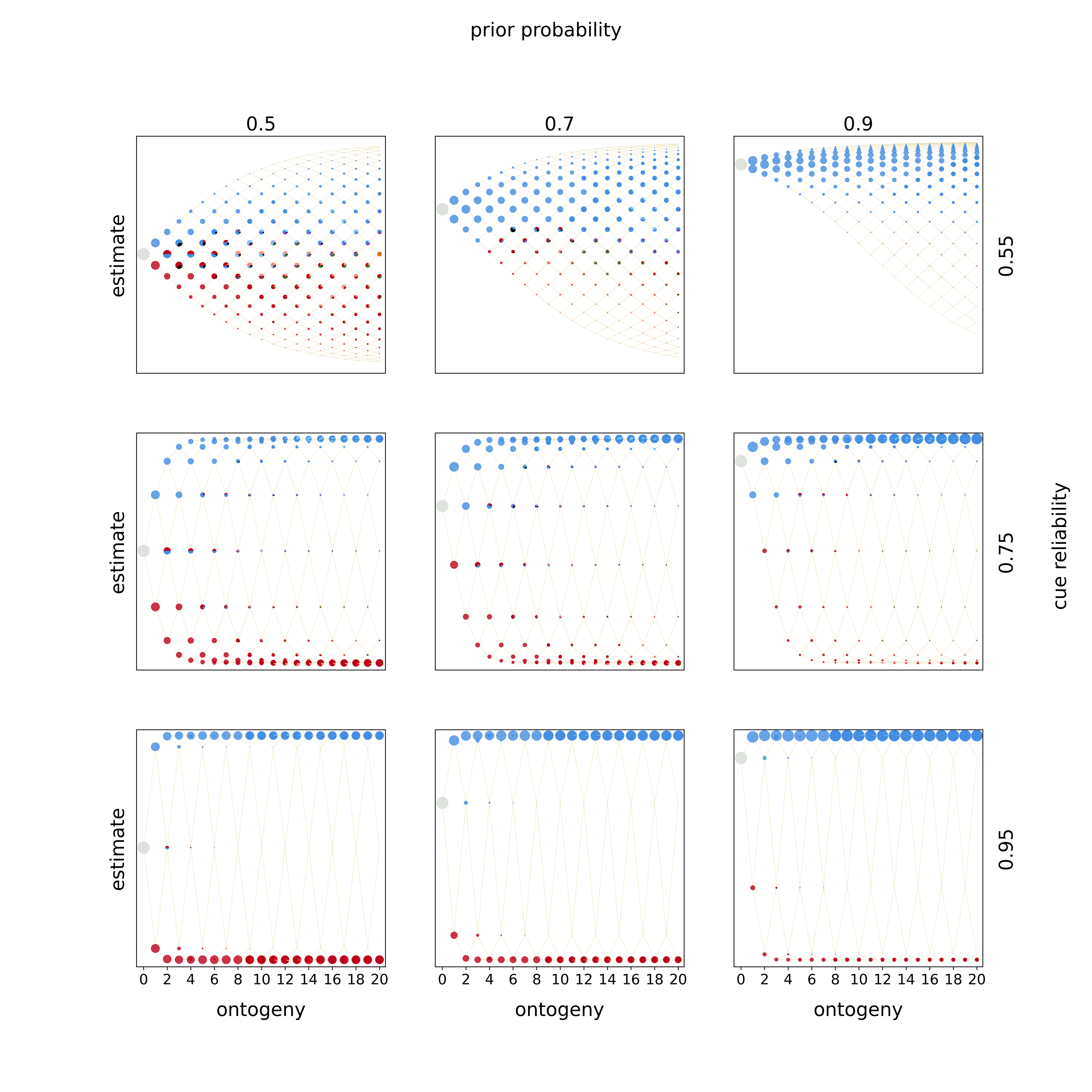

*Figure S2.9:* Optimal policies. Optimal policies are shown for a model with incremental deconstruction and diminishing rewards and diminishing penalties. Rows indicate the prior estimate of being in $E_{1}$ and columns indicate the cue reliability. Within each panel, the horizontal axis denotes ontogeny and the vertical axis the posterior estimates of being in $E_{1}$. The entire population starts ontogeny with zero cues sampled and the prior estimate indicated by the row (indicated by the grey circle). In each time period organisms sample a cue (either $C_{0}$ or $C_{1}$), update their estimate, and make a phenotypic decision (colored circles). Beige lines indicate developmental trajectories through this decision space, with lines branching upwards indicating the sampling of $C_{1}$ and lines branching downwards indicating the sampling of $C_{o}$. Colors denote the optimal, fitness-maximizing phenotypic choice in each state. Pies indicate cases in which organisms with the same posterior estimates make different phenotypic decisions. The area of a circle (pie piece) is proportional the probability of reaching that particular state. Colors indicate the following phenotypic decisions: Black corresponds to waiting, red to constructing $P_{0}$, blue to constructing $P_{1}$, purple to deconstructing $P_{0}$, green to deconstructing $P_{1}$, light red to a tie between constructing $P_{0}$ and deconstructing $P_{1}$, light blue to a tie between constructing $P_{1}$ and deconstructing $P_{0}$, brown to a tie between constructing either phenotypic target, yellow to a tie between deconstructing either target, grey to a tie between construction and waiting, dark grey to a tie between deconstruction and waiting, and lastly ochre to a tie between all options.

**Optimal Policies (complete deconstruction)**

*Linear rewards and linear penalties*

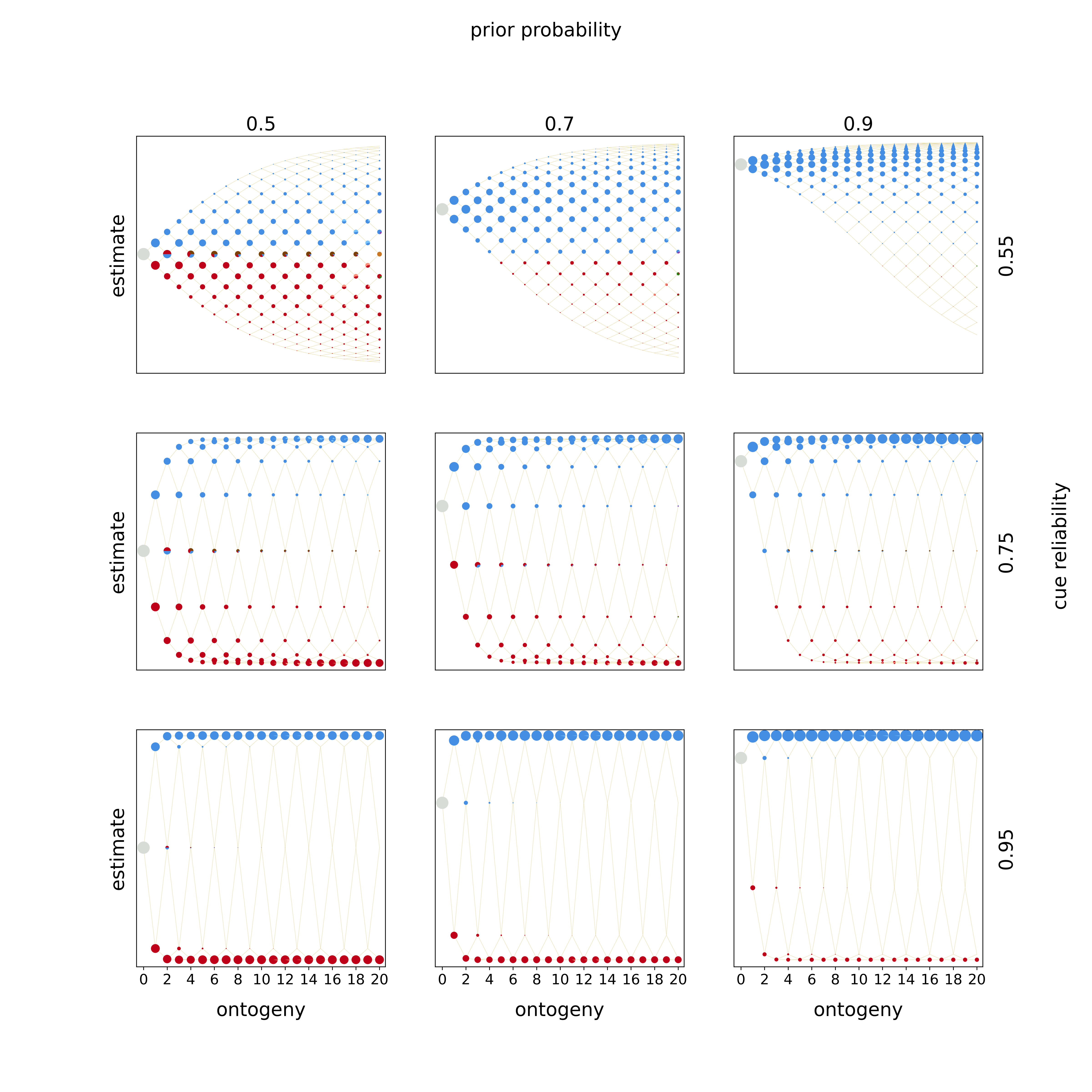

*Figure S2.10:* Optimal policies. Optimal policies are shown for a model with complete deconstruction and linear rewards and penalties. Rows indicate the prior estimate of being in $E_{1}$ and columns indicate the cue reliability. Within each panel, the horizontal axis denotes ontogeny and the vertical axis the posterior estimates of being in $E_{1}$. The entire population starts ontogeny with zero cues sampled and the prior estimate indicated by the row (indicated by the grey circle). In each time period organisms sample a cue (either $C_{0}$ or $C_{1}$), update their estimate, and make a phenotypic decision (colored circles). Beige lines indicate developmental trajectories through this decision space, with lines branching upwards indicating the sampling of $C_{1}$ and lines branching downwards indicating the sampling of $C_{o}$. Colors denote the optimal, fitness-maximizing phenotypic choice in each state. Pies indicate cases in which organisms with the same posterior estimates make different phenotypic decisions. The area of a circle (pie piece) is proportional the probability of reaching that particular state. Colors indicate the following phenotypic decisions: Black corresponds to waiting, red to constructing $P_{0}$, blue to constructing $P_{1}$, purple to deconstructing $P_{0}$, green to deconstructing $P_{1}$, light red to a tie between constructing $P_{0}$ and deconstructing $P_{1}$, light blue to a tie between constructing $P_{1}$ and deconstructing $P_{0}$, brown to a tie between constructing either phenotypic target, yellow to a tie between deconstructing either target, grey to a tie between construction and waiting, dark grey to a tie between deconstruction and waiting, and lastly ochre to a tie between all options.

*Linear rewards and increasing penalties*

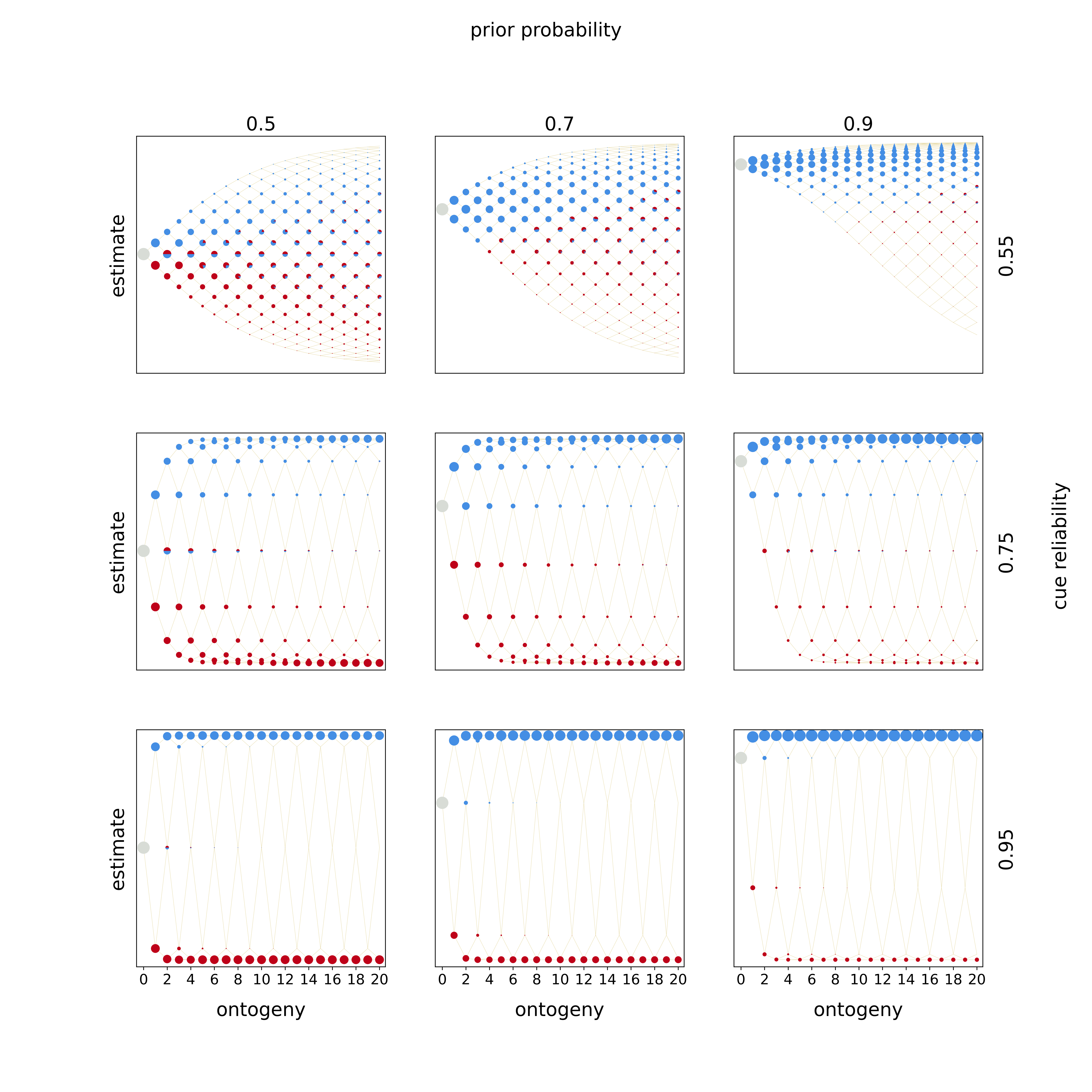

*Figure S2.11:* Optimal policies. Optimal policies are shown for a model with complete deconstruction and linear rewards and increasing penalties. Rows indicate the prior estimate of being in $E_{1}$ and columns indicate the cue reliability. Within each panel, the horizontal axis denotes ontogeny and the vertical axis the posterior estimates of being in $E_{1}$. The entire population starts ontogeny with zero cues sampled and the prior estimate indicated by the row (indicated by the grey circle). In each time period organisms sample a cue (either $C_{0}$ or $C_{1}$), update their estimate, and make a phenotypic decision (colored circles). Beige lines indicate developmental trajectories through this decision space, with lines branching upwards indicating the sampling of $C_{1}$ and lines branching downwards indicating the sampling of $C_{o}$. Colors denote the optimal, fitness-maximizing phenotypic choice in each state. Pies indicate cases in which organisms with the same posterior estimates make different phenotypic decisions. The area of a circle (pie piece) is proportional the probability of reaching that particular state. Colors indicate the following phenotypic decisions: Black corresponds to waiting, red to constructing $P_{0}$, blue to constructing $P_{1}$, purple to deconstructing $P_{0}$, green to deconstructing $P_{1}$, light red to a tie between constructing $P_{0}$ and deconstructing $P_{1}$, light blue to a tie between constructing $P_{1}$ and deconstructing $P_{0}$, brown to a tie between constructing either phenotypic target, yellow to a tie between deconstructing either target, grey to a tie between construction and waiting, dark grey to a tie between deconstruction and waiting, and lastly ochre to a tie between all options.

*Linear rewards and diminishing penalties*

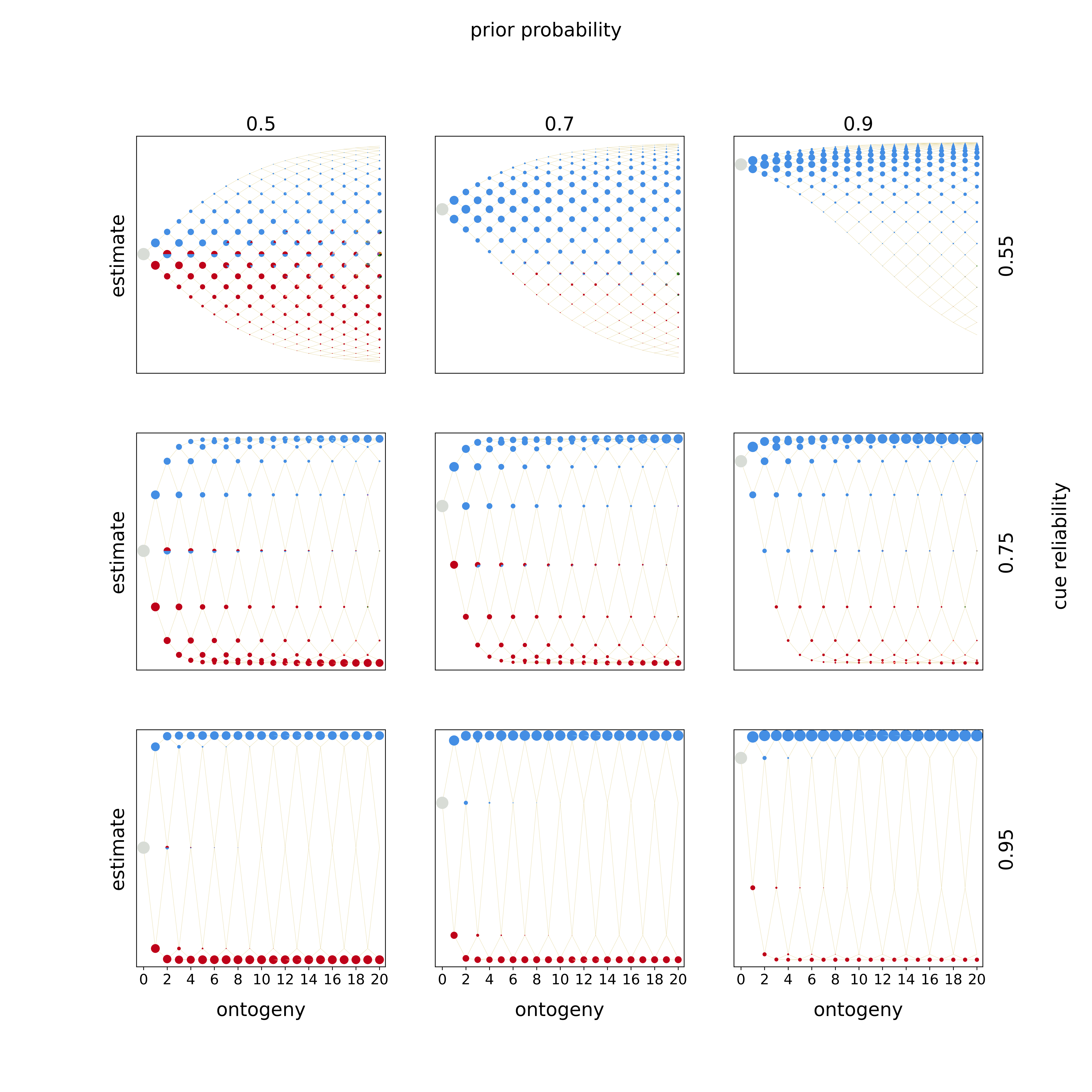

*Figure S2.12:* Optimal policies. Optimal policies are shown for a model with complete deconstruction and linear rewards and diminishing penalties. Rows indicate the prior estimate of being in $E_{1}$ and columns indicate the cue reliability. Within each panel, the horizontal axis denotes ontogeny and the vertical axis the posterior estimates of being in $E_{1}$. The entire population starts ontogeny with zero cues sampled and the prior estimate indicated by the row (indicated by the grey circle). In each time period organisms sample a cue (either $C_{0}$ or $C_{1}$), update their estimate, and make a phenotypic decision (colored circles). Beige lines indicate developmental trajectories through this decision space, with lines branching upwards indicating the sampling of $C_{1}$ and lines branching downwards indicating the sampling of $C_{o}$. Colors denote the optimal, fitness-maximizing phenotypic choice in each state. Pies indicate cases in which organisms with the same posterior estimates make different phenotypic decisions. The area of a circle (pie piece) is proportional the probability of reaching that particular state. Colors indicate the following phenotypic decisions: Black corresponds to waiting, red to constructing $P_{0}$, blue to constructing $P_{1}$, purple to deconstructing $P_{0}$, green to deconstructing $P_{1}$, light red to a tie between constructing $P_{0}$ and deconstructing $P_{1}$, light blue to a tie between constructing $P_{1}$ and deconstructing $P_{0}$, brown to a tie between constructing either phenotypic target, yellow to a tie between deconstructing either target, grey to a tie between construction and waiting, dark grey to a tie between deconstruction and waiting, and lastly ochre to a tie between all options.

*Increasing rewards and linear penalties*

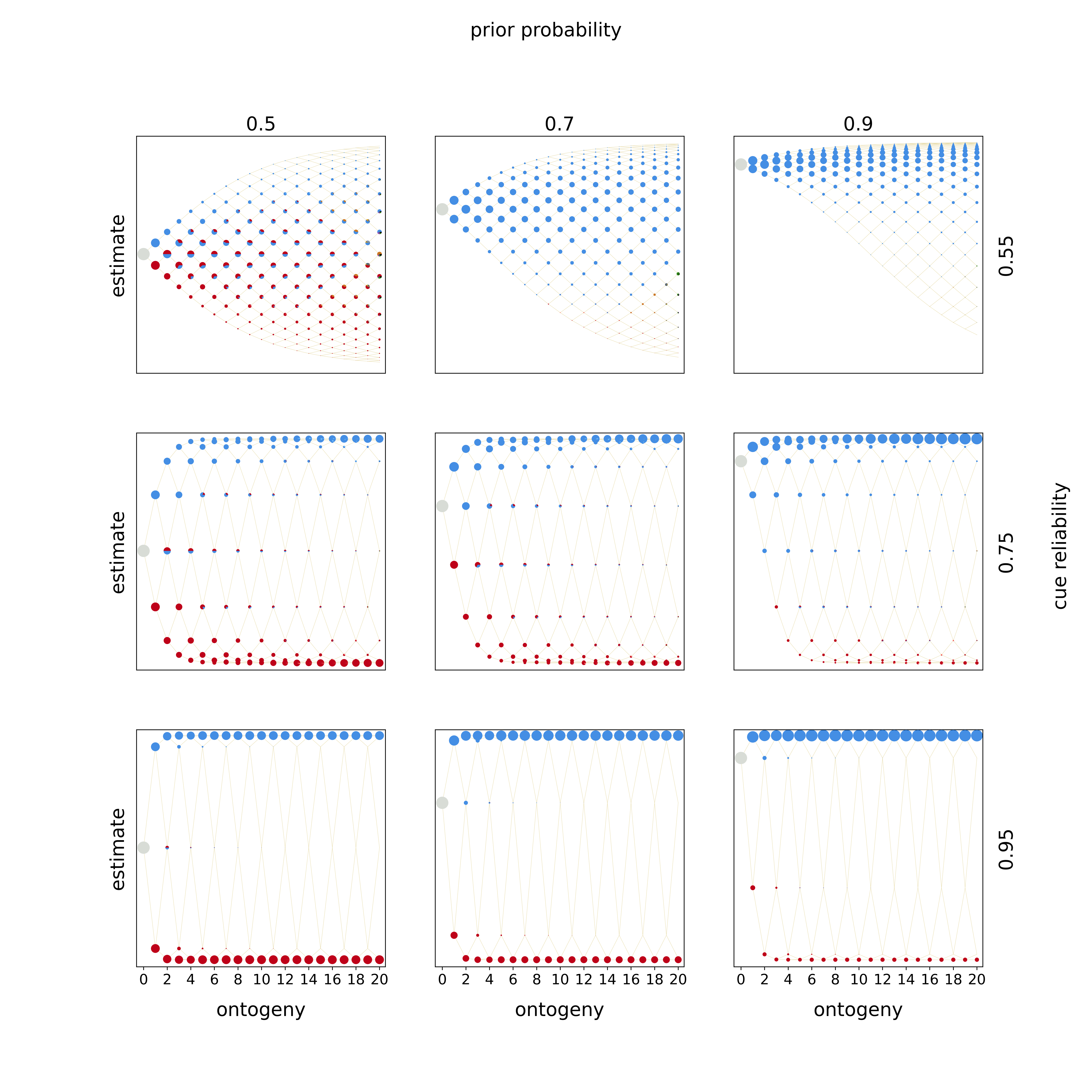

*Figure S2.13:* Optimal policies. Optimal policies are shown for a model with complete deconstruction and increasing rewards and linearpenalties. Rows indicate the prior estimate of being in $E_{1}$ and columns indicate the cue reliability. Within each panel, the horizontal axis denotes ontogeny and the vertical axis the posterior estimates of being in $E_{1}$. The entire population starts ontogeny with zero cues sampled and the prior estimate indicated by the row (indicated by the grey circle). In each time period organisms sample a cue (either $C_{0}$ or $C_{1}$), update their estimate, and make a phenotypic decision (colored circles). Beige lines indicate developmental trajectories through this decision space, with lines branching upwards indicating the sampling of $C_{1}$ and lines branching downwards indicating the sampling of $C_{o}$. Colors denote the optimal, fitness-maximizing phenotypic choice in each state. Pies indicate cases in which organisms with the same posterior estimates make different phenotypic decisions. The area of a circle (pie piece) is proportional the probability of reaching that particular state. Colors indicate the following phenotypic decisions: Black corresponds to waiting, red to constructing $P_{0}$, blue to constructing $P_{1}$, purple to deconstructing $P_{0}$, green to deconstructing $P_{1}$, light red to a tie between constructing $P_{0}$ and deconstructing $P_{1}$, light blue to a tie between constructing $P_{1}$ and deconstructing $P_{0}$, brown to a tie between constructing either phenotypic target, yellow to a tie between deconstructing either target, grey to a tie between construction and waiting, dark grey to a tie between deconstruction and waiting, and lastly ochre to a tie between all options.

*Increasing rewards and increasing penalties*

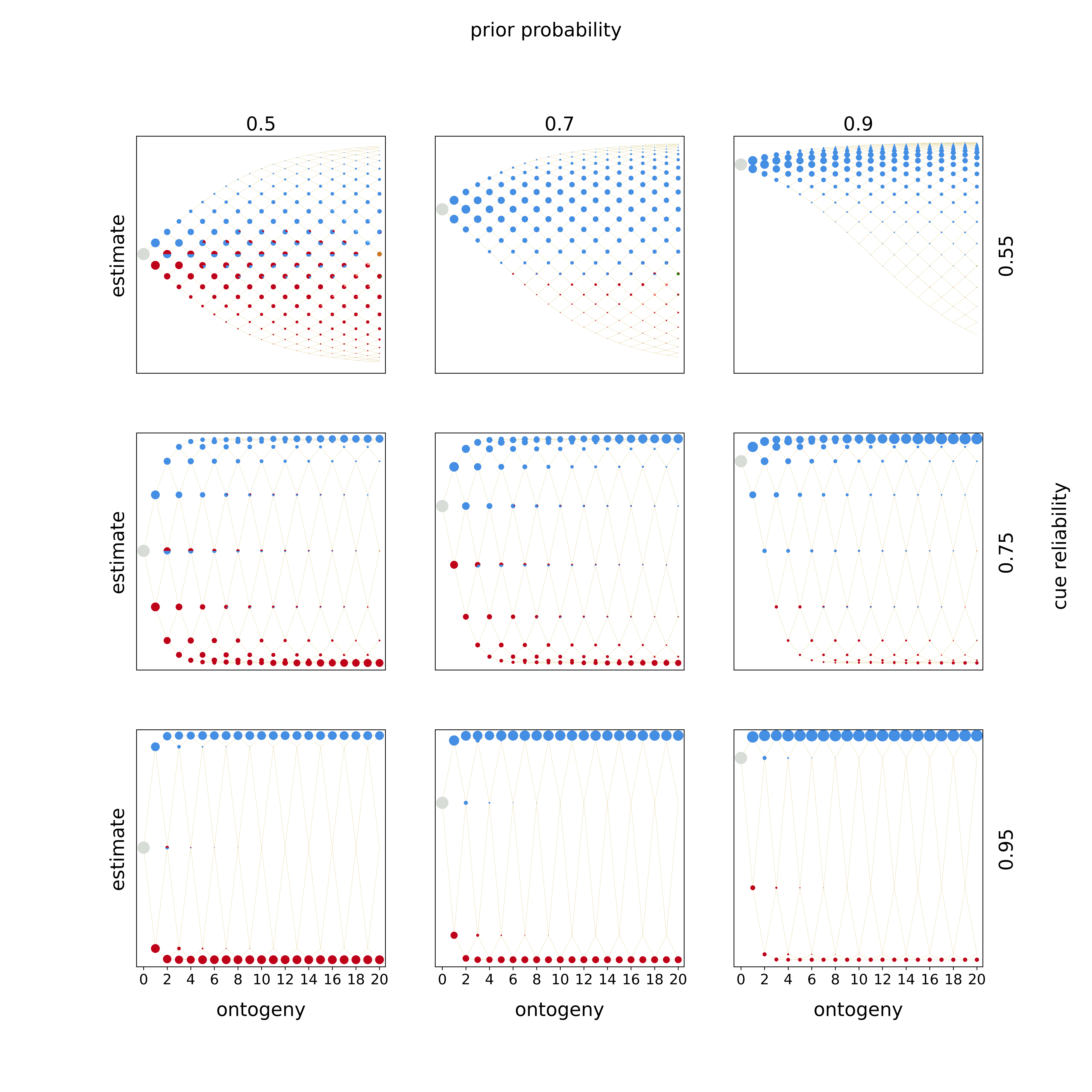

*Figure S2.14:* Optimal policies. Optimal policies are shown for a model with complete deconstruction and increasing rewards and increasing penalties. Rows indicate the prior estimate of being in $E_{1}$ and columns indicate the cue reliability. Within each panel, the horizontal axis denotes ontogeny and the vertical axis the posterior estimates of being in $E_{1}$. The entire population starts ontogeny with zero cues sampled and the prior estimate indicated by the row (indicated by the grey circle). In each time period organisms sample a cue (either $C_{0}$ or $C_{1}$), update their estimate, and make a phenotypic decision (colored circles). Beige lines indicate developmental trajectories through this decision space, with lines branching upwards indicating the sampling of $C_{1}$ and lines branching downwards indicating the sampling of $C_{o}$. Colors denote the optimal, fitness-maximizing phenotypic choice in each state. Pies indicate cases in which organisms with the same posterior estimates make different phenotypic decisions. The area of a circle (pie piece) is proportional the probability of reaching that particular state. Colors indicate the following phenotypic decisions: Black corresponds to waiting, red to constructing $P_{0}$, blue to constructing $P_{1}$, purple to deconstructing $P_{0}$, green to deconstructing $P_{1}$, light red to a tie between constructing $P_{0}$ and deconstructing $P_{1}$, light blue to a tie between constructing $P_{1}$ and deconstructing $P_{0}$, brown to a tie between constructing either phenotypic target, yellow to a tie between deconstructing either target, grey to a tie between construction and waiting, dark grey to a tie between deconstruction and waiting, and lastly ochre to a tie between all options.

*Increasing rewards and diminishing penalties*

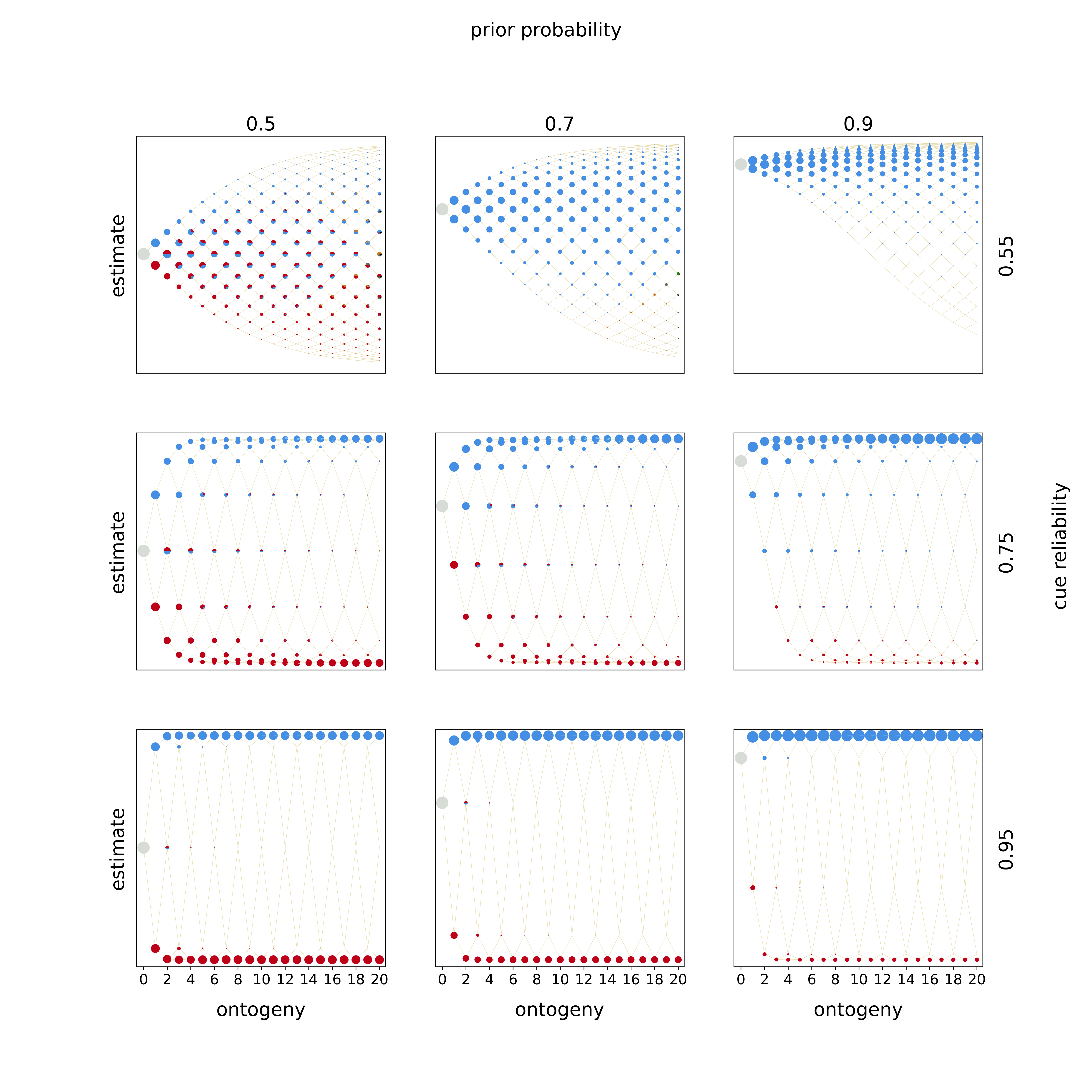

*Figure S2.15:* Optimal policies. Optimal policies are shown for a model with complete deconstruction and increasing rewards and diminishing penalties. Rows indicate the prior estimate of being in $E_{1}$ and columns indicate the cue reliability. Within each panel, the horizontal axis denotes ontogeny and the vertical axis the posterior estimates of being in $E_{1}$. The entire population starts ontogeny with zero cues sampled and the prior estimate indicated by the row (indicated by the grey circle). In each time period organisms sample a cue (either $C_{0}$ or $C_{1}$), update their estimate, and make a phenotypic decision (colored circles). Beige lines indicate developmental trajectories through this decision space, with lines branching upwards indicating the sampling of $C_{1}$ and lines branching downwards indicating the sampling of $C_{o}$. Colors denote the optimal, fitness-maximizing phenotypic choice in each state. Pies indicate cases in which organisms with the same posterior estimates make different phenotypic decisions. The area of a circle (pie piece) is proportional the probability of reaching that particular state. Colors indicate the following phenotypic decisions: Black corresponds to waiting, red to constructing $P_{0}$, blue to constructing $P_{1}$, purple to deconstructing $P_{0}$, green to deconstructing $P_{1}$, light red to a tie between constructing $P_{0}$ and deconstructing $P_{1}$, light blue to a tie between constructing $P_{1}$ and deconstructing $P_{0}$, brown to a tie between constructing either phenotypic target, yellow to a tie between deconstructing either target, grey to a tie between construction and waiting, dark grey to a tie between deconstruction and waiting, and lastly ochre to a tie between all options.

*Diminishing rewards and linear penalties*

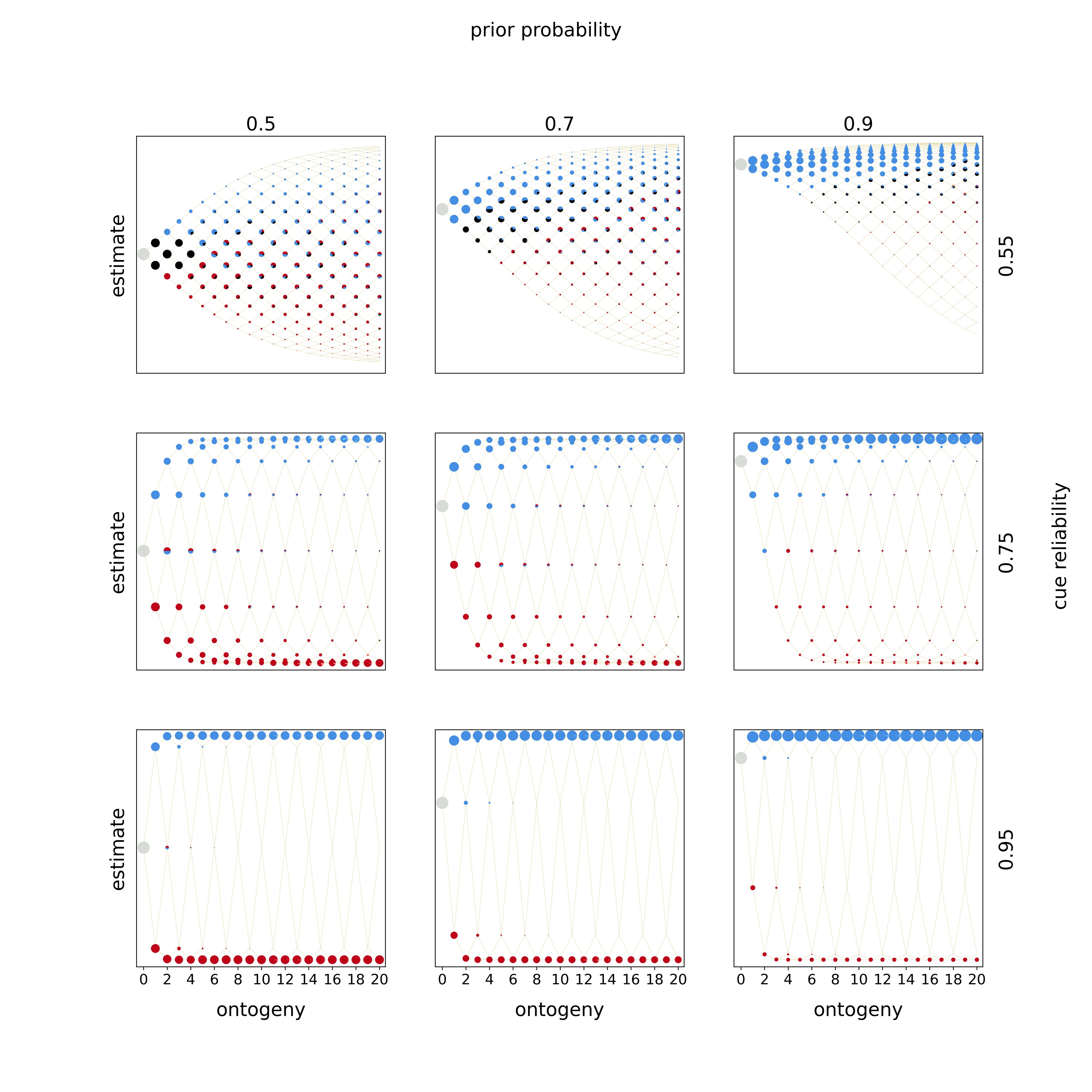

*Figure S2.16:* Optimal policies. Optimal policies are shown for a model with complete deconstruction and diminishing rewards and linear penalties. Rows indicate the prior estimate of being in $E_{1}$ and columns indicate the cue reliability. Within each panel, the horizontal axis denotes ontogeny and the vertical axis the posterior estimates of being in $E_{1}$. The entire population starts ontogeny with zero cues sampled and the prior estimate indicated by the row (indicated by the grey circle). In each time period organisms sample a cue (either $C_{0}$ or $C_{1}$), update their estimate, and make a phenotypic decision (colored circles). Beige lines indicate developmental trajectories through this decision space, with lines branching upwards indicating the sampling of $C_{1}$ and lines branching downwards indicating the sampling of $C_{o}$. Colors denote the optimal, fitness-maximizing phenotypic choice in each state. Pies indicate cases in which organisms with the same posterior estimates make different phenotypic decisions. The area of a circle (pie piece) is proportional the probability of reaching that particular state. Colors indicate the following phenotypic decisions: Black corresponds to waiting, red to constructing $P_{0}$, blue to constructing $P_{1}$, purple to deconstructing $P_{0}$, green to deconstructing $P_{1}$, light red to a tie between constructing $P_{0}$ and deconstructing $P_{1}$, light blue to a tie between constructing $P_{1}$ and deconstructing $P_{0}$, brown to a tie between constructing either phenotypic target, yellow to a tie between deconstructing either target, grey to a tie between construction and waiting, dark grey to a tie between deconstruction and waiting, and lastly ochre to a tie between all options.

*Diminishing rewards and increasing penalties*

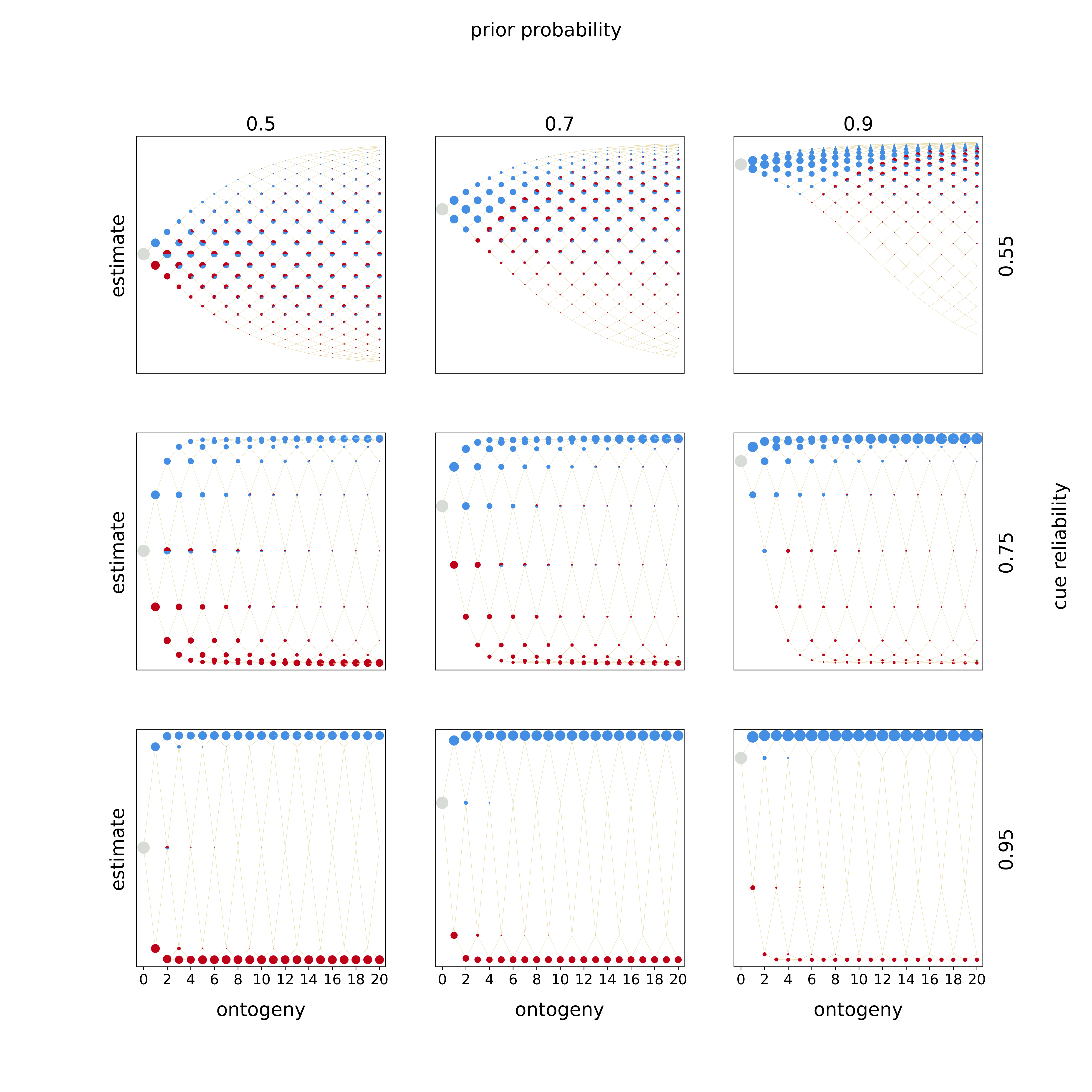

*Figure S2.17:* Optimal policies. Optimal policies are shown for a model with complete deconstruction and diminishing rewards and increasing penalties. Rows indicate the prior estimate of being in $E_{1}$ and columns indicate the cue reliability. Within each panel, the horizontal axis denotes ontogeny and the vertical axis the posterior estimates of being in $E_{1}$. The entire population starts ontogeny with zero cues sampled and the prior estimate indicated by the row (indicated by the grey circle). In each time period organisms sample a cue (either $C_{0}$ or $C_{1}$), update their estimate, and make a phenotypic decision (colored circles). Beige lines indicate developmental trajectories through this decision space, with lines branching upwards indicating the sampling of $C_{1}$ and lines branching downwards indicating the sampling of $C_{o}$. Colors denote the optimal, fitness-maximizing phenotypic choice in each state. Pies indicate cases in which organisms with the same posterior estimates make different phenotypic decisions. The area of a circle (pie piece) is proportional the probability of reaching that particular state. Colors indicate the following phenotypic decisions: Black corresponds to waiting, red to constructing $P_{0}$, blue to constructing $P_{1}$, purple to deconstructing $P_{0}$, green to deconstructing $P_{1}$, light red to a tie between constructing $P_{0}$ and deconstructing $P_{1}$, light blue to a tie between constructing $P_{1}$ and deconstructing $P_{0}$, brown to a tie between constructing either phenotypic target, yellow to a tie between deconstructing either target, grey to a tie between construction and waiting, dark grey to a tie between deconstruction and waiting, and lastly ochre to a tie between all options.

*Diminishing rewards and diminishing penalties*

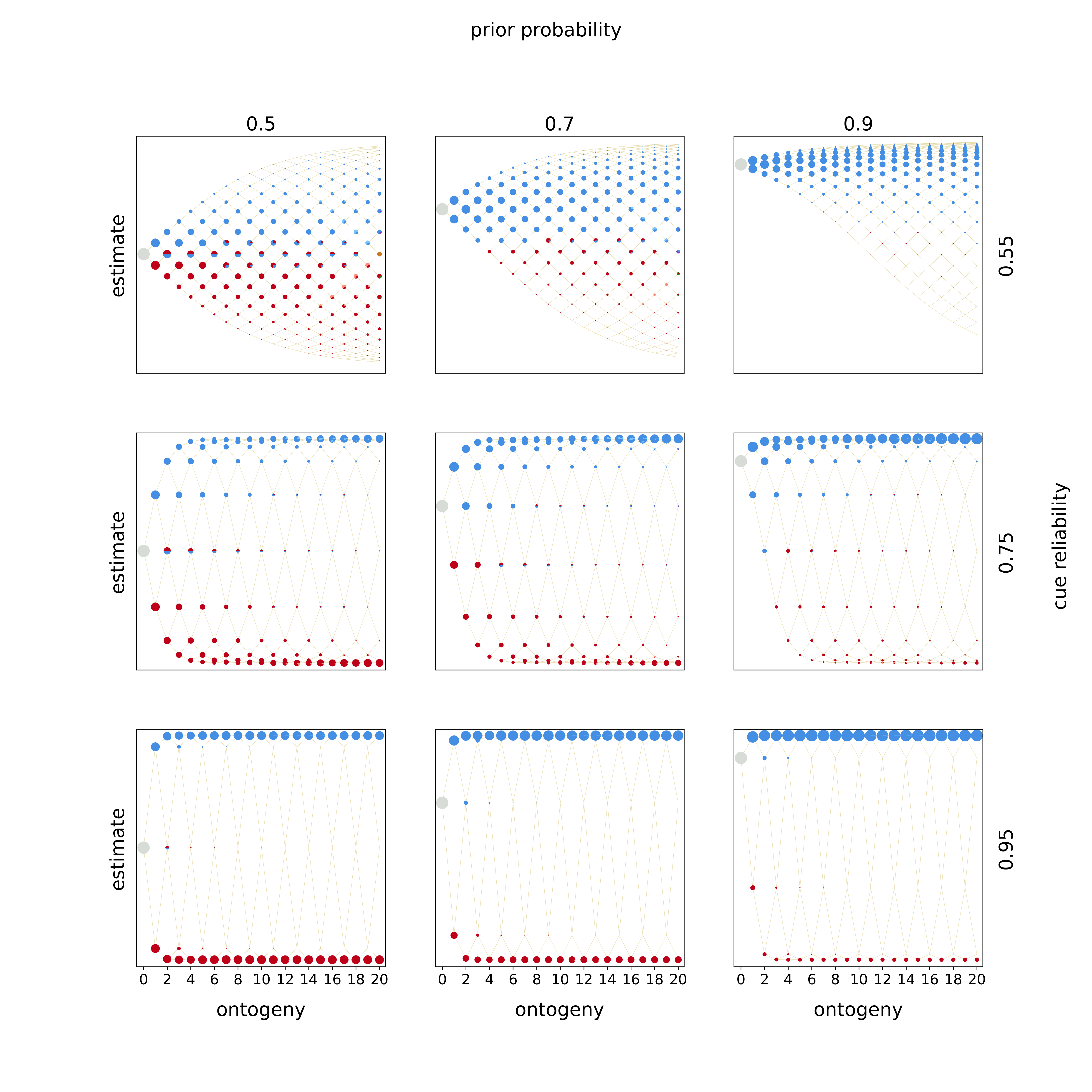

*Figure S2.18:* Optimal policies. Optimal policies are shown for a model with complete deconstruction and diminishing rewards and diminishing penalties. Rows indicate the prior estimate of being in $E_{1}$ and columns indicate the cue reliability. Within each panel, the horizontal axis denotes ontogeny and the vertical axis the posterior estimates of being in $E_{1}$. The entire population starts ontogeny with zero cues sampled and the prior estimate indicated by the row (indicated by the grey circle). In each time period organisms sample a cue (either $C_{0}$ or $C_{1}$), update their estimate, and make a phenotypic decision (colored circles). Beige lines indicate developmental trajectories through this decision space, with lines branching upwards indicating the sampling of $C_{1}$ and lines branching downwards indicating the sampling of $C_{o}$. Colors denote the optimal, fitness-maximizing phenotypic choice in each state. Pies indicate cases in which organisms with the same posterior estimates make different phenotypic decisions. The area of a circle (pie piece) is proportional the probability of reaching that particular state. Colors indicate the following phenotypic decisions: Black corresponds to waiting, red to constructing $P_{0}$, blue to constructing $P_{1}$, purple to deconstructing $P_{0}$, green to deconstructing $P_{1}$, light red to a tie between constructing $P_{0}$ and deconstructing $P_{1}$, light blue to a tie between constructing $P_{1}$ and deconstructing $P_{0}$, brown to a tie between constructing either phenotypic target, yellow to a tie between deconstructing either target, grey to a tie between construction and waiting, dark grey to a tie between deconstruction and waiting, and lastly ochre to a tie between all options.

**Distributions of mature phenotypes (incremental deconstruction)**

*Linear rewards and linear penalties*

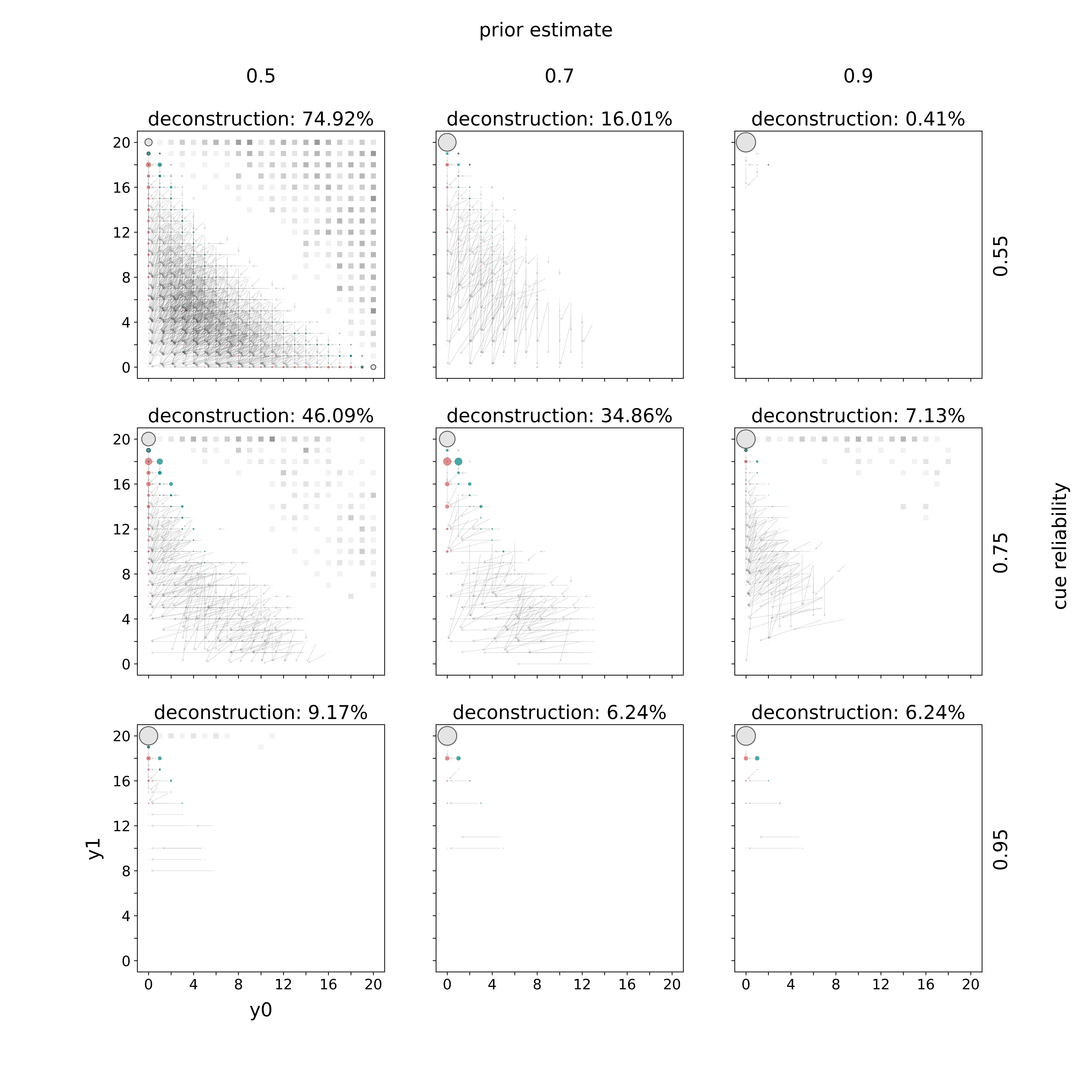

*Figure S2.19:* Distributions of mature phenotypes. Distributions of mature phenotypes are shown for a model with incremental deconstruction and linear rewards and penalties. Rows indicate the prior estimate of being in $E_{1}$ and columns indicate the cue reliability. The populations of mature phenotypes have been simulated in $E_{1}$. The title of each panel indicates the percentage of mature phenotypes that have deconstructed at some point during ontogeny. Within each panel the horizontal axis indicates the number of specializations towards $E_{0}$ and the vertical axis towards $E_{1}$. The lower triangle indicates how much mature phenotypes have constructed (teal circles) and what their phenotype looked like after deconstruction (red circles). Grey arrows connect phenotypes before (teal) and after (red) deconstruction. Grey circles with a black outline belong to mature phenotypes that never deconstructed. The area of a circle is proportional to the number mature organisms with this phenotype. The upper triangle indicates waiting. For each mature phenotype (after deconstruction) below the diagonal the corresponding square above the diagonal highlights the amount of waiting. The color intensity is proportional to the amount of waiting. Black squares indicate phenotypes that waited all of ontogeny (i.e. 20 time periods) and white squares phenotypes that never waited.

*Linear rewards and increasing penalties*

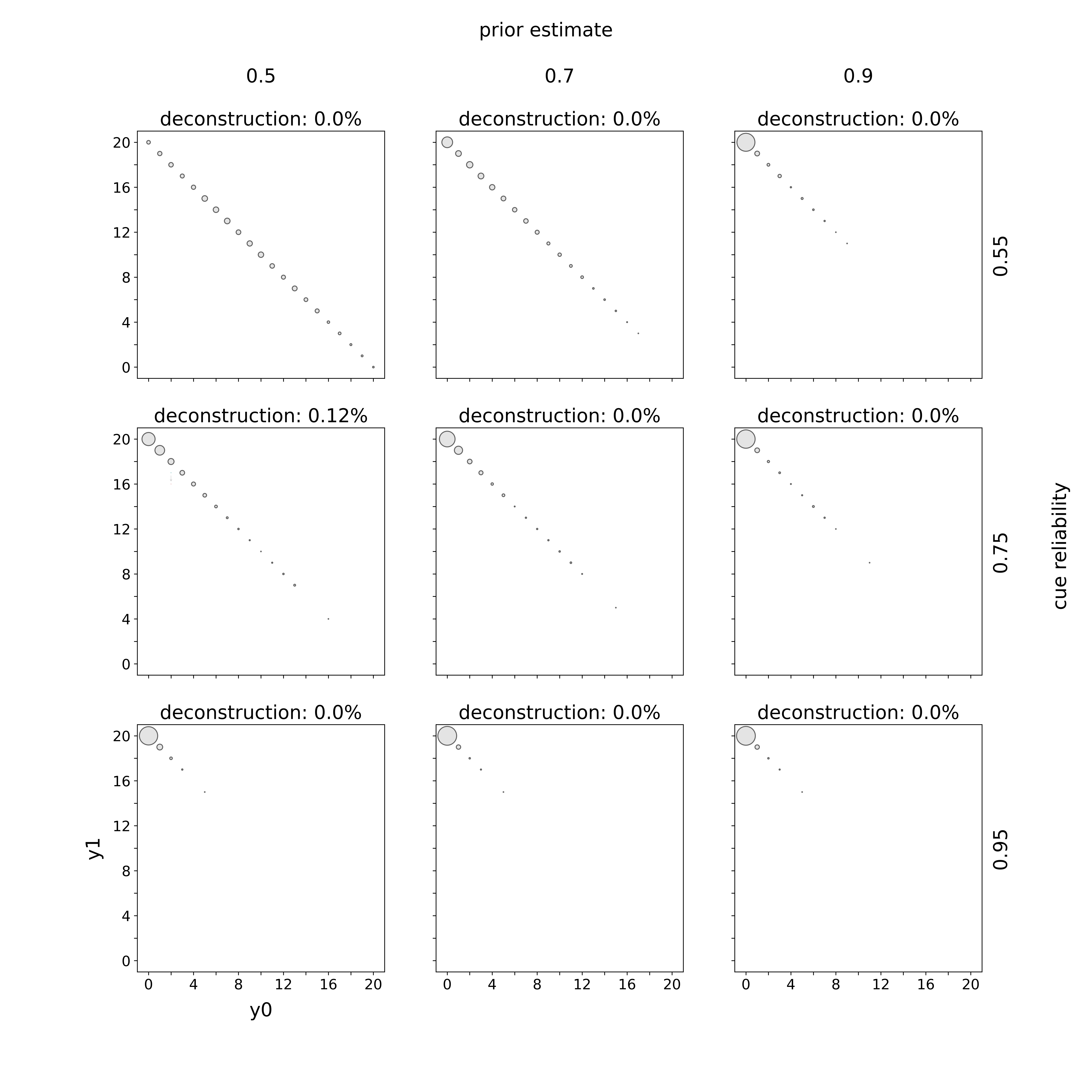

*Figure S2.20:* Distributions of mature phenotypes. Distributions of mature phenotypes are shown for a model with incremental deconstruction and linear rewards and increasing penalties. Rows indicate the prior estimate of being in $E_{1}$ and columns indicate the cue reliability. The populations of mature phenotypes have been simulated in $E_{1}$. The title of each panel indicates the percentage of mature phenotypes that have deconstructed at some point during ontogeny. Within each panel the horizontal axis indicates the number of specializations towards $E_{0}$ and the vertical axis towards $E_{1}$. The lower triangle indicates how much mature phenotypes have constructed (teal circles) and what their phenotype looked like after deconstruction (red circles). Grey arrows connect phenotypes before (teal) and after (red) deconstruction. Grey circles with a black outline belong to mature phenotypes that never deconstructed. The area of a circle is proportional to the number mature organisms with this phenotype. The upper triangle indicates waiting. For each mature phenotype (after deconstruction) below the diagonal the corresponding square above the diagonal highlights the amount of waiting. The color intensity is proportional to the amount of waiting. Black squares indicate phenotypes that waited all of ontogeny (i.e. 20 time periods) and white squares phenotypes that never waited.

*Linear rewards and diminishing penalties*

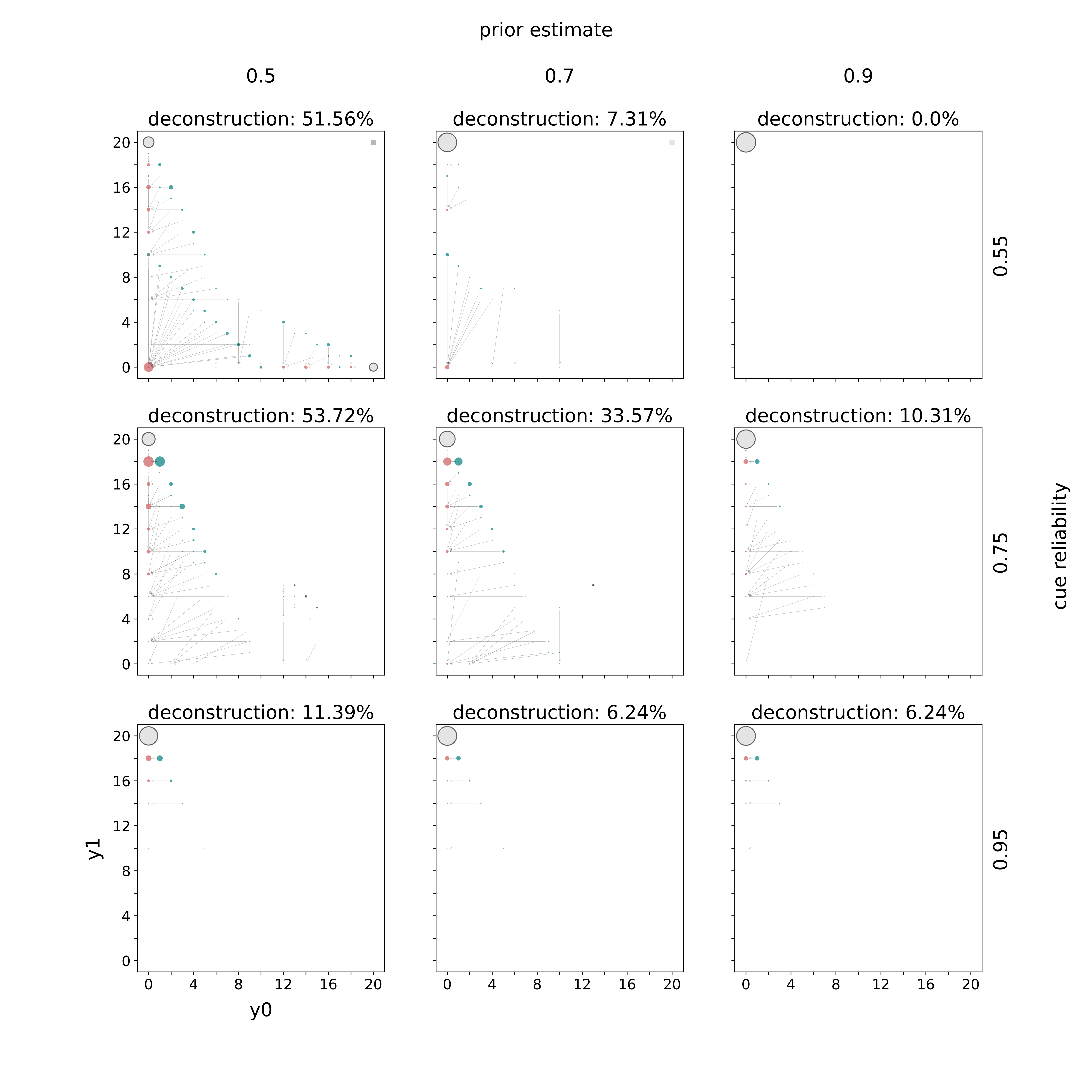

*Figure S2.21:* Distributions of mature phenotypes. Distributions of mature phenotypes are shown for a model with incremental deconstruction and linear rewards and diminishing penalties. Rows indicate the prior estimate of being in $E_{1}$ and columns indicate the cue reliability. The populations of mature phenotypes have been simulated in $E_{1}$. The title of each panel indicates the percentage of mature phenotypes that have deconstructed at some point during ontogeny. Within each panel the horizontal axis indicates the number of specializations towards $E_{0}$ and the vertical axis towards $E_{1}$. The lower triangle indicates how much mature phenotypes have constructed (teal circles) and what their phenotype looked like after deconstruction (red circles). Grey arrows connect phenotypes before (teal) and after (red) deconstruction. Grey circles with a black outline belong to mature phenotypes that never deconstructed. The area of a circle is proportional to the number mature organisms with this phenotype. The upper triangle indicates waiting. For each mature phenotype (after deconstruction) below the diagonal the corresponding square above the diagonal highlights the amount of waiting. The color intensity is proportional to the amount of waiting. Black squares indicate phenotypes that waited all of ontogeny (i.e. 20 time periods) and white squares phenotypes that never waited.

*Increasing rewards and linear penalties*

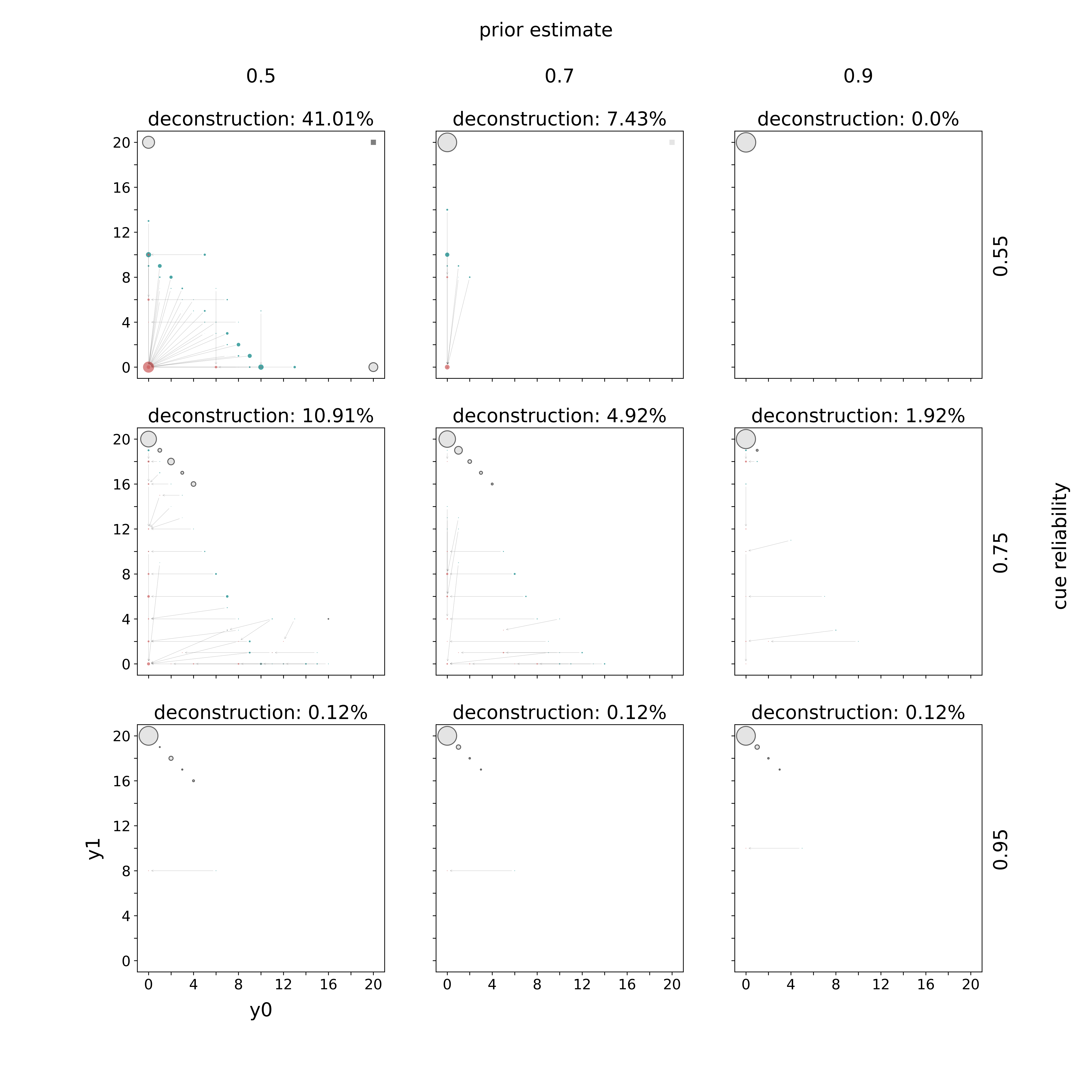

*Figure S2.22:* Distributions of mature phenotypes. Distributions of mature phenotypes are shown for a model with incremental deconstruction and increasing rewards and linear penalties. Rows indicate the prior estimate of being in $E_{1}$ and columns indicate the cue reliability. The populations of mature phenotypes have been simulated in $E_{1}$. The title of each panel indicates the percentage of mature phenotypes that have deconstructed at some point during ontogeny. Within each panel the horizontal axis indicates the number of specializations towards $E_{0}$ and the vertical axis towards $E_{1}$. The lower triangle indicates how much mature phenotypes have constructed (teal circles) and what their phenotype looked like after deconstruction (red circles). Grey arrows connect phenotypes before (teal) and after (red) deconstruction. Grey circles with a black outline belong to mature phenotypes that never deconstructed. The area of a circle is proportional to the number mature organisms with this phenotype. The upper triangle indicates waiting. For each mature phenotype (after deconstruction) below the diagonal the corresponding square above the diagonal highlights the amount of waiting. The color intensity is proportional to the amount of waiting. Black squares indicate phenotypes that waited all of ontogeny (i.e. 20 time periods) and white squares phenotypes that never waited.

*Increasing rewards and increasing penalties*

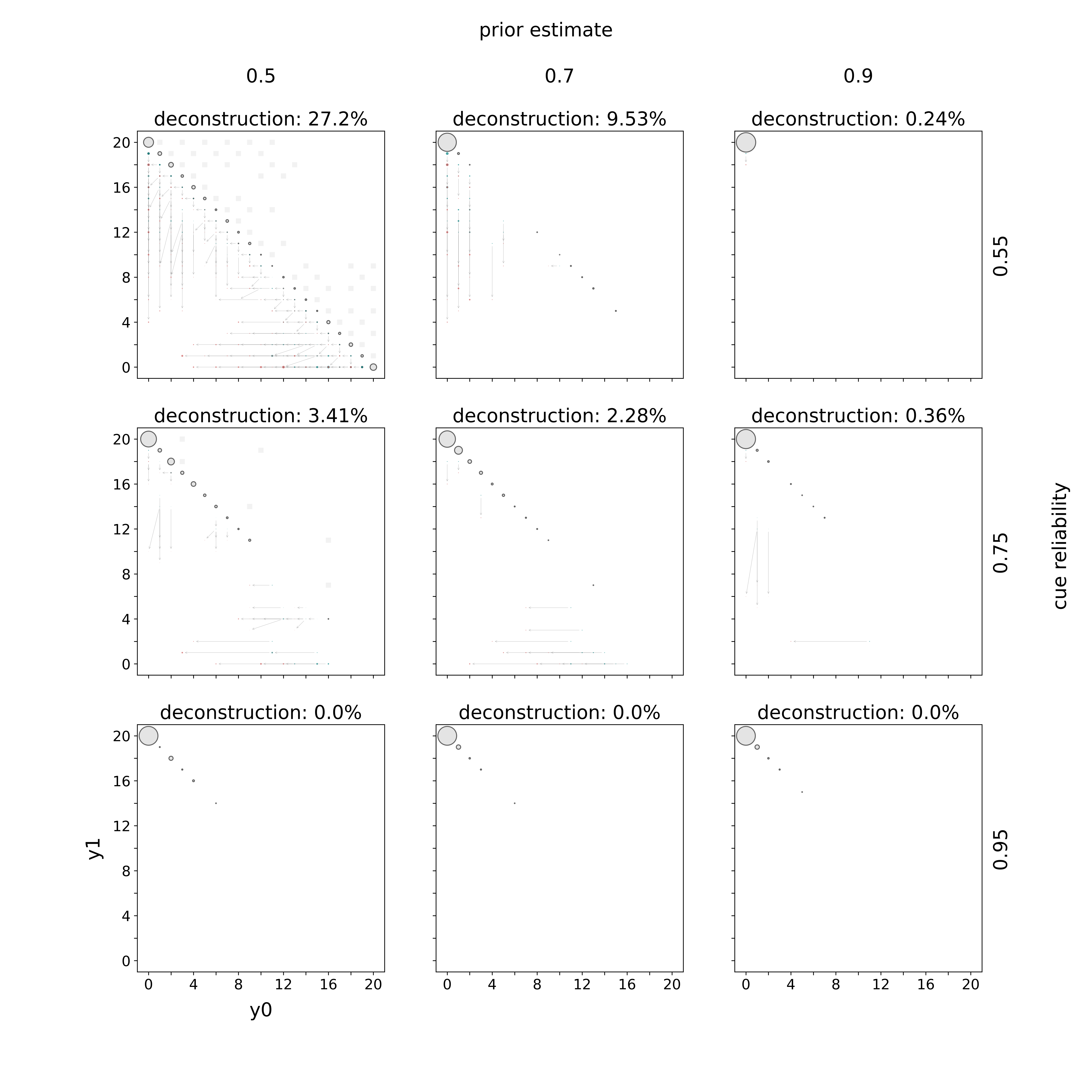

*Figure S2.23:* Distributions of mature phenotypes. Distributions of mature phenotypes are shown for a model with incremental deconstruction and increasing rewards and increasing penalties. Rows indicate the prior estimate of being in $E_{1}$ and columns indicate the cue reliability. The populations of mature phenotypes have been simulated in $E_{1}$. The title of each panel indicates the percentage of mature phenotypes that have deconstructed at some point during ontogeny. Within each panel the horizontal axis indicates the number of specializations towards $E_{0}$ and the vertical axis towards $E_{1}$. The lower triangle indicates how much mature phenotypes have constructed (teal circles) and what their phenotype looked like after deconstruction (red circles). Grey arrows connect phenotypes before (teal) and after (red) deconstruction. Grey circles with a black outline belong to mature phenotypes that never deconstructed. The area of a circle is proportional to the number mature organisms with this phenotype. The upper triangle indicates waiting. For each mature phenotype (after deconstruction) below the diagonal the corresponding square above the diagonal highlights the amount of waiting. The color intensity is proportional to the amount of waiting. Black squares indicate phenotypes that waited all of ontogeny (i.e. 20 time periods) and white squares phenotypes that never waited.

*Increasing rewards and diminishing penalties*

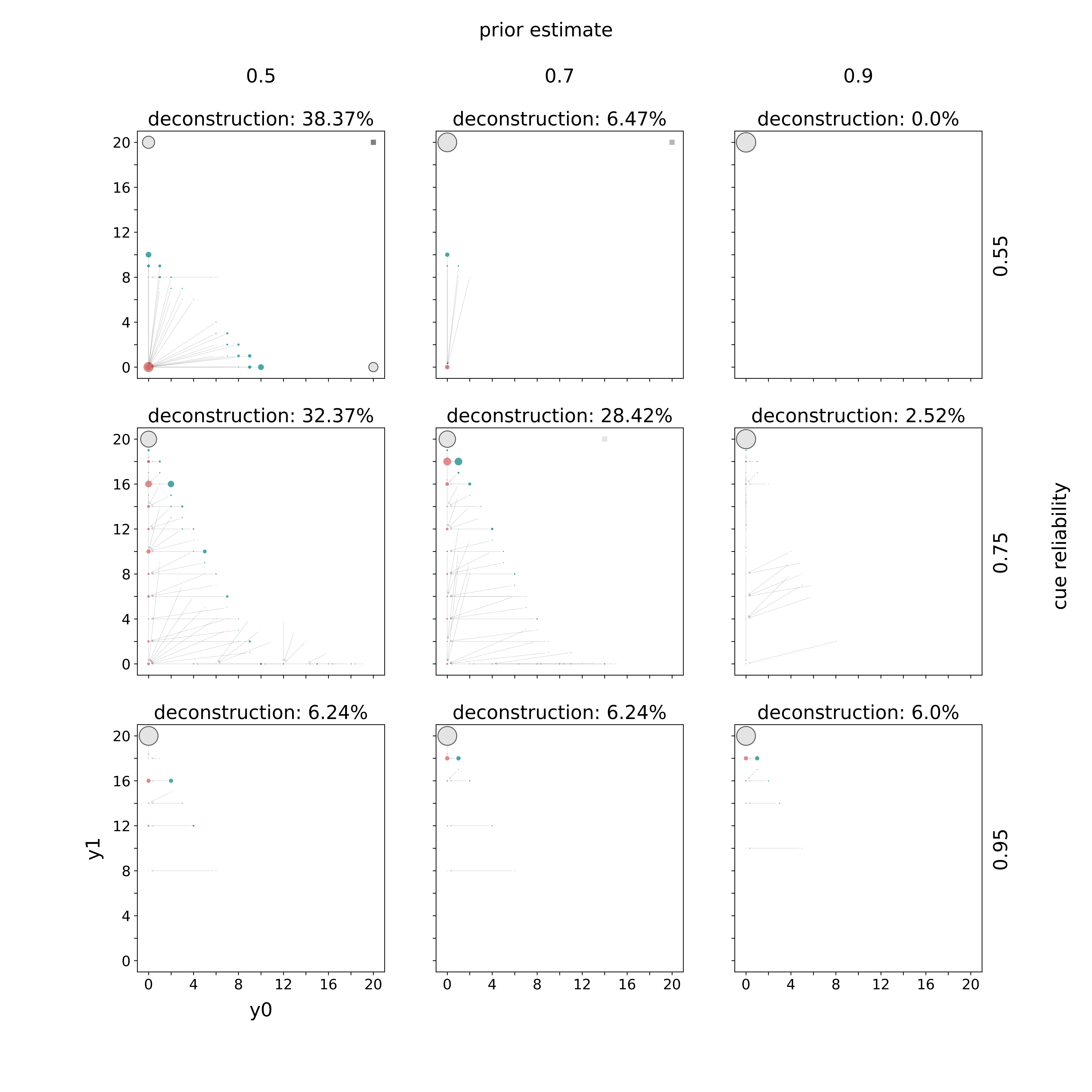

*Figure S2.24:* Distributions of mature phenotypes. Distributions of mature phenotypes are shown for a model with incremental deconstruction and increasing rewards and diminishing penalties. Rows indicate the prior estimate of being in $E_{1}$ and columns indicate the cue reliability. The populations of mature phenotypes have been simulated in $E_{1}$. The title of each panel indicates the percentage of mature phenotypes that have deconstructed at some point during ontogeny. Within each panel the horizontal axis indicates the number of specializations towards $E_{0}$ and the vertical axis towards $E_{1}$. The lower triangle indicates how much mature phenotypes have constructed (teal circles) and what their phenotype looked like after deconstruction (red circles). Grey arrows connect phenotypes before (teal) and after (red) deconstruction. Grey circles with a black outline belong to mature phenotypes that never deconstructed. The area of a circle is proportional to the number mature organisms with this phenotype. The upper triangle indicates waiting. For each mature phenotype (after deconstruction) below the diagonal the corresponding square above the diagonal highlights the amount of waiting. The color intensity is proportional to the amount of waiting. Black squares indicate phenotypes that waited all of ontogeny (i.e. 20 time periods) and white squares phenotypes that never waited.

*Diminishing rewards and linear penalties*

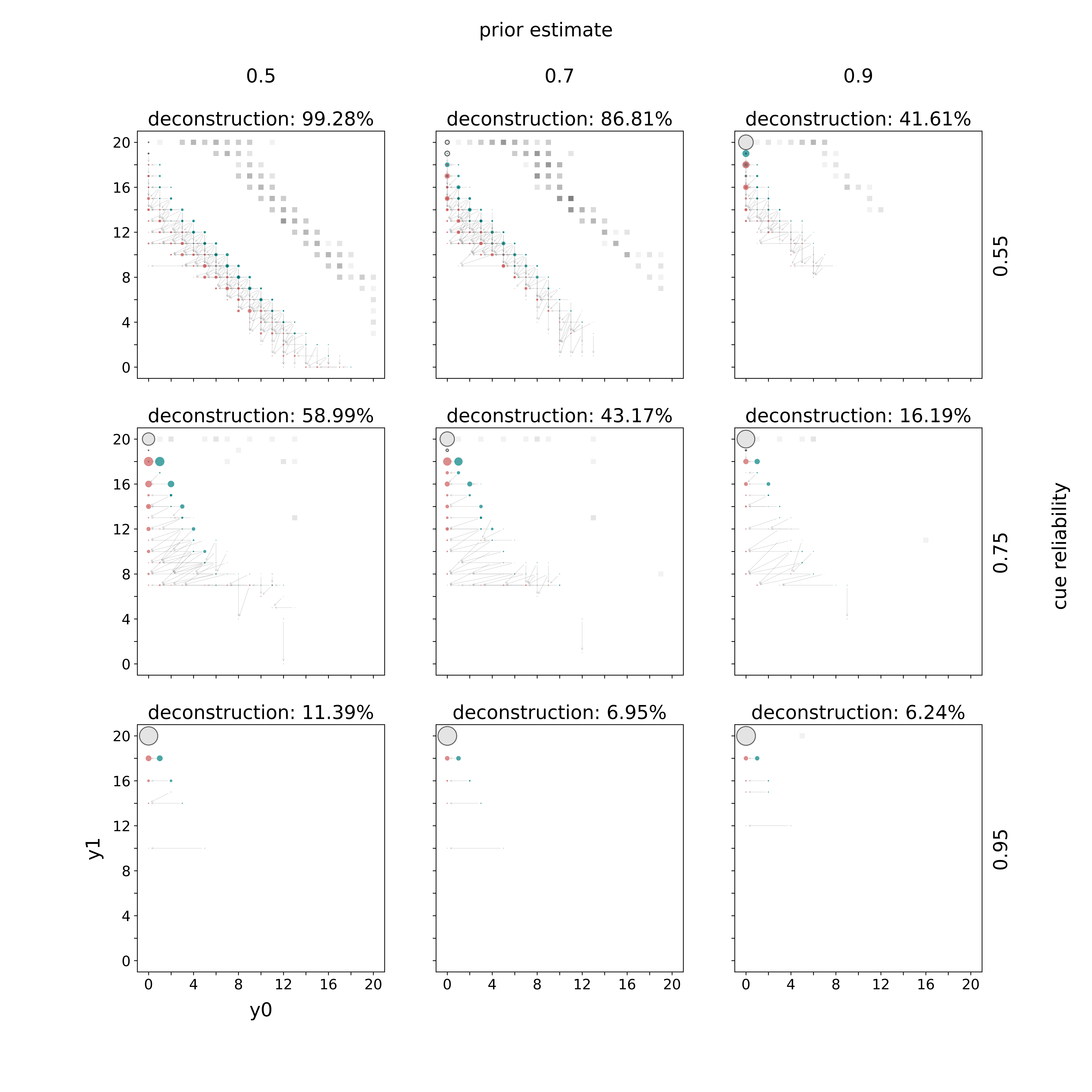

*Figure S2.25:* Distributions of mature phenotypes. Distributions of mature phenotypes are shown for a model with incremental deconstruction and diminishing rewards and linear penalties. Rows indicate the prior estimate of being in $E_{1}$ and columns indicate the cue reliability. The populations of mature phenotypes have been simulated in $E_{1}$. The title of each panel indicates the percentage of mature phenotypes that have deconstructed at some point during ontogeny. Within each panel the horizontal axis indicates the number of specializations towards $E_{0}$ and the vertical axis towards $E_{1}$. The lower triangle indicates how much mature phenotypes have constructed (teal circles) and what their phenotype looked like after deconstruction (red circles). Grey arrows connect phenotypes before (teal) and after (red) deconstruction. Grey circles with a black outline belong to mature phenotypes that never deconstructed. The area of a circle is proportional to the number mature organisms with this phenotype. The upper triangle indicates waiting. For each mature phenotype (after deconstruction) below the diagonal the corresponding square above the diagonal highlights the amount of waiting. The color intensity is proportional to the amount of waiting. Black squares indicate phenotypes that waited all of ontogeny (i.e. 20 time periods) and white squares phenotypes that never waited.

*Diminishing rewards and increasing penalties*

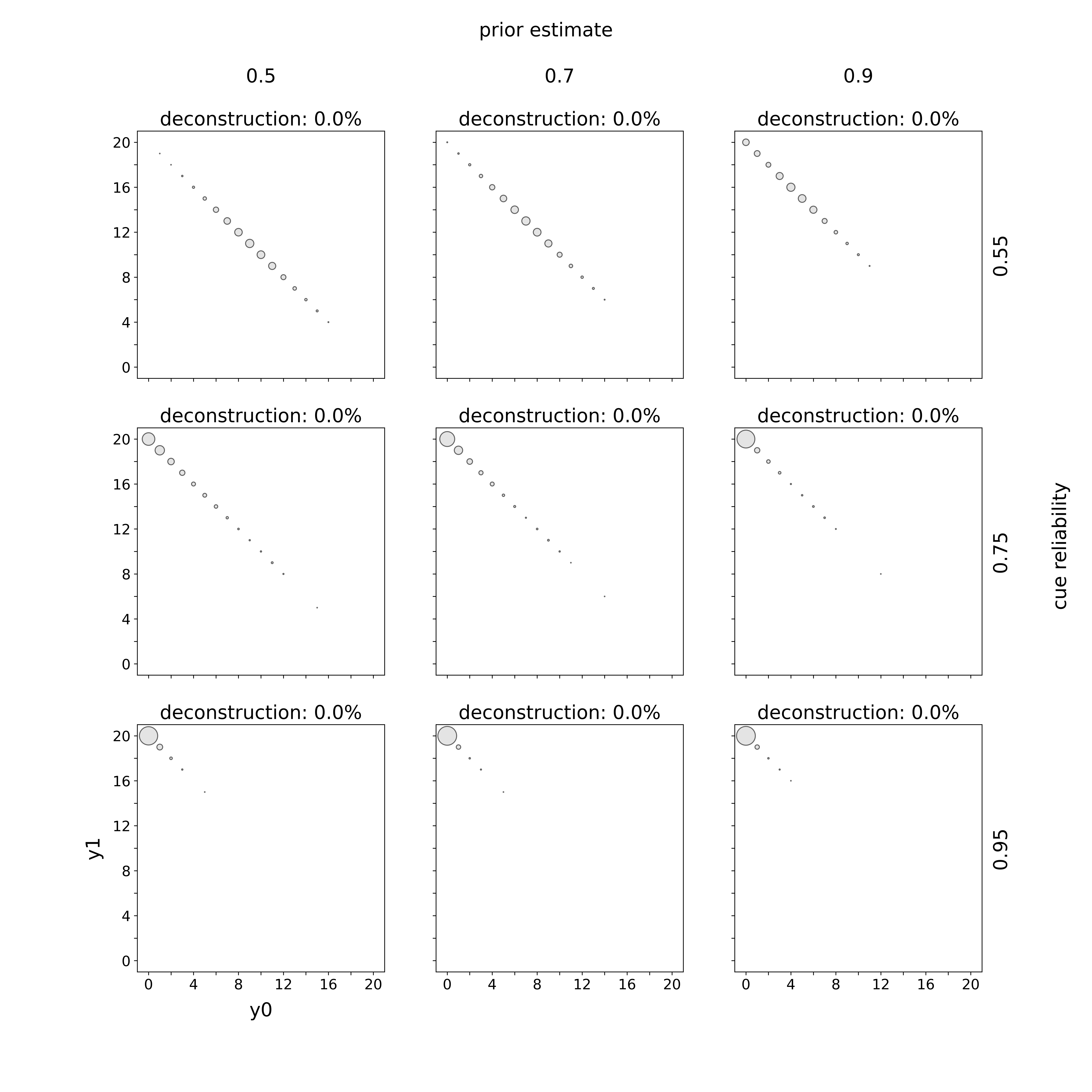

*Figure S2.26:* Distributions of mature phenotypes. Distributions of mature phenotypes are shown for a model with incremental deconstruction and diminishing rewards and increasing penalties. Rows indicate the prior estimate of being in $E_{1}$ and columns indicate the cue reliability. The populations of mature phenotypes have been simulated in $E_{1}$. The title of each panel indicates the percentage of mature phenotypes that have deconstructed at some point during ontogeny. Within each panel the horizontal axis indicates the number of specializations towards $E_{0}$ and the vertical axis towards $E_{1}$. The lower triangle indicates how much mature phenotypes have constructed (teal circles) and what their phenotype looked like after deconstruction (red circles). Grey arrows connect phenotypes before (teal) and after (red) deconstruction. Grey circles with a black outline belong to mature phenotypes that never deconstructed. The area of a circle is proportional to the number mature organisms with this phenotype. The upper triangle indicates waiting. For each mature phenotype (after deconstruction) below the diagonal the corresponding square above the diagonal highlights the amount of waiting. The color intensity is proportional to the amount of waiting. Black squares indicate phenotypes that waited all of ontogeny (i.e. 20 time periods) and white squares phenotypes that never waited.

*Diminishing rewards and diminishing penalties*

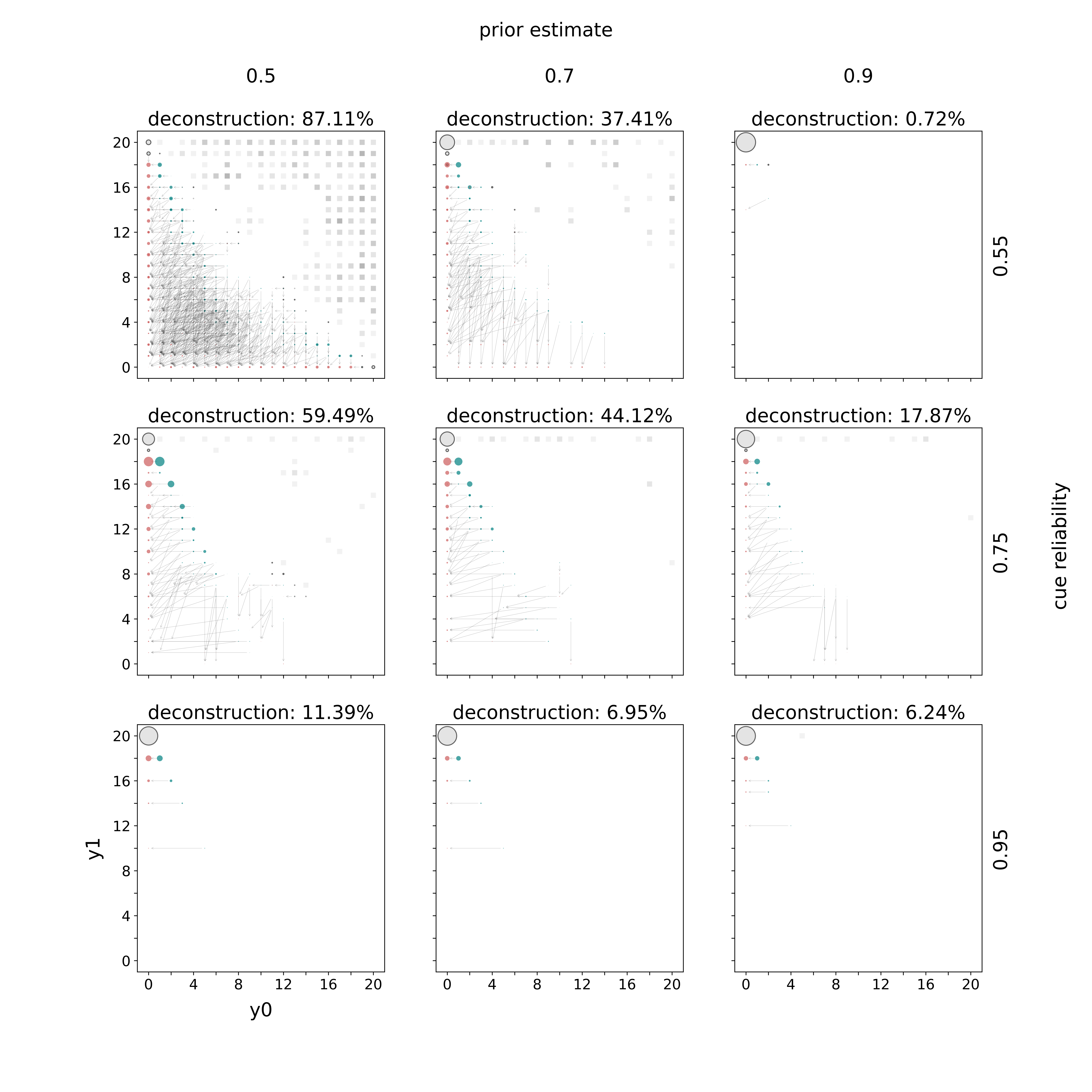

*Figure S2.27:* Distributions of mature phenotypes. Distributions of mature phenotypes are shown for a model with incremental deconstruction and diminishing rewards and diminishing penalties. Rows indicate the prior estimate of being in $E_{1}$ and columns indicate the cue reliability. The populations of mature phenotypes have been simulated in $E_{1}$. The title of each panel indicates the percentage of mature phenotypes that have deconstructed at some point during ontogeny. Within each panel the horizontal axis indicates the number of specializations towards $E_{0}$ and the vertical axis towards $E_{1}$. The lower triangle indicates how much mature phenotypes have constructed (teal circles) and what their phenotype looked like after deconstruction (red circles). Grey arrows connect phenotypes before (teal) and after (red) deconstruction. Grey circles with a black outline belong to mature phenotypes that never deconstructed. The area of a circle is proportional to the number mature organisms with this phenotype. The upper triangle indicates waiting. For each mature phenotype (after deconstruction) below the diagonal the corresponding square above the diagonal highlights the amount of waiting. The color intensity is proportional to the amount of waiting. Black squares indicate phenotypes that waited all of ontogeny (i.e. 20 time periods) and white squares phenotypes that never waited.

**Distributions of mature phenotypes (complete deconstruction)**

*Linear rewards and linear penalties*

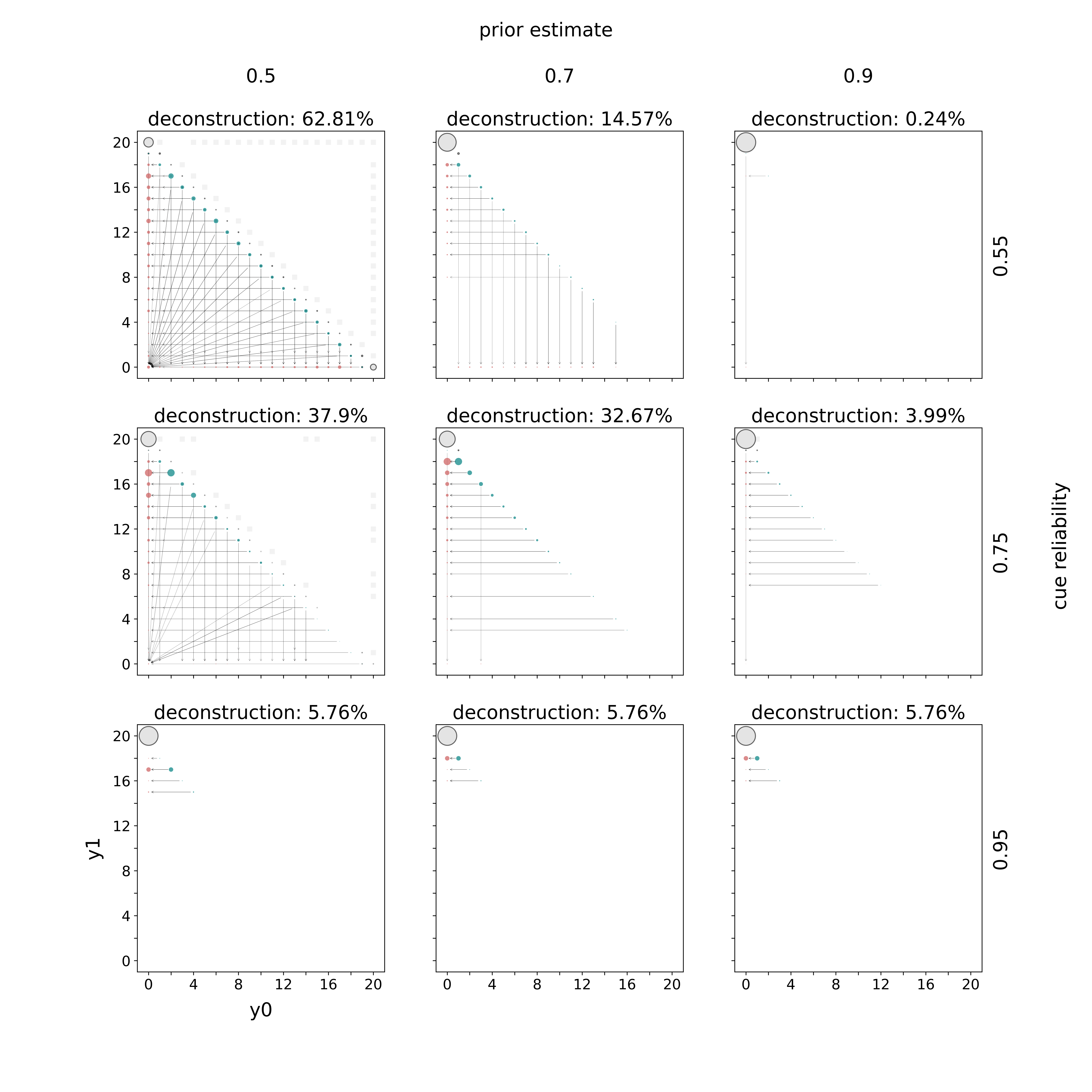

*Figure S2.28:* Distributions of mature phenotypes. Distributions of mature phenotypes are shown for a model with complete deconstruction and linear rewards and penalties. Rows indicate the prior estimate of being in $E_{1}$ and columns indicate the cue reliability. The populations of mature phenotypes have been simulated in $E_{1}$. The title of each panel indicates the percentage of mature phenotypes that have deconstructed at some point during ontogeny. Within each panel the horizontal axis indicates the number of specializations towards $E_{0}$ and the vertical axis towards $E_{1}$. The lower triangle indicates how much mature phenotypes have constructed (teal circles) and what their phenotype looked like after deconstruction (red circles). Grey arrows connect phenotypes before (teal) and after (red) deconstruction. Grey circles with a black outline belong to mature phenotypes that never deconstructed. The area of a circle is proportional to the number mature organisms with this phenotype. The upper triangle indicates waiting. For each mature phenotype (after deconstruction) below the diagonal the corresponding square above the diagonal highlights the amount of waiting. The color intensity is proportional to the amount of waiting. Black squares indicate phenotypes that waited all of ontogeny (i.e. 20 time periods) and white squares phenotypes that never waited.

*Linear rewards and increasing penalties*

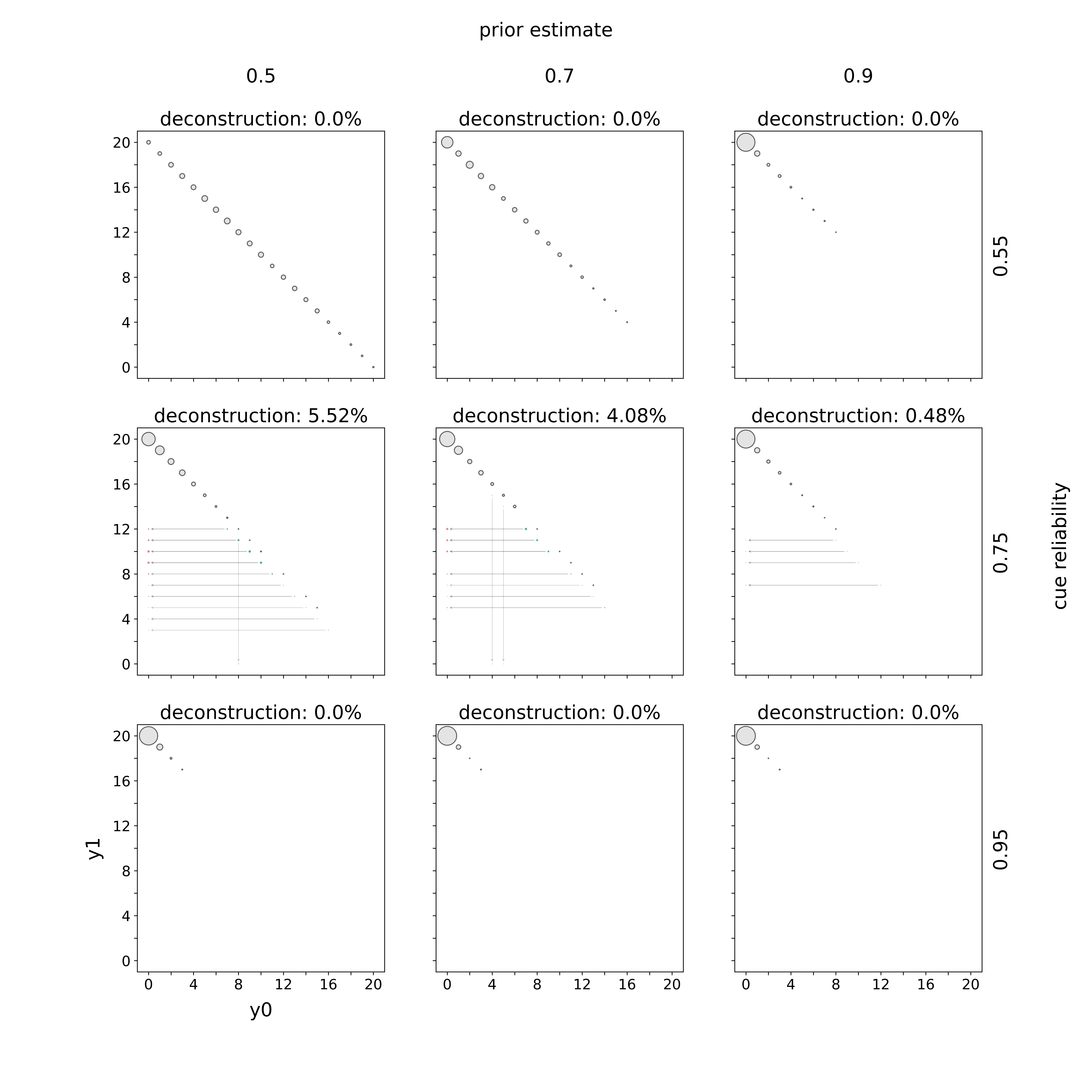

*Figure S2.29:* Distributions of mature phenotypes. Distributions of mature phenotypes are shown for a model with complete deconstruction and linear rewards and increasing penalties. Rows indicate the prior estimate of being in $E_{1}$ and columns indicate the cue reliability. The populations of mature phenotypes have been simulated in $E_{1}$. The title of each panel indicates the percentage of mature phenotypes that have deconstructed at some point during ontogeny. Within each panel the horizontal axis indicates the number of specializations towards $E_{0}$ and the vertical axis towards $E_{1}$. The lower triangle indicates how much mature phenotypes have constructed (teal circles) and what their phenotype looked like after deconstruction (red circles). Grey arrows connect phenotypes before (teal) and after (red) deconstruction. Grey circles with a black outline belong to mature phenotypes that never deconstructed. The area of a circle is proportional to the number mature organisms with this phenotype. The upper triangle indicates waiting. For each mature phenotype (after deconstruction) below the diagonal the corresponding square above the diagonal highlights the amount of waiting. The color intensity is proportional to the amount of waiting. Black squares indicate phenotypes that waited all of ontogeny (i.e. 20 time periods) and white squares phenotypes that never waited.

*Linear rewards and diminishing penalties*

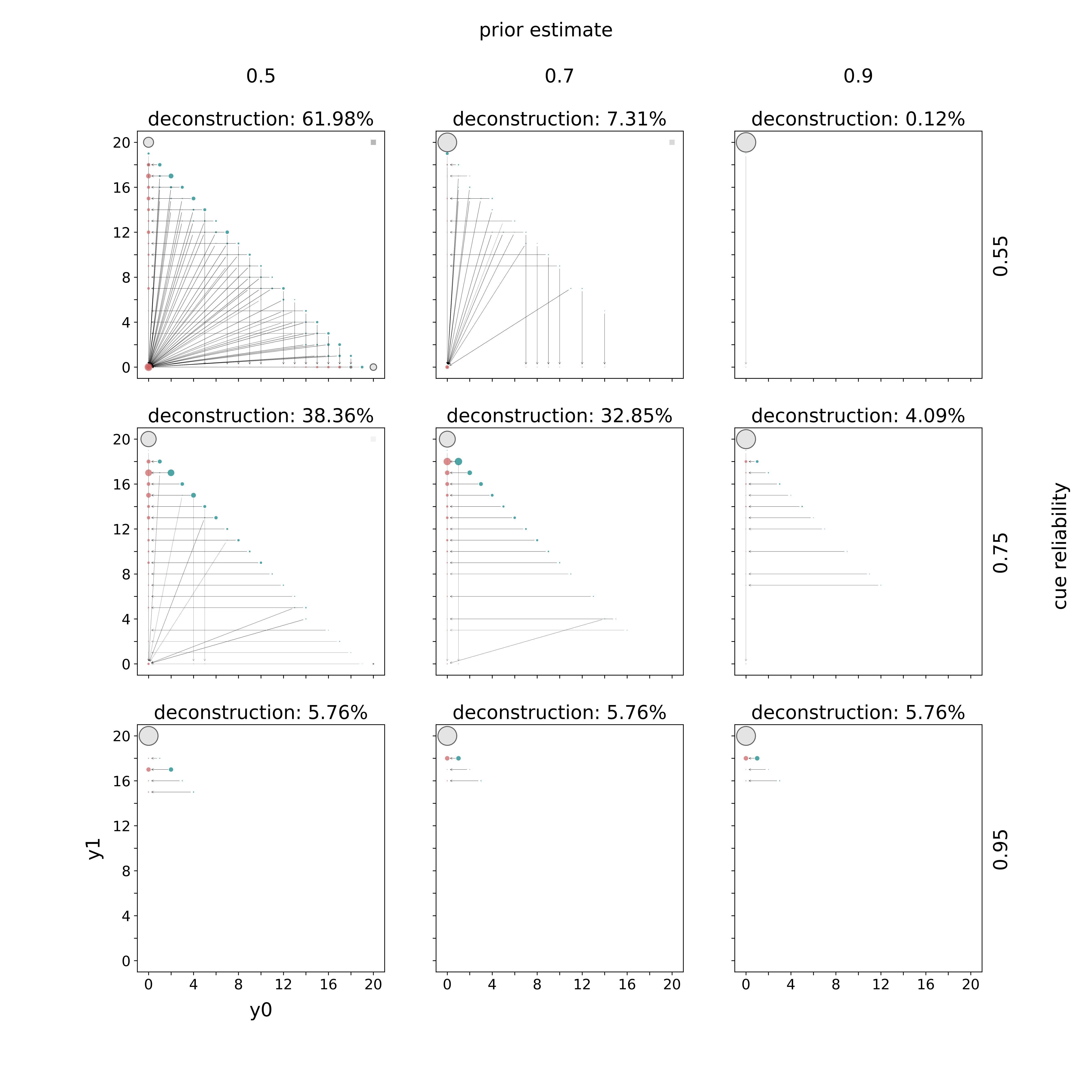

*Figure S2.30:* Distributions of mature phenotypes. Distributions of mature phenotypes are shown for a model with complete deconstruction and linear rewards and diminishing penalties. Rows indicate the prior estimate of being in $E_{1}$ and columns indicate the cue reliability. The populations of mature phenotypes have been simulated in $E_{1}$. The title of each panel indicates the percentage of mature phenotypes that have deconstructed at some point during ontogeny. Within each panel the horizontal axis indicates the number of specializations towards $E_{0}$ and the vertical axis towards $E_{1}$. The lower triangle indicates how much mature phenotypes have constructed (teal circles) and what their phenotype looked like after deconstruction (red circles). Grey arrows connect phenotypes before (teal) and after (red) deconstruction. Grey circles with a black outline belong to mature phenotypes that never deconstructed. The area of a circle is proportional to the number mature organisms with this phenotype. The upper triangle indicates waiting. For each mature phenotype (after deconstruction) below the diagonal the corresponding square above the diagonal highlights the amount of waiting. The color intensity is proportional to the amount of waiting. Black squares indicate phenotypes that waited all of ontogeny (i.e. 20 time periods) and white squares phenotypes that never waited.

*Increasing rewards and linear penalties*

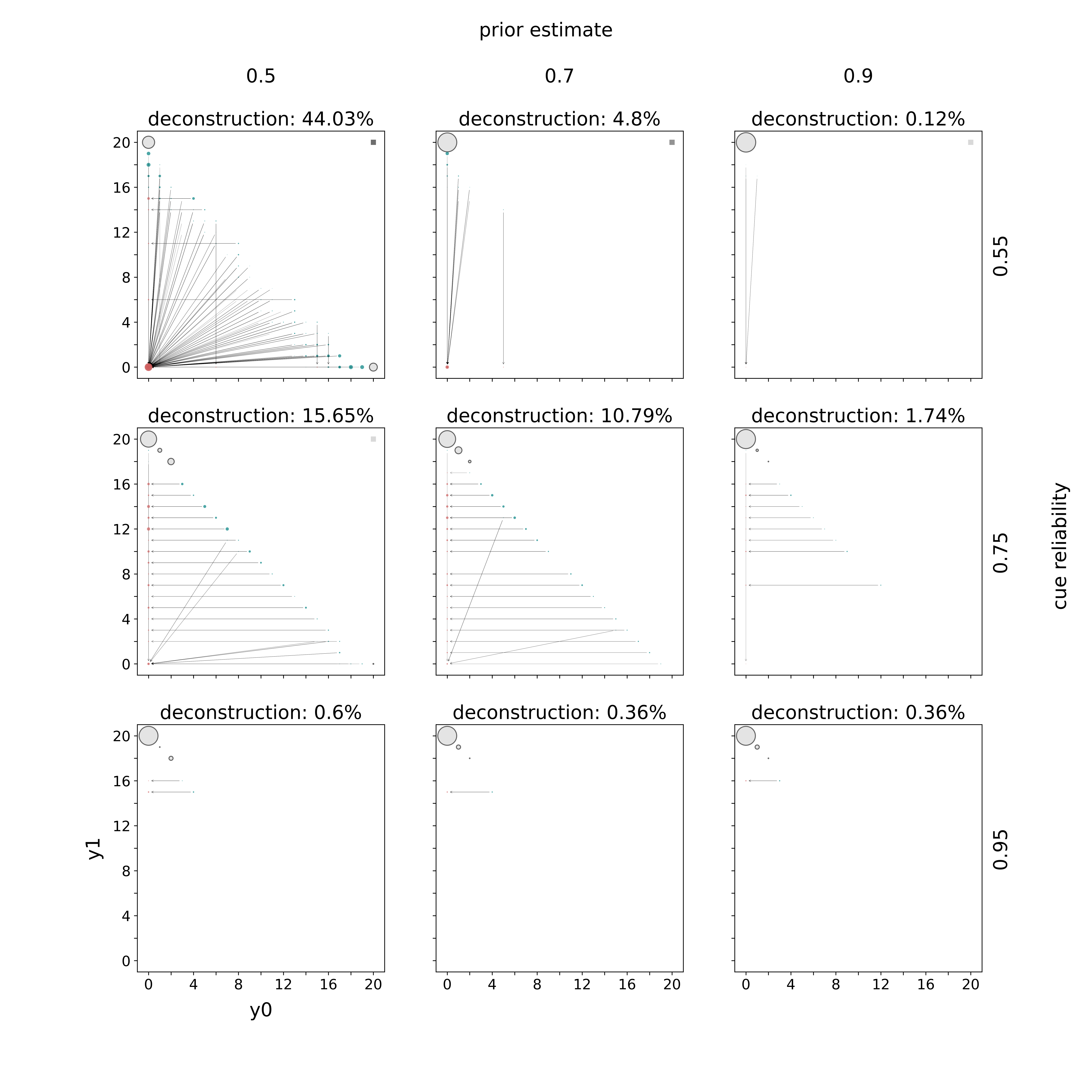

*Figure S2.31:* Distributions of mature phenotypes. Distributions of mature phenotypes are shown for a model with complete deconstruction and increasing rewards and linear penalties. Rows indicate the prior estimate of being in $E_{1}$ and columns indicate the cue reliability. The populations of mature phenotypes have been simulated in $E_{1}$. The title of each panel indicates the percentage of mature phenotypes that have deconstructed at some point during ontogeny. Within each panel the horizontal axis indicates the number of specializations towards $E_{0}$ and the vertical axis towards $E_{1}$. The lower triangle indicates how much mature phenotypes have constructed (teal circles) and what their phenotype looked like after deconstruction (red circles). Grey arrows connect phenotypes before (teal) and after (red) deconstruction. Grey circles with a black outline belong to mature phenotypes that never deconstructed. The area of a circle is proportional to the number mature organisms with this phenotype. The upper triangle indicates waiting. For each mature phenotype (after deconstruction) below the diagonal the corresponding square above the diagonal highlights the amount of waiting. The color intensity is proportional to the amount of waiting. Black squares indicate phenotypes that waited all of ontogeny (i.e. 20 time periods) and white squares phenotypes that never waited.

*Increasing rewards and increasing penalties*

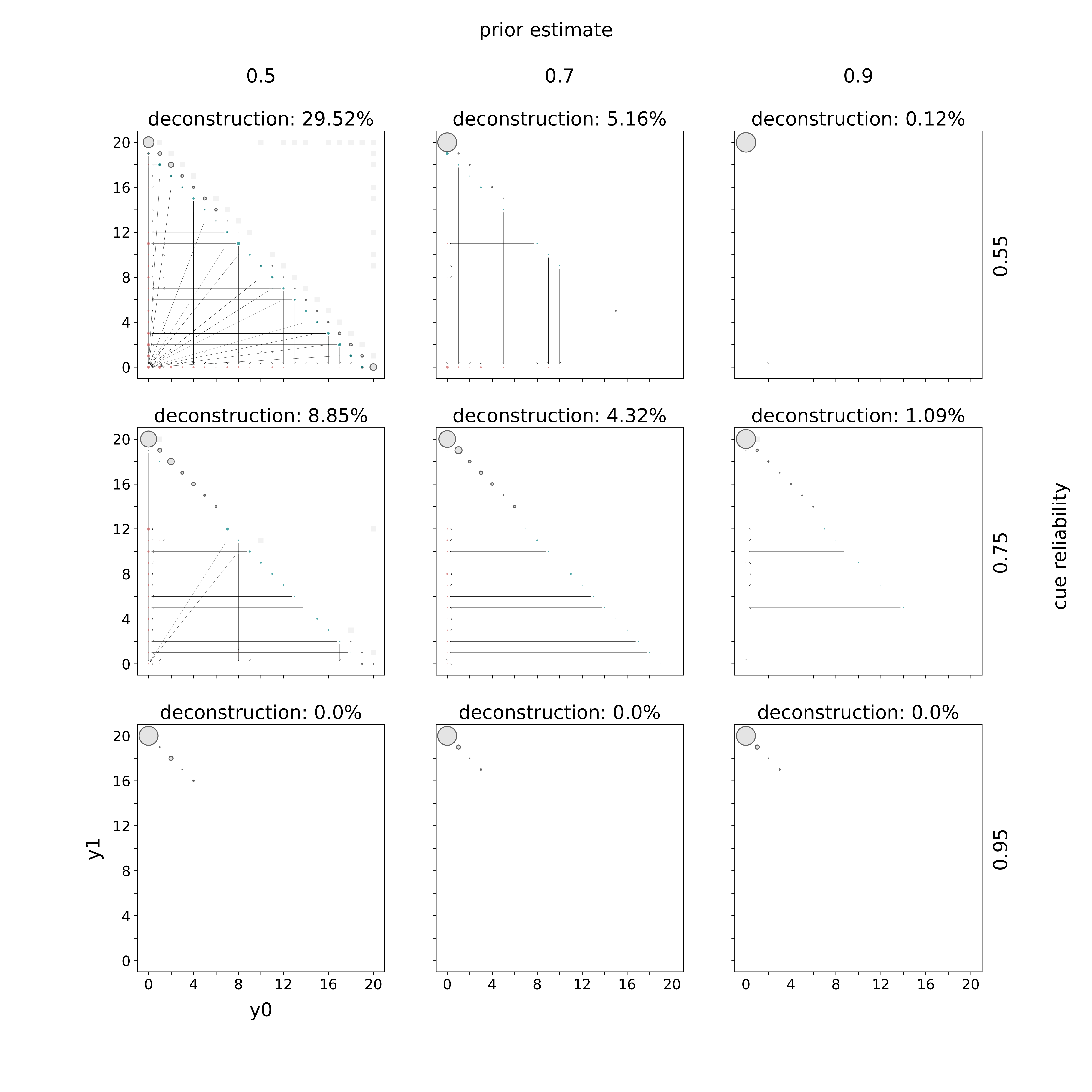

*Figure S2.32:* Distributions of mature phenotypes. Distributions of mature phenotypes are shown for a model with complete deconstruction and increasing rewards and increasing penalties. Rows indicate the prior estimate of being in $E_{1}$ and columns indicate the cue reliability. The populations of mature phenotypes have been simulated in $E_{1}$. The title of each panel indicates the percentage of mature phenotypes that have deconstructed at some point during ontogeny. Within each panel the horizontal axis indicates the number of specializations towards $E_{0}$ and the vertical axis towards $E_{1}$. The lower triangle indicates how much mature phenotypes have constructed (teal circles) and what their phenotype looked like after deconstruction (red circles). Grey arrows connect phenotypes before (teal) and after (red) deconstruction. Grey circles with a black outline belong to mature phenotypes that never deconstructed. The area of a circle is proportional to the number mature organisms with this phenotype. The upper triangle indicates waiting. For each mature phenotype (after deconstruction) below the diagonal the corresponding square above the diagonal highlights the amount of waiting. The color intensity is proportional to the amount of waiting. Black squares indicate phenotypes that waited all of ontogeny (i.e. 20 time periods) and white squares phenotypes that never waited.

*Increasing rewards and diminishing penalties*

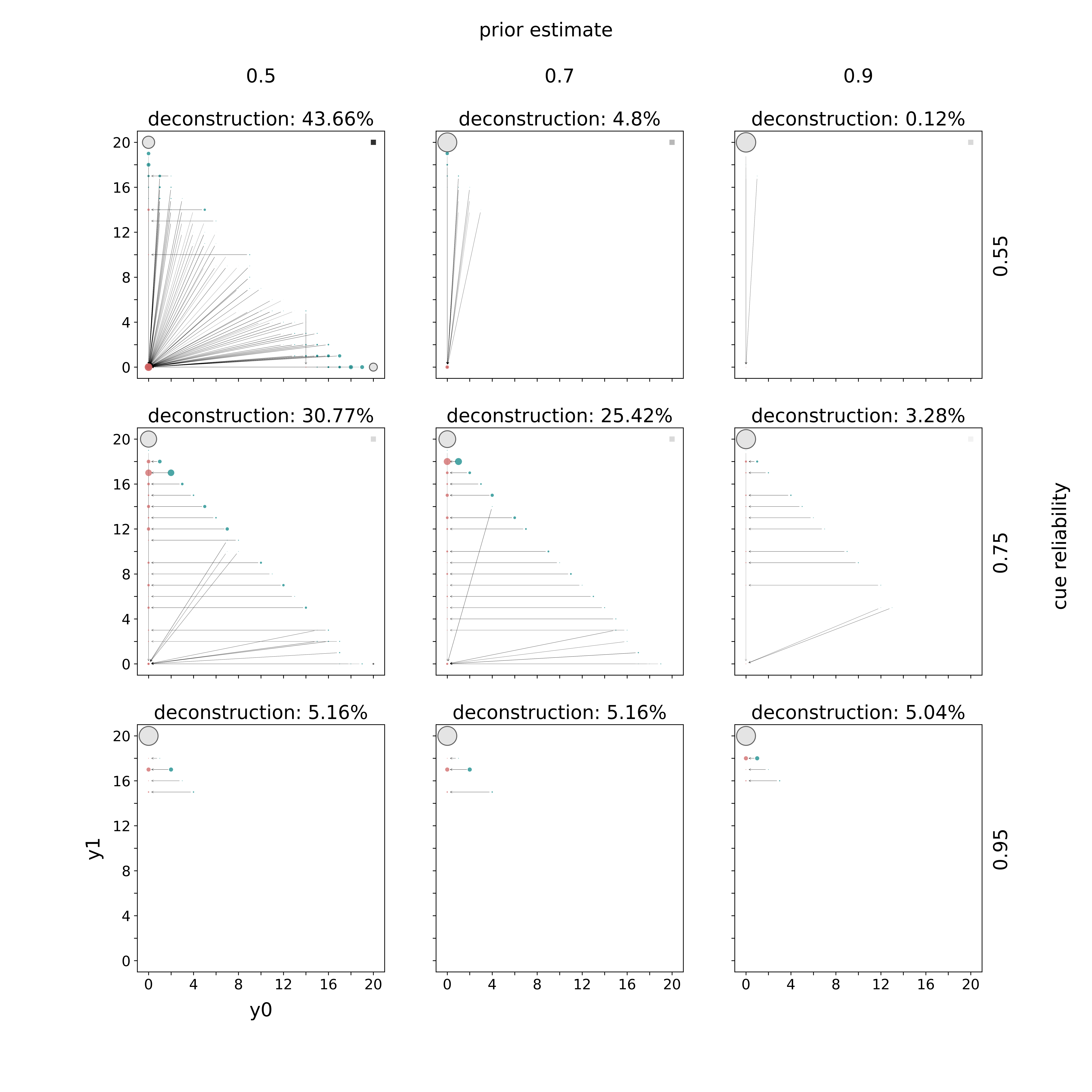

*Figure S2.33:* Distributions of mature phenotypes. Distributions of mature phenotypes are shown for a model with complete deconstruction and increasing rewards and diminishing penalties. Rows indicate the prior estimate of being in $E_{1}$ and columns indicate the cue reliability. The populations of mature phenotypes have been simulated in $E_{1}$. The title of each panel indicates the percentage of mature phenotypes that have deconstructed at some point during ontogeny. Within each panel the horizontal axis indicates the number of specializations towards $E_{0}$ and the vertical axis towards $E_{1}$. The lower triangle indicates how much mature phenotypes have constructed (teal circles) and what their phenotype looked like after deconstruction (red circles). Grey arrows connect phenotypes before (teal) and after (red) deconstruction. Grey circles with a black outline belong to mature phenotypes that never deconstructed. The area of a circle is proportional to the number mature organisms with this phenotype. The upper triangle indicates waiting. For each mature phenotype (after deconstruction) below the diagonal the corresponding square above the diagonal highlights the amount of waiting. The color intensity is proportional to the amount of waiting. Black squares indicate phenotypes that waited all of ontogeny (i.e. 20 time periods) and white squares phenotypes that never waited.

*Diminishing rewards and linear penalties*

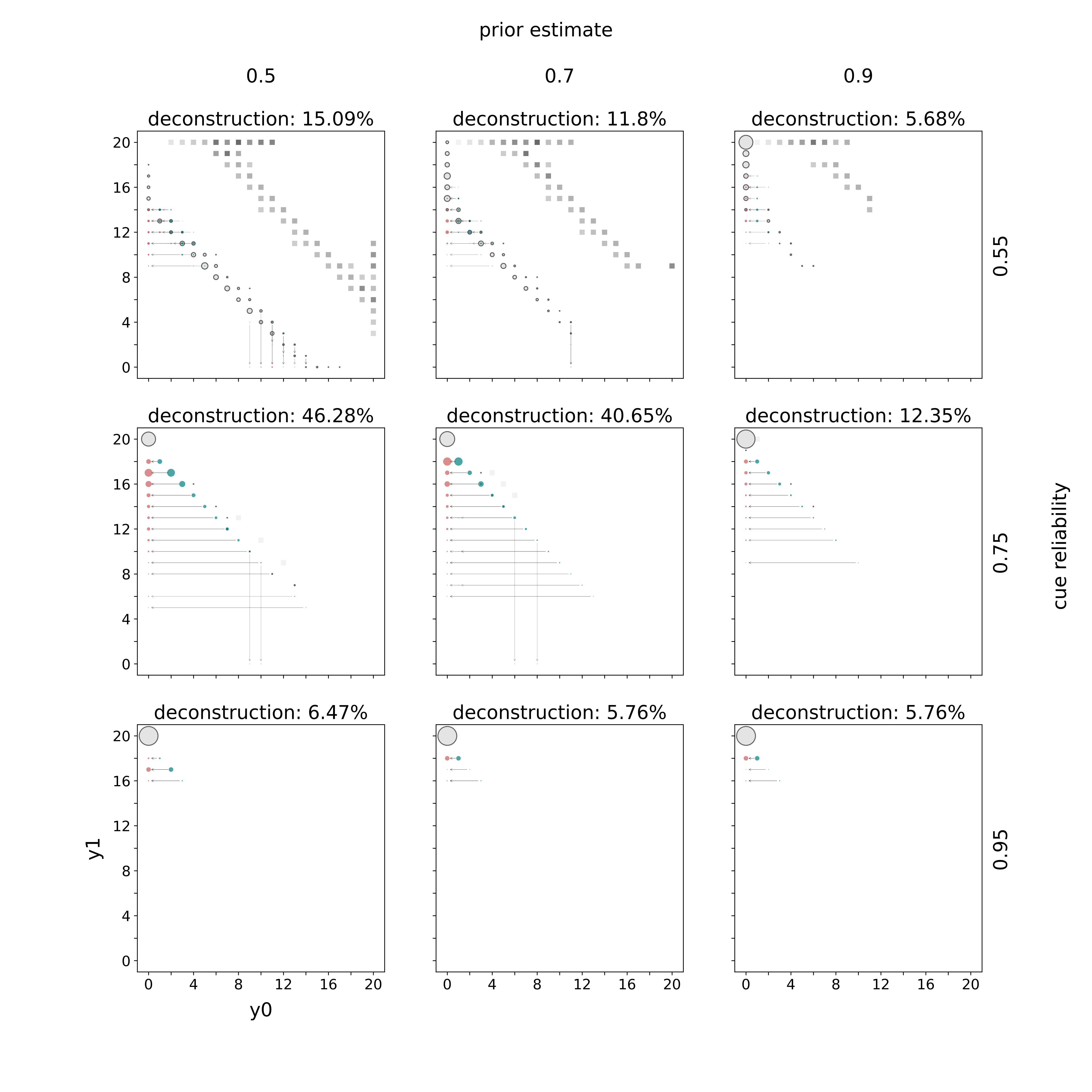

*Figure S2.34:* Distributions of mature phenotypes. Distributions of mature phenotypes are shown for a model with complete deconstruction and diminishing rewards and linear penalties. Rows indicate the prior estimate of being in $E_{1}$ and columns indicate the cue reliability. The populations of mature phenotypes have been simulated in $E_{1}$. The title of each panel indicates the percentage of mature phenotypes that have deconstructed at some point during ontogeny. Within each panel the horizontal axis indicates the number of specializations towards $E_{0}$ and the vertical axis towards $E_{1}$. The lower triangle indicates how much mature phenotypes have constructed (teal circles) and what their phenotype looked like after deconstruction (red circles). Grey arrows connect phenotypes before (teal) and after (red) deconstruction. Grey circles with a black outline belong to mature phenotypes that never deconstructed. The area of a circle is proportional to the number mature organisms with this phenotype. The upper triangle indicates waiting. For each mature phenotype (after deconstruction) below the diagonal the corresponding square above the diagonal highlights the amount of waiting. The color intensity is proportional to the amount of waiting. Black squares indicate phenotypes that waited all of ontogeny (i.e. 20 time periods) and white squares phenotypes that never waited.

*Diminishing rewards and increasing penalties*

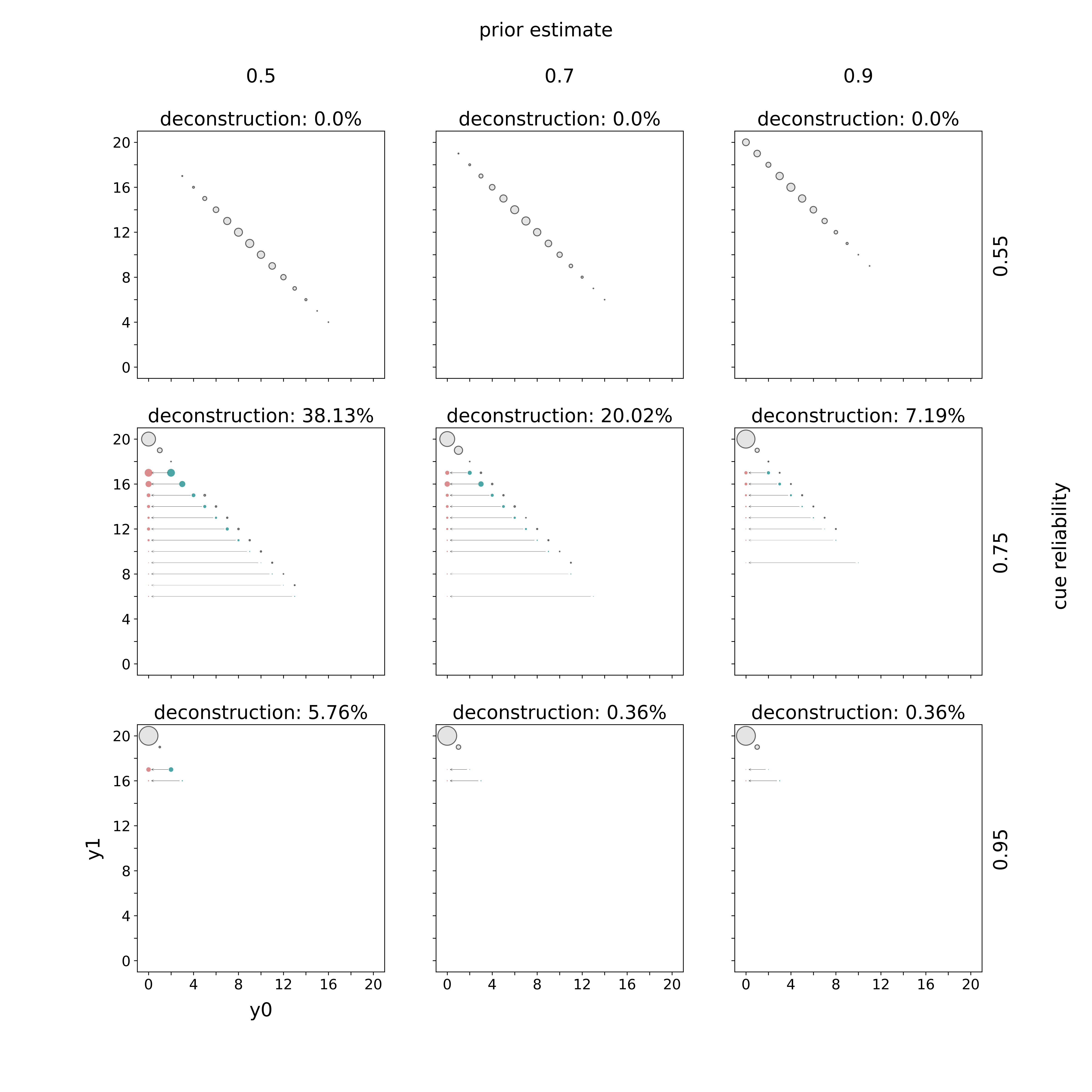

*Figure S2.35:* Distributions of mature phenotypes. Distributions of mature phenotypes are shown for a model with complete deconstruction and diminishing rewards and increasing penalties. Rows indicate the prior estimate of being in $E_{1}$ and columns indicate the cue reliability. The populations of mature phenotypes have been simulated in $E_{1}$. The title of each panel indicates the percentage of mature phenotypes that have deconstructed at some point during ontogeny. Within each panel the horizontal axis indicates the number of specializations towards $E_{0}$ and the vertical axis towards $E_{1}$. The lower triangle indicates how much mature phenotypes have constructed (teal circles) and what their phenotype looked like after deconstruction (red circles). Grey arrows connect phenotypes before (teal) and after (red) deconstruction. Grey circles with a black outline belong to mature phenotypes that never deconstructed. The area of a circle is proportional to the number mature organisms with this phenotype. The upper triangle indicates waiting. For each mature phenotype (after deconstruction) below the diagonal the corresponding square above the diagonal highlights the amount of waiting. The color intensity is proportional to the amount of waiting. Black squares indicate phenotypes that waited all of ontogeny (i.e. 20 time periods) and white squares phenotypes that never waited.

*Diminishing rewards and diminishing penalties*

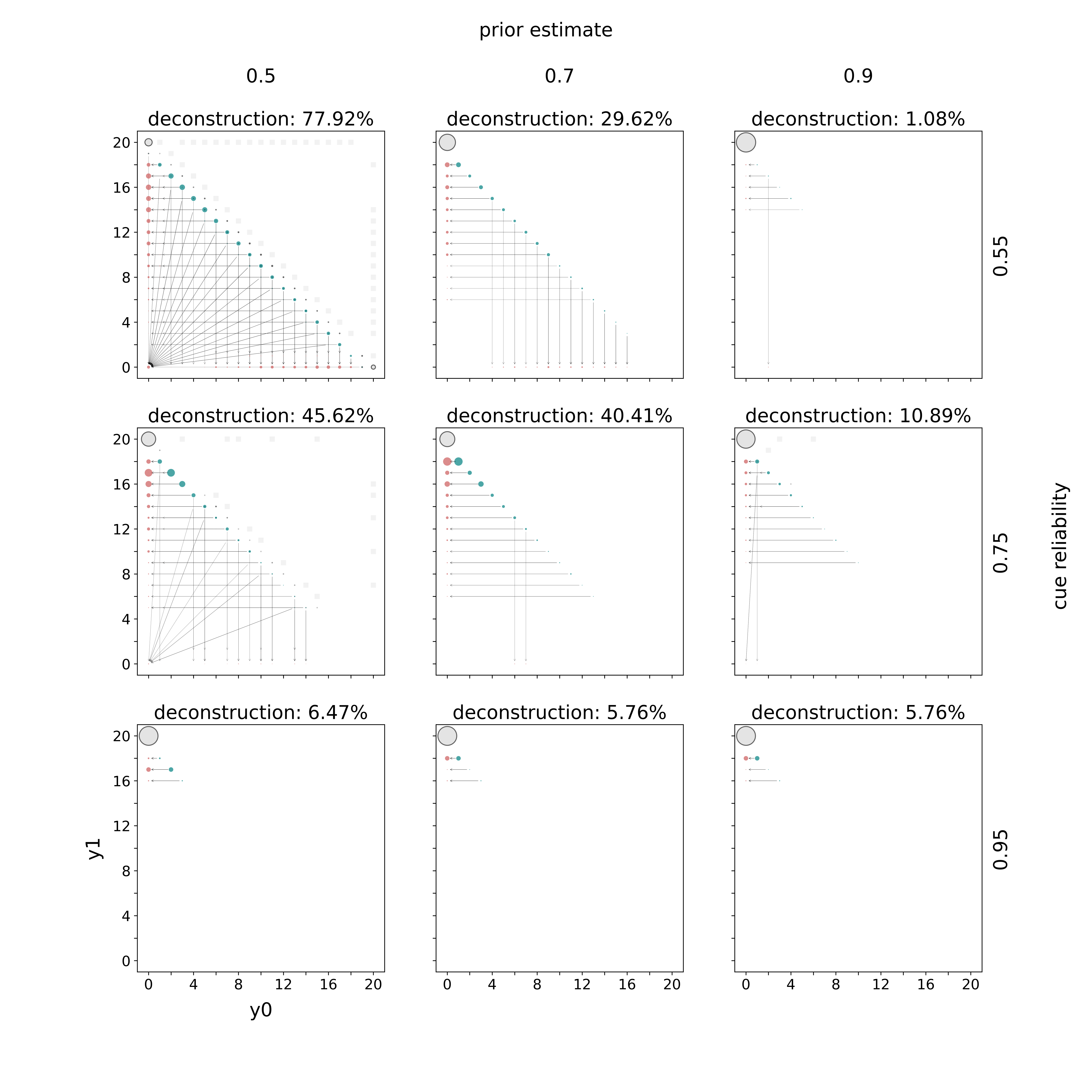

*Figure S2.36:* Distributions of mature phenotypes. Distributions of mature phenotypes are shown for a model with complete deconstruction and diminishing rewards and diminishing penalties. Rows indicate the prior estimate of being in $E_{1}$ and columns indicate the cue reliability. The populations of mature phenotypes have been simulated in $E_{1}$. The title of each panel indicates the percentage of mature phenotypes that have deconstructed at some point during ontogeny. Within each panel the horizontal axis indicates the number of specializations towards $E_{0}$ and the vertical axis towards $E_{1}$. The lower triangle indicates how much mature phenotypes have constructed (teal circles) and what their phenotype looked like after deconstruction (red circles). Grey arrows connect phenotypes before (teal) and after (red) deconstruction. Grey circles with a black outline belong to mature phenotypes that never deconstructed. The area of a circle is proportional to the number mature organisms with this phenotype. The upper triangle indicates waiting. For each mature phenotype (after deconstruction) below the diagonal the corresponding square above the diagonal highlights the amount of waiting. The color intensity is proportional to the amount of waiting. Black squares indicate phenotypes that waited all of ontogeny (i.e. 20 time periods) and white squares phenotypes that never waited.

**Fitness differences (incremental and complete deconstruction combined)**

*Linear rewards and linear penalties*

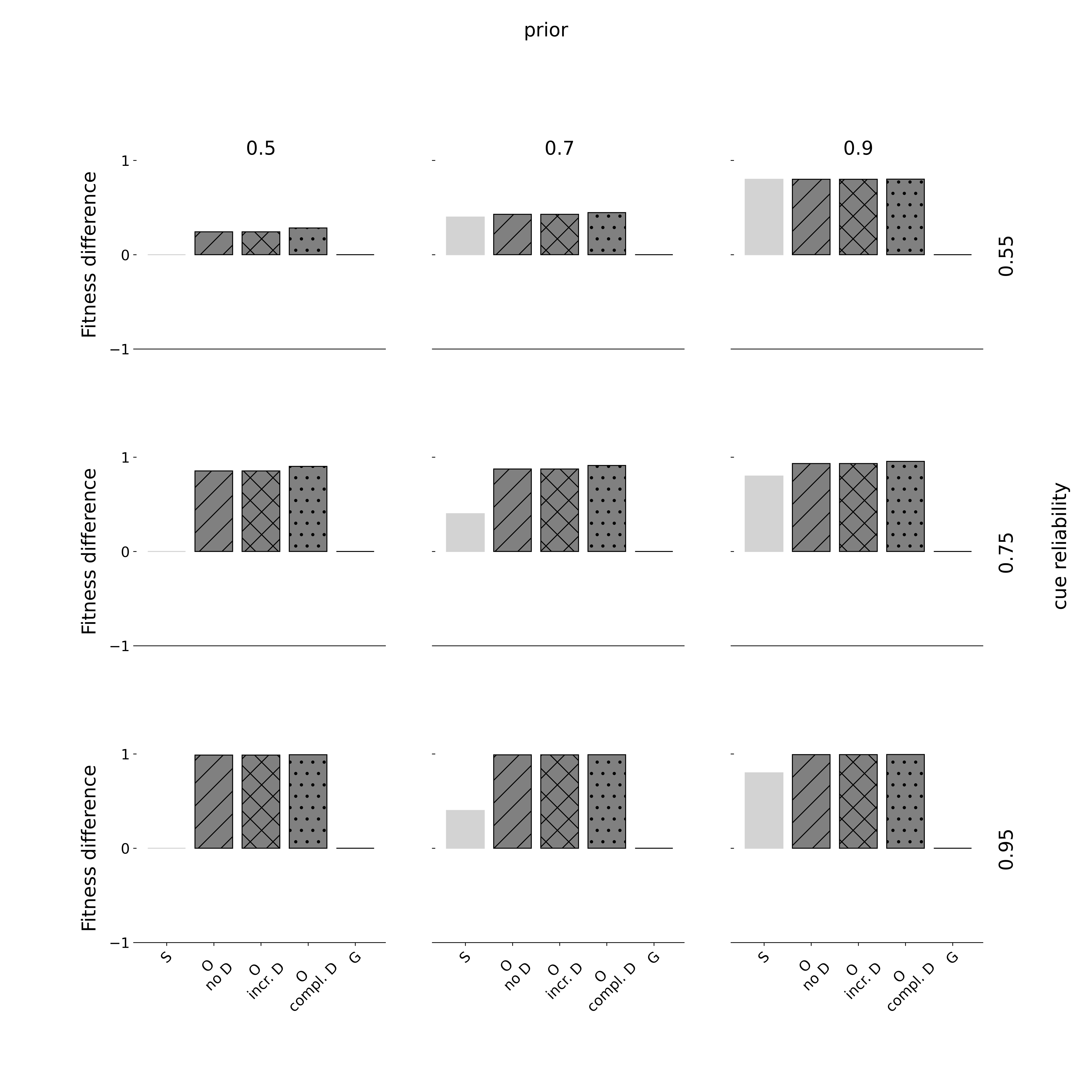

*Figure S2.37:* Fitness differences. Fitness differences are shown for linear rewards and linear penalties. Rows indicate the prior estimate of being in $E_{1}$ and columns indicate the cue reliability. Within each panel, the horizontal axis denotes the type of strategy where ‘S’ corresponds to a pure specialist strategy, ‘O’ to an optimal policy, and ‘G’ to a pure generalist strategy. The horizontal axis denotes fitness differences from baseline (corresponding to 0), normalized to range between -1 and 1. We show fitness of three different optimal policies: without deconstruction (‘no D’), with incremental deconstruction (‘incr. D’), and complete deconstruction (‘compl. D’). Specialists always fully specialize according to the prior distribution. When priors are uninformative (0.5), half the population fully specializes towards $P_{0}$ and the other one towards $P_{1}$. Generalists always specialize halfway towards either phenotypic target.

*Linear rewards and increasing penalties*

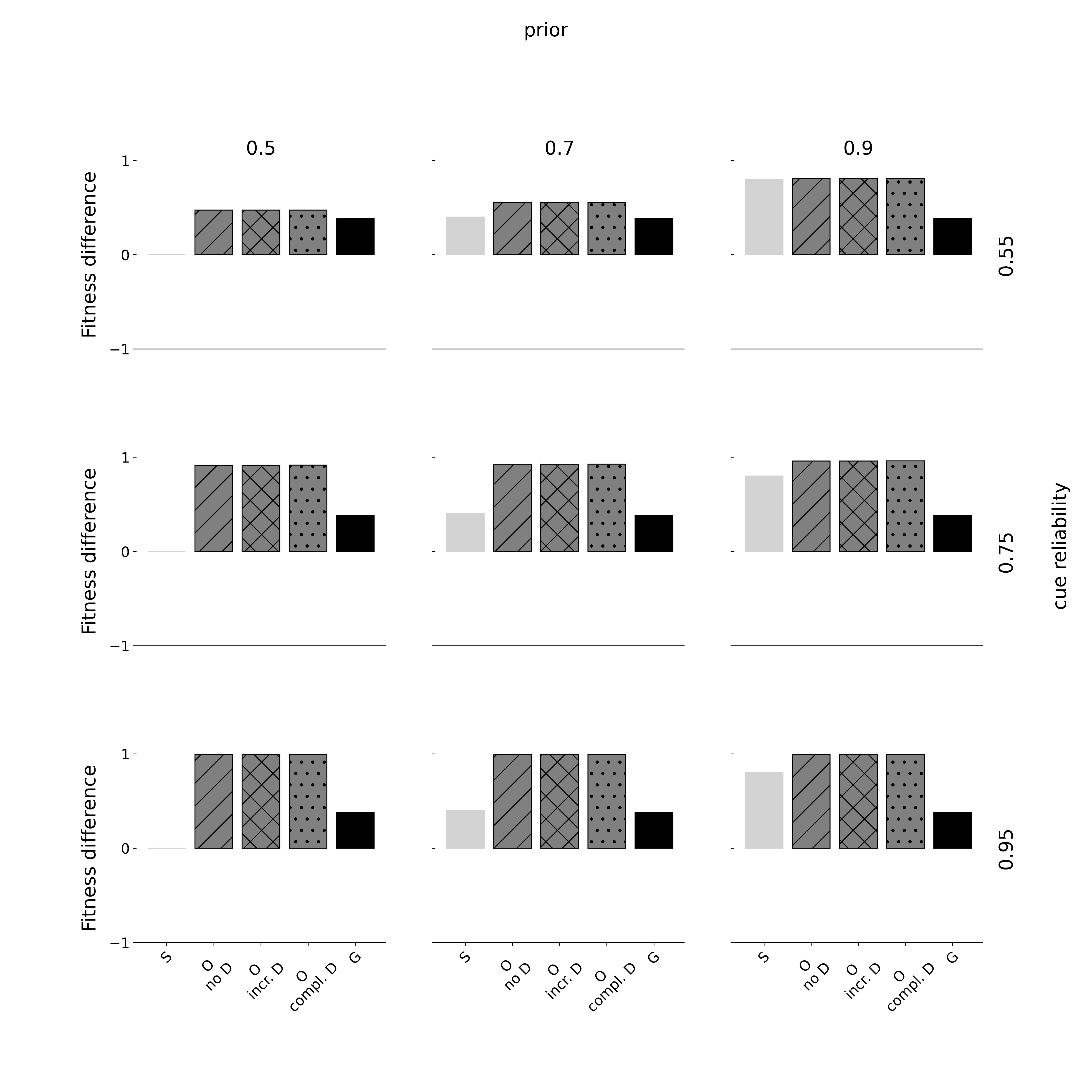

*Figure S2.38:* Fitness differences. Fitness differences are shown for linear rewards and increasing penalties. Rows indicate the prior estimate of being in $E_{1}$ and columns indicate the cue reliability. Within each panel, the horizontal axis denotes the type of strategy where ‘S’ corresponds to a pure specialist strategy, ‘O’ to an optimal policy, and ‘G’ to a pure generalist strategy. The horizontal axis denotes fitness differences from baseline (corresponding to 0), normalized to range between -1 and 1. We show fitness of three different optimal policies: without deconstruction (‘no D’), with incremental deconstruction (‘incr. D’), and complete deconstruction (‘compl. D’). Specialists always fully specialize according to the prior distribution. When priors are uninformative (0.5), half the population fully specializes towards $P_{0}$ and the other one towards $P_{1}$. Generalists always specialize halfway towards either phenotypic target.

*Linear rewards and diminishing penalties*

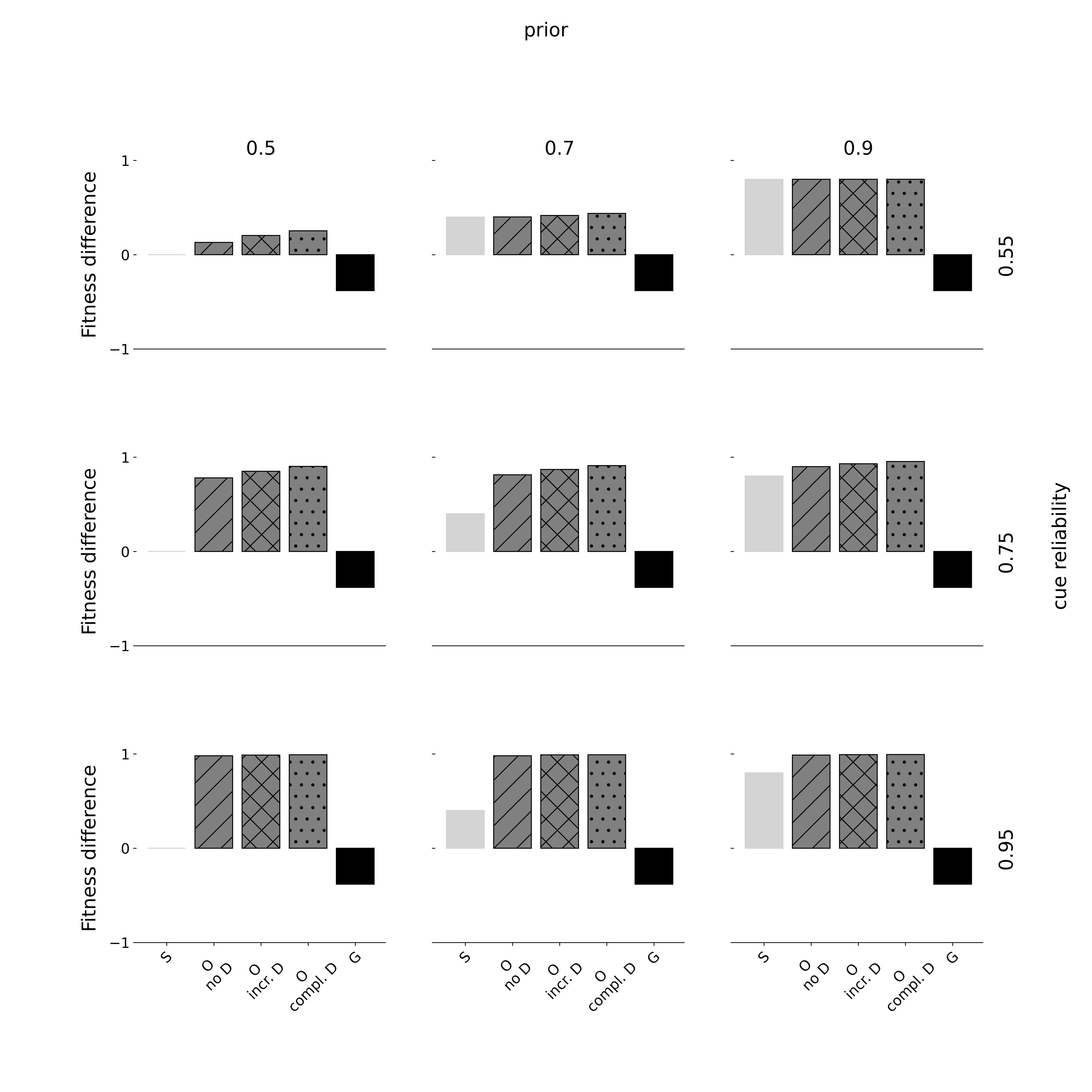

*Figure S2.39:* Fitness differences. Fitness differences are shown for linear rewards and diminishing penalties. Rows indicate the prior estimate of being in $E_{1}$ and columns indicate the cue reliability. Within each panel, the horizontal axis denotes the type of strategy where ‘S’ corresponds to a pure specialist strategy, ‘O’ to an optimal policy, and ‘G’ to a pure generalist strategy. The horizontal axis denotes fitness differences from baseline (corresponding to 0), normalized to range between -1 and 1. We show fitness of three different optimal policies: without deconstruction (‘no D’), with incremental deconstruction (‘incr. D’), and complete deconstruction (‘compl. D’). Specialists always fully specialize according to the prior distribution. When priors are uninformative (0.5), half the population fully specializes towards $P_{0}$ and the other one towards $P_{1}$. Generalists always specialize halfway towards either phenotypic target.

*Increasing rewards and linear penalties*

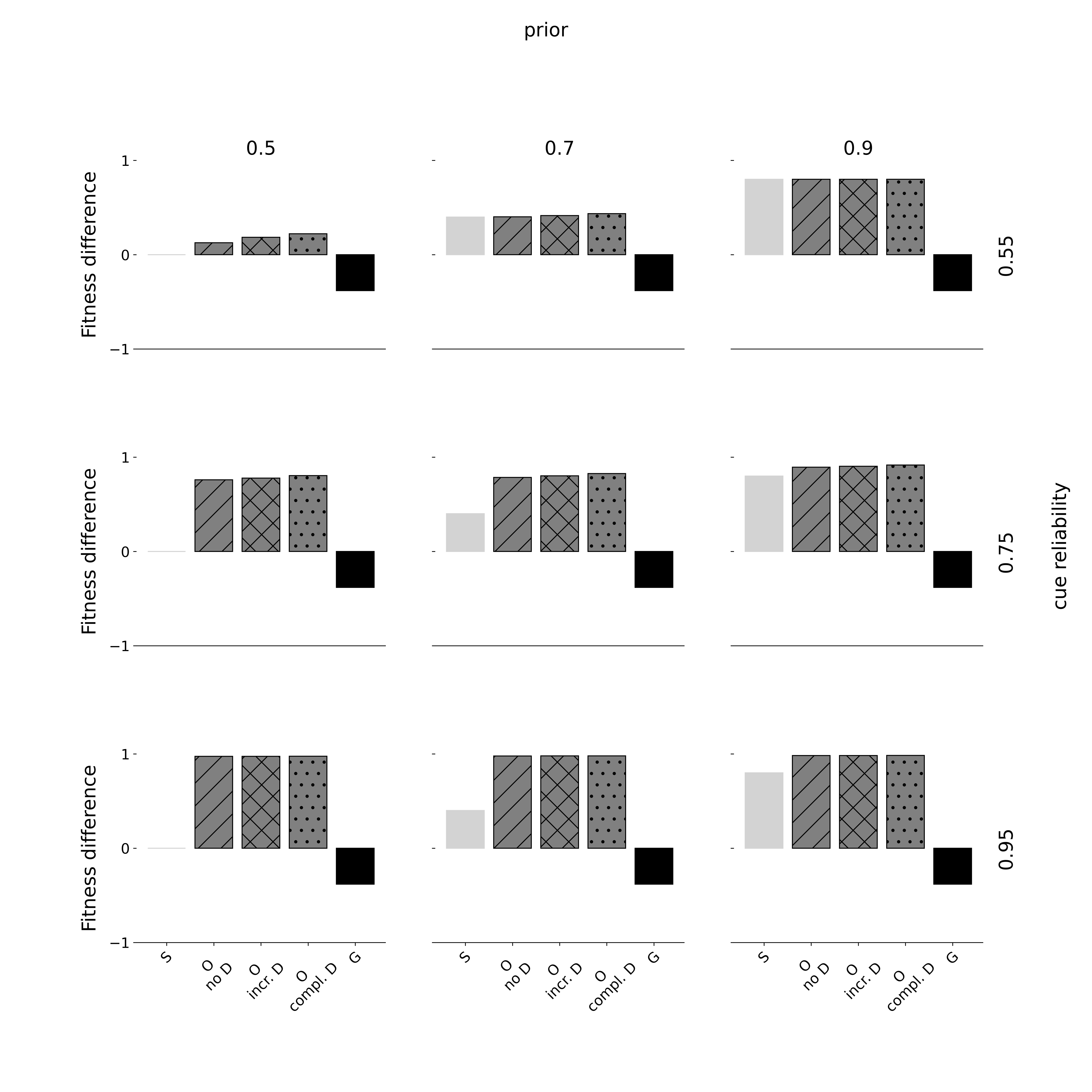

*Figure S2.40:* Fitness differences. Fitness differences are shown for increasing rewards and linear penalties. Rows indicate the prior estimate of being in $E_{1}$ and columns indicate the cue reliability. Within each panel, the horizontal axis denotes the type of strategy where ‘S’ corresponds to a pure specialist strategy, ‘O’ to an optimal policy, and ‘G’ to a pure generalist strategy. The horizontal axis denotes fitness differences from baseline (corresponding to 0), normalized to range between -1 and 1. We show fitness of three different optimal policies: without deconstruction (‘no D’), with incremental deconstruction (‘incr. D’), and complete deconstruction (‘compl. D’). Specialists always fully specialize according to the prior distribution. When priors are uninformative (0.5), half the population fully specializes towards $P_{0}$ and the other one towards $P_{1}$. Generalists always specialize halfway towards either phenotypic target.

*Increasing rewards and increasing penalties*

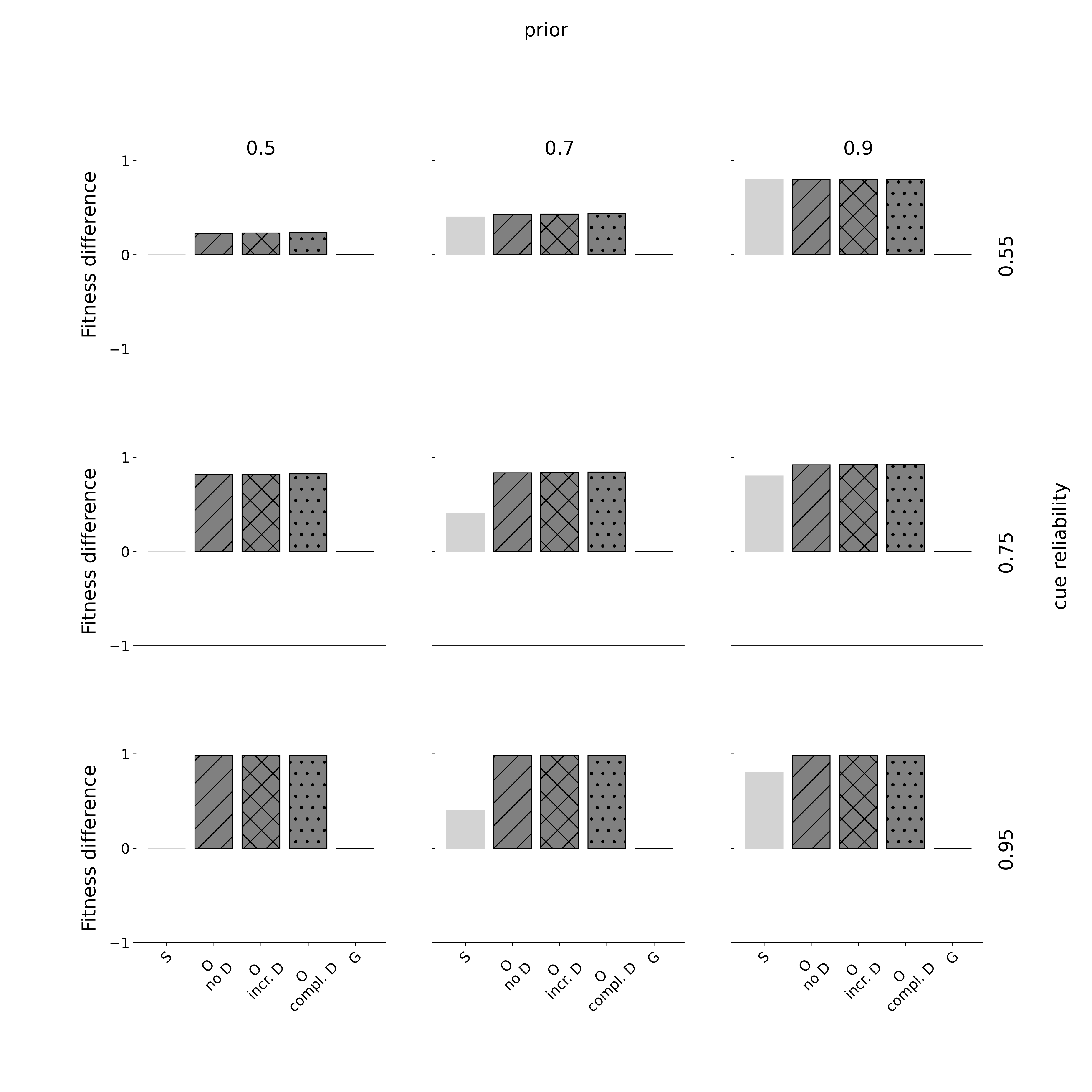

*Figure S2.41:* Fitness differences. Fitness differences are shown for increasing rewards and increasing penalties. Rows indicate the prior estimate of being in $E_{1}$ and columns indicate the cue reliability. Within each panel, the horizontal axis denotes the type of strategy where ‘S’ corresponds to a pure specialist strategy, ‘O’ to an optimal policy, and ‘G’ to a pure generalist strategy. The horizontal axis denotes fitness differences from baseline (corresponding to 0), normalized to range between -1 and 1. We show fitness of three different optimal policies: without deconstruction (‘no D’), with incremental deconstruction (‘incr. D’), and complete deconstruction (‘compl. D’). Specialists always fully specialize according to the prior distribution. When priors are uninformative (0.5), half the population fully specializes towards $P_{0}$ and the other one towards $P_{1}$. Generalists always specialize halfway towards either phenotypic target.

*Increasing rewards and diminishing penalties*

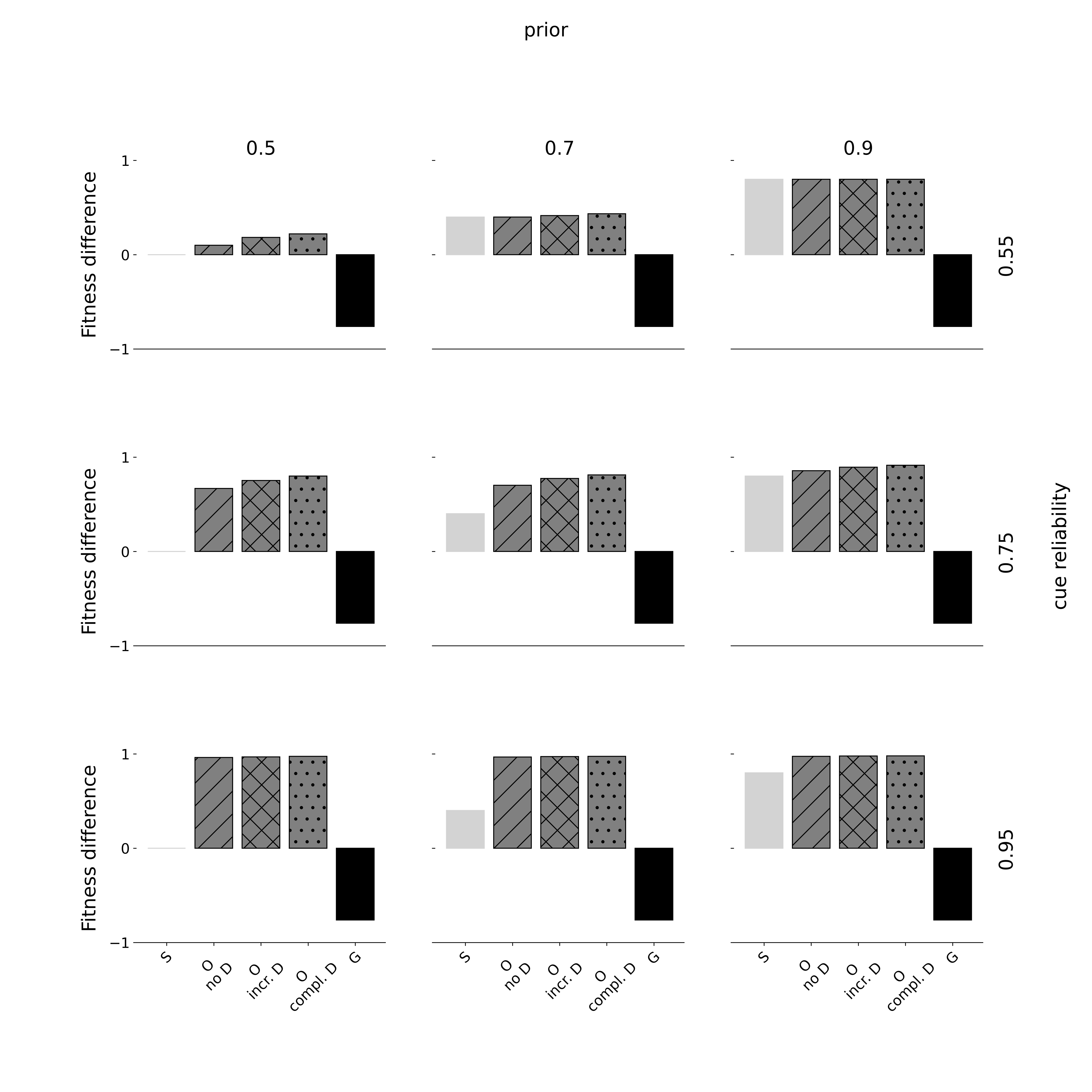

*Figure S2.42:* Fitness differences. Fitness differences are shown for increasing rewards and diminishing penalties. Rows indicate the prior estimate of being in $E_{1}$ and columns indicate the cue reliability. Within each panel, the horizontal axis denotes the type of strategy where ‘S’ corresponds to a pure specialist strategy, ‘O’ to an optimal policy, and ‘G’ to a pure generalist strategy. The horizontal axis denotes fitness differences from baseline (corresponding to 0), normalized to range between -1 and 1. We show fitness of three different optimal policies: without deconstruction (‘no D’), with incremental deconstruction (‘incr. D’), and complete deconstruction (‘compl. D’). Specialists always fully specialize according to the prior distribution. When priors are uninformative (0.5), half the population fully specializes towards $P_{0}$ and the other one towards $P_{1}$. Generalists always specialize halfway towards either phenotypic target.

*Diminishing rewards and linear penalties*

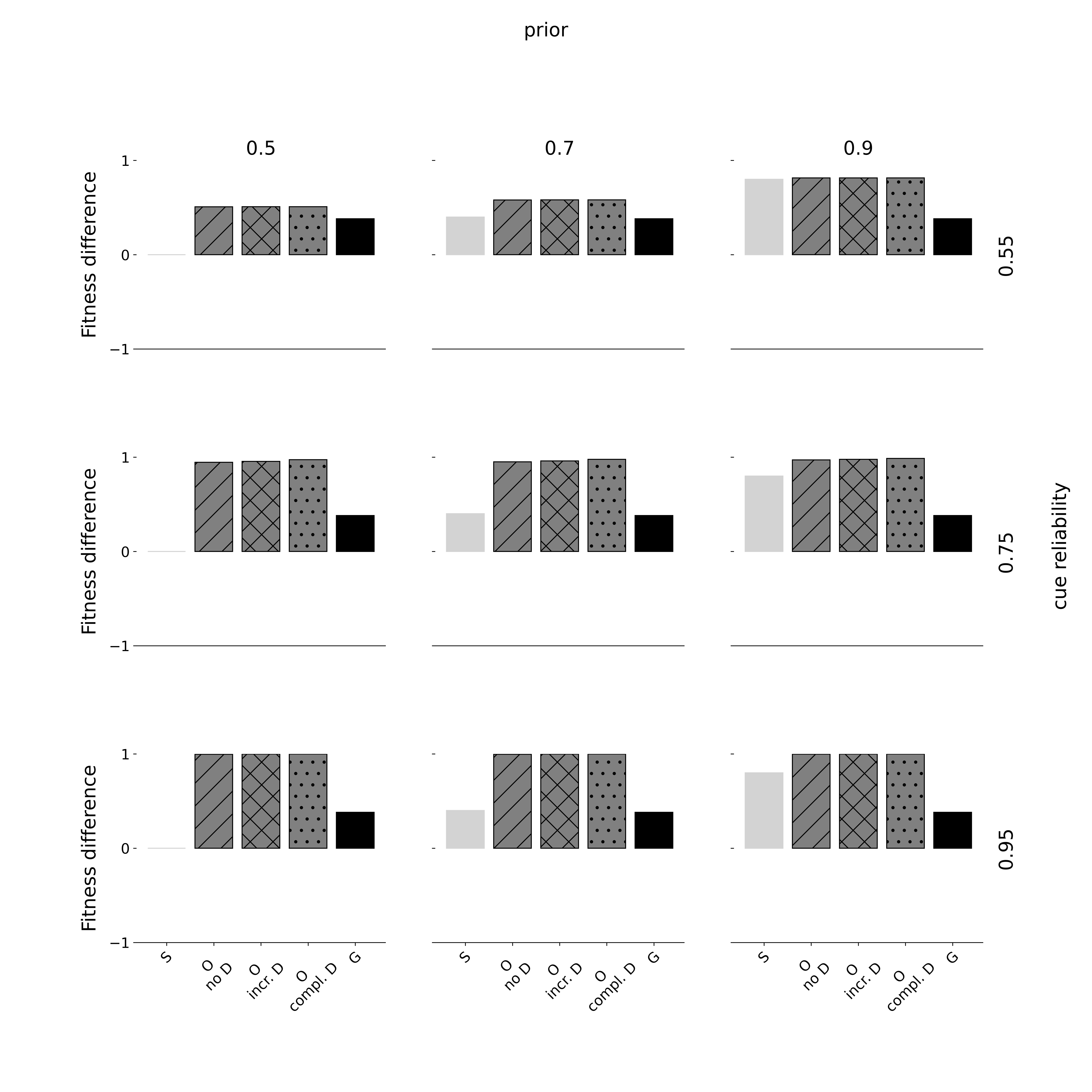

*Figure S2.43:* Fitness differences. Fitness differences are shown for diminishing rewards and linear penalties. Rows indicate the prior estimate of being in $E_{1}$ and columns indicate the cue reliability. Within each panel, the horizontal axis denotes the type of strategy where ‘S’ corresponds to a pure specialist strategy, ‘O’ to an optimal policy, and ‘G’ to a pure generalist strategy. The horizontal axis denotes fitness differences from baseline (corresponding to 0), normalized to range between -1 and 1. We show fitness of three different optimal policies: without deconstruction (‘no D’), with incremental deconstruction (‘incr. D’), and complete deconstruction (‘compl. D’). Specialists always fully specialize according to the prior distribution. When priors are uninformative (0.5), half the population fully specializes towards $P_{0}$ and the other one towards $P_{1}$. Generalists always specialize halfway towards either phenotypic target.

*Diminishing rewards and increasing penalties*

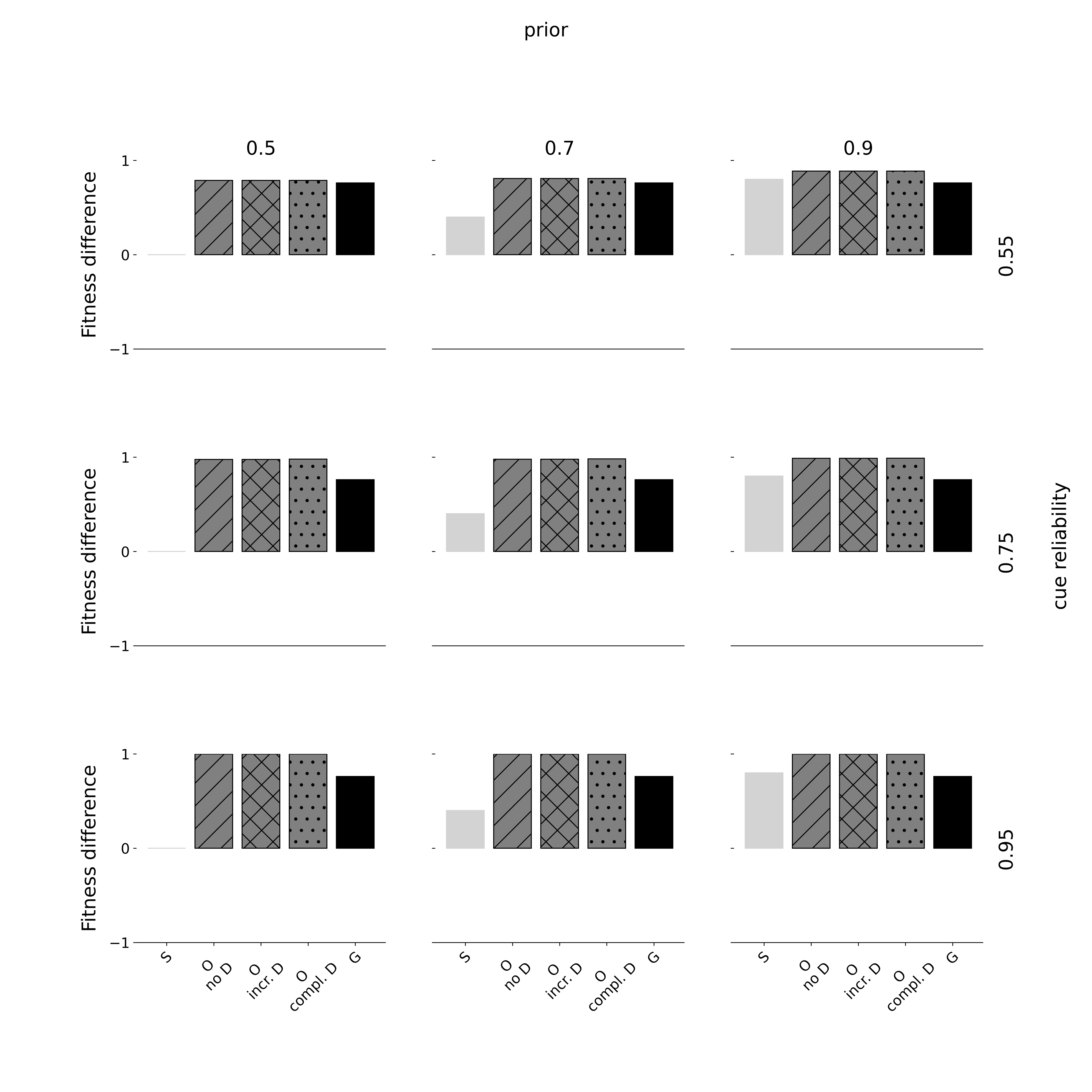

*Figure S2.44:* Fitness differences. Fitness differences are shown for diminishing rewards and increasing penalties. Rows indicate the prior estimate of being in $E_{1}$ and columns indicate the cue reliability. Within each panel, the horizontal axis denotes the type of strategy where ‘S’ corresponds to a pure specialist strategy, ‘O’ to an optimal policy, and ‘G’ to a pure generalist strategy. The horizontal axis denotes fitness differences from baseline (corresponding to 0), normalized to range between -1 and 1. We show fitness of three different optimal policies: without deconstruction (‘no D’), with incremental deconstruction (‘incr. D’), and complete deconstruction (‘compl. D’). Specialists always fully specialize according to the prior distribution. When priors are uninformative (0.5), half the population fully specializes towards $P_{0}$ and the other one towards $P_{1}$. Generalists always specialize halfway towards either phenotypic target.

*Diminishing rewards and diminishing penalties*

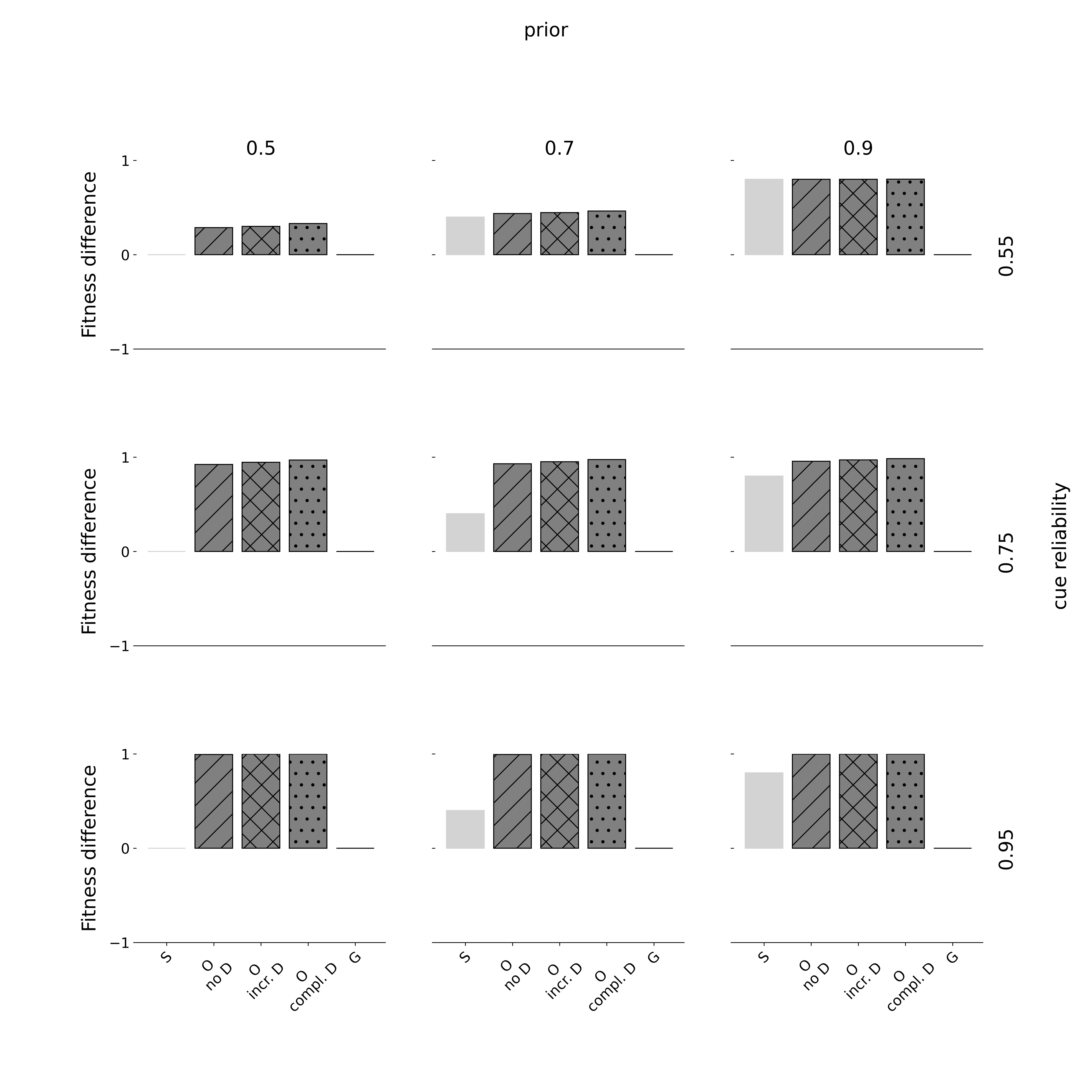

*Figure S2.45:* Fitness differences. Fitness differences are shown for diminishing rewards and diminishing penalties. Rows indicate the prior estimate of being in $E_{1}$ and columns indicate the cue reliability. Within each panel, the horizontal axis denotes the type of strategy where ‘S’ corresponds to a pure specialist strategy, ‘O’ to an optimal policy, and ‘G’ to a pure generalist strategy. The horizontal axis denotes fitness differences from baseline (corresponding to 0), normalized to range between -1 and 1. We show fitness of three different optimal policies: without deconstruction (‘no D’), with incremental deconstruction (‘incr. D’), and complete deconstruction (‘compl. D’). Specialists always fully specialize according to the prior distribution. When priors are uninformative (0.5), half the population fully specializes towards $P_{0}$ and the other one towards $P_{1}$. Generalists always specialize halfway towards either phenotypic target.

**Plasticity across ontogeny (incremental and complete deconstruction combined)**

*Linear rewards and linear penalties*

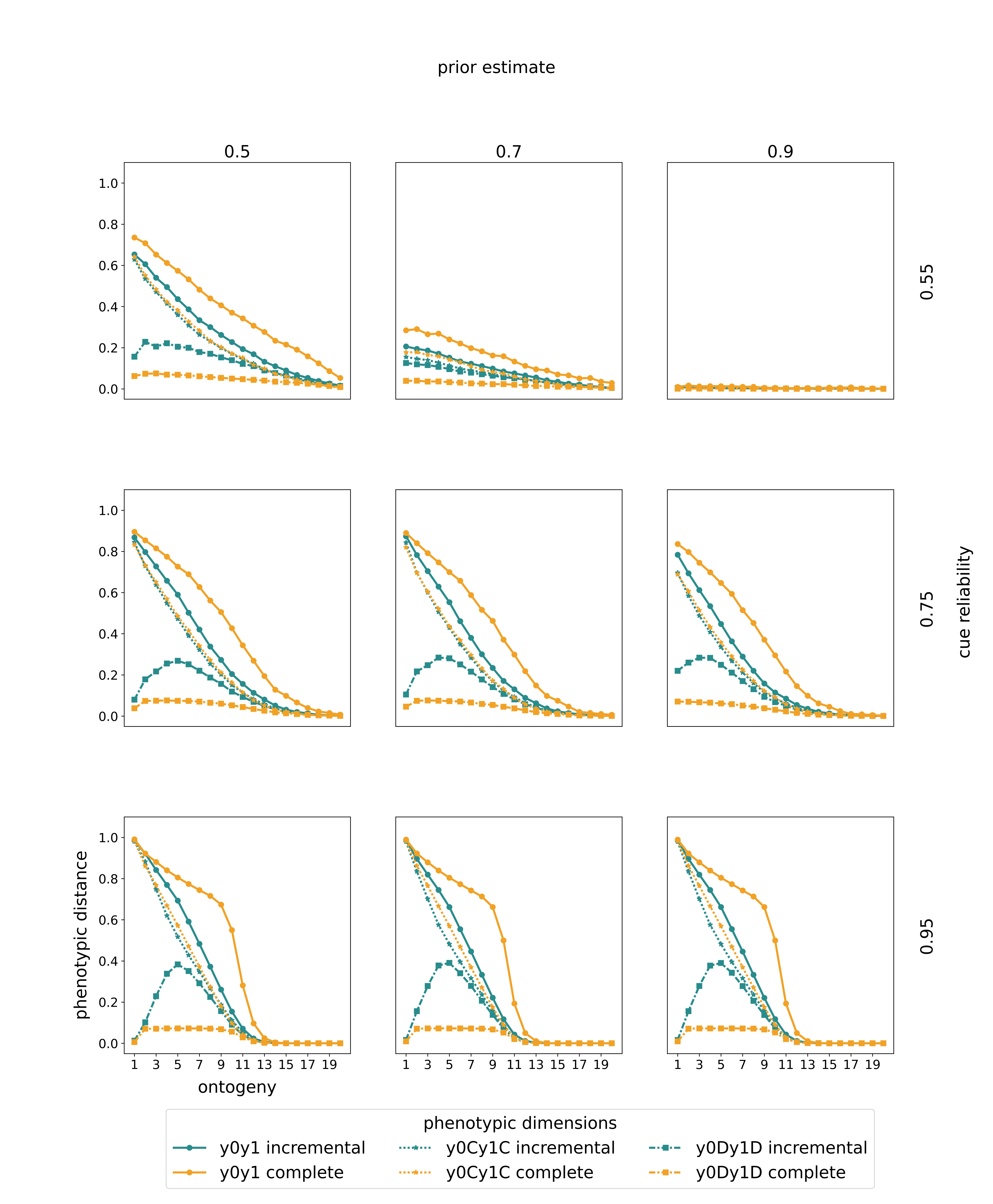

*Figure S2.46:* Changes in plasticity. Changes in plasticity are shown for linear rewards and linear penalties. Rows indicate the prior estimate of being in $E_{1}$ and columns indicate the cue reliability. Within each panel, the horizontal axis denotes ontogeny. The vertical axis denotes the normalized average phenotypic distance (our plasticity measure) across 10.000 pairs of simulated clones measured at the end of ontogeny. A specific point on any of the curves corresponds to a simulation experiment in which clones have been separated at the time point indicated by the horizontal axis. We show results from a model with incremental deconstruction in teal and with complete deconstruction in yellow. For each mode of deconstruction, we present three types of plasticity curves. First, we show plasticity in construction (dotted lines and stars) as the Euclidean distance between the number of time points spent constructing. Second, we show plasticity in deconstruction (dash-dotted lines and squares) as the Euclidean distance between the number of time points spent deconstruction. Third, we show total plasticity (solid lines and circles) as the Euclidean distance between the number of specialization steps towards either target (after accounting for deconstruction). We normalize phenotypic distance in construction and total phenotypes by dividing by the maximally possible Euclidean distance, corresponding to $2*\sqrt{T=20}$. The normalization constant for deconstruction is $2*\sqrt{T/2=10}$.

*Linear rewards and increasing penalties*

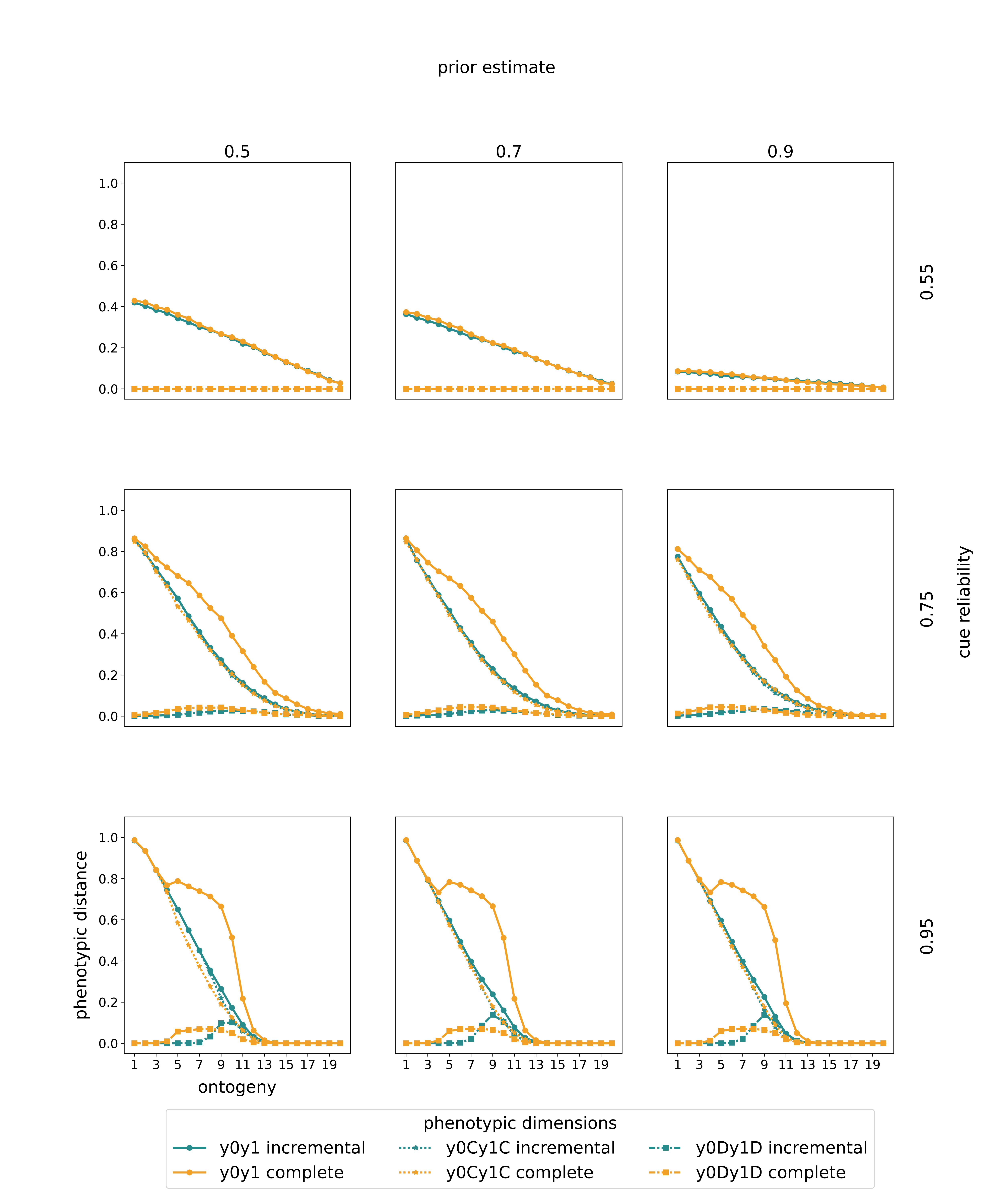

*Figure S2.47:* Changes in plasticity. Changes in plasticity are shown for linear rewards and increasing penalties. Rows indicate the prior estimate of being in $E_{1}$ and columns indicate the cue reliability. Within each panel, the horizontal axis denotes ontogeny. The vertical axis denotes the normalized average phenotypic distance (our plasticity measure) across 10.000 pairs of simulated clones measured at the end of ontogeny. A specific point on any of the curves corresponds to a simulation experiment in which clones have been separated at the time point indicated by the horizontal axis. We show results from a model with incremental deconstruction in teal and with complete deconstruction in yellow. For each mode of deconstruction, we present three types of plasticity curves. First, we show plasticity in construction (dotted lines and stars) as the Euclidean distance between the number of time points spent constructing. Second, we show plasticity in deconstruction (dash-dotted lines and squares) as the Euclidean distance between the number of time points spent deconstruction. Third, we show total plasticity (solid lines and circles) as the Euclidean distance between the number of specialization steps towards either target (after accounting for deconstruction). We normalize phenotypic distance in construction and total phenotypes by dividing by the maximally possible Euclidean distance, corresponding to $2*\sqrt{T=20}$. The normalization constant for deconstruction is $2*\sqrt{T/2=10}$.

*Linear rewards and diminishing penalties*

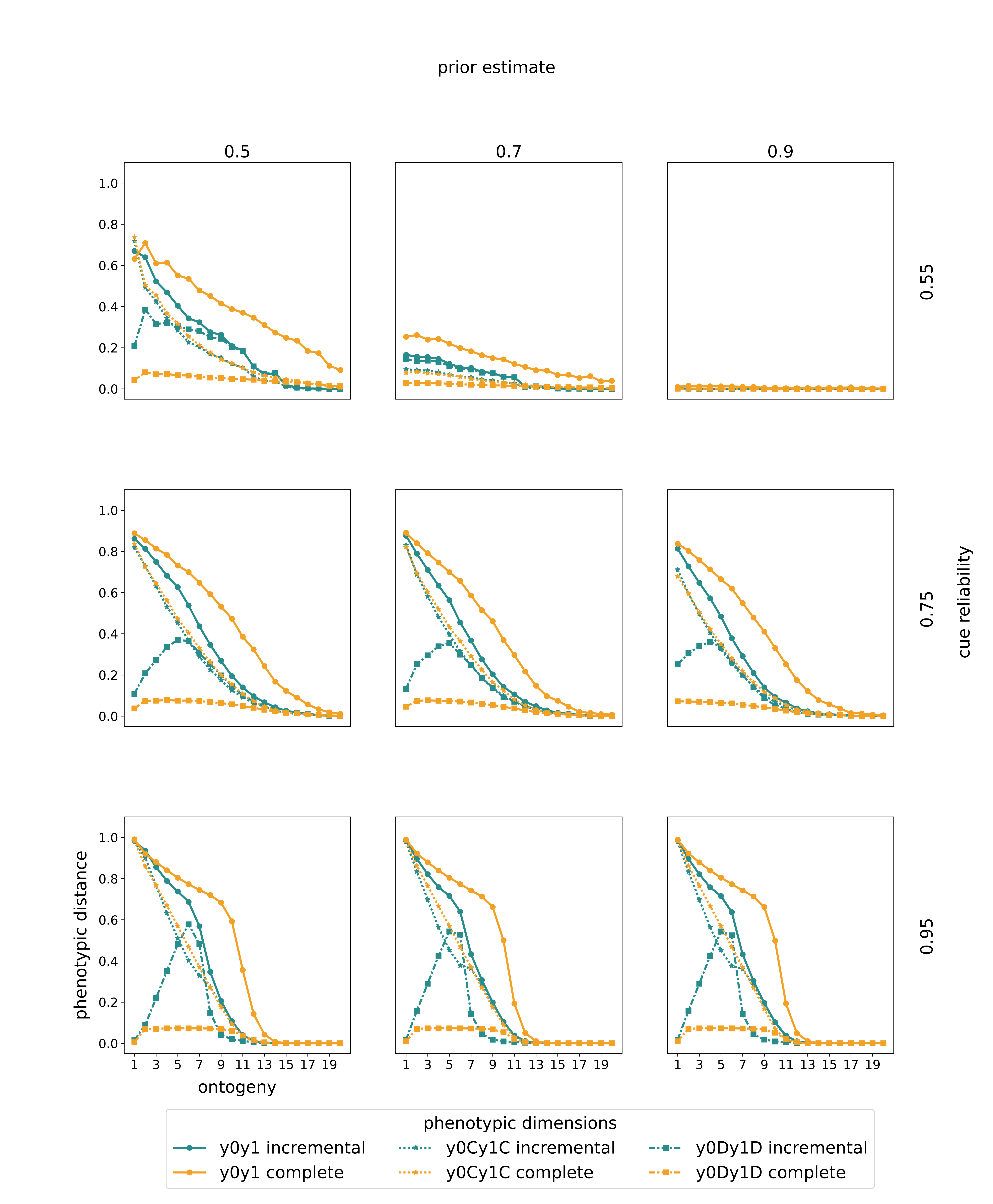

*Figure S2.48:* Changes in plasticity. Changes in plasticity are shown for linear rewards and diminishing penalties. Rows indicate the prior estimate of being in $E_{1}$ and columns indicate the cue reliability. Within each panel, the horizontal axis denotes ontogeny. The vertical axis denotes the normalized average phenotypic distance (our plasticity measure) across 10.000 pairs of simulated clones measured at the end of ontogeny. A specific point on any of the curves corresponds to a simulation experiment in which clones have been separated at the time point indicated by the horizontal axis. We show results from a model with incremental deconstruction in teal and with complete deconstruction in yellow. For each mode of deconstruction, we present three types of plasticity curves. First, we show plasticity in construction (dotted lines and stars) as the Euclidean distance between the number of time points spent constructing. Second, we show plasticity in deconstruction (dash-dotted lines and squares) as the Euclidean distance between the number of time points spent deconstruction. Third, we show total plasticity (solid lines and circles) as the Euclidean distance between the number of specialization steps towards either target (after accounting for deconstruction). We normalize phenotypic distance in construction and total phenotypes by dividing by the maximally possible Euclidean distance, corresponding to $2*\sqrt{T=20}$. The normalization constant for deconstruction is $2*\sqrt{T/2=10}$.

*Increasing rewards and linear penalties*

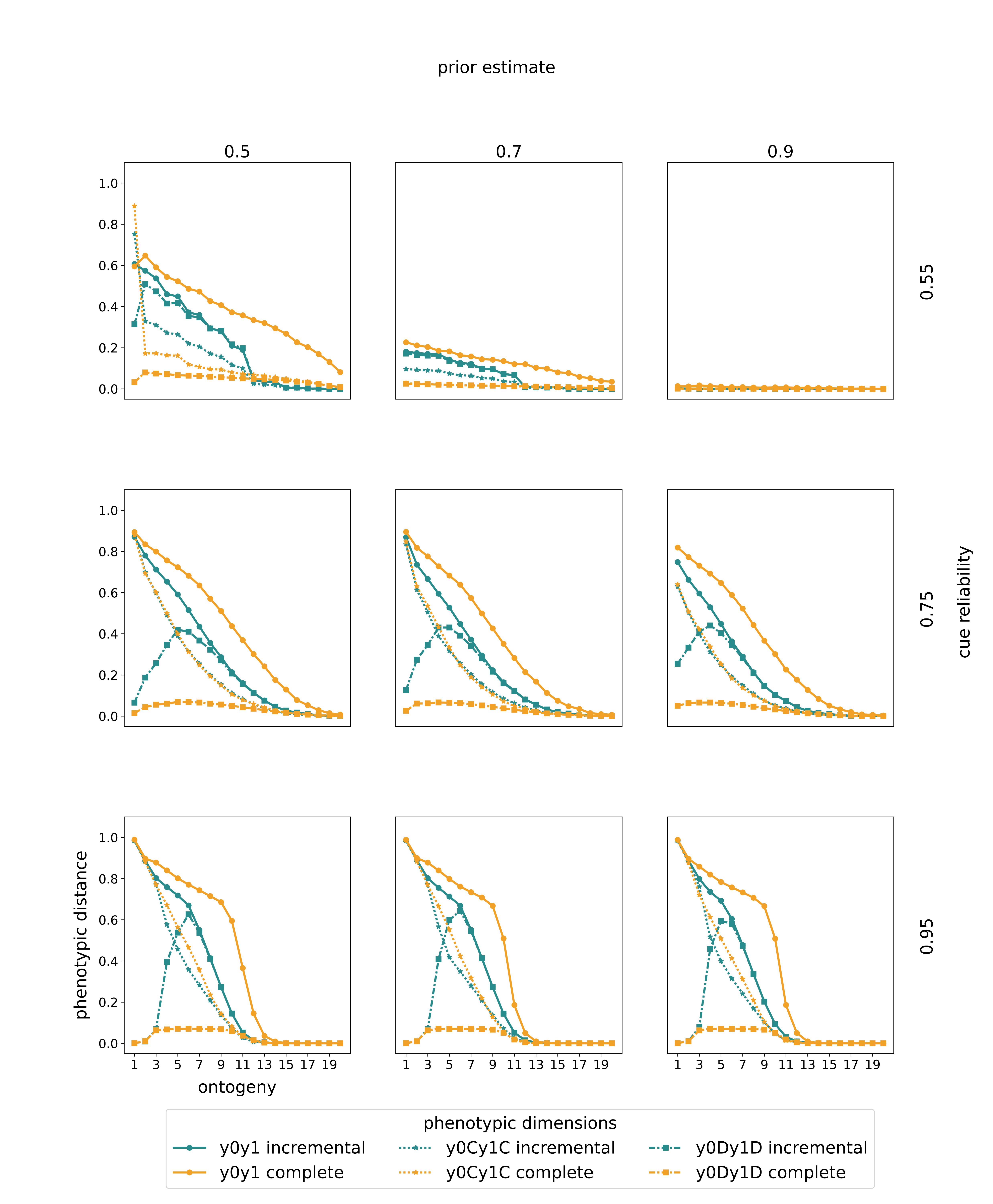

*Figure S2.49:* Changes in plasticity. Changes in plasticity are shown for increasing rewards and linear penalties. Rows indicate the prior estimate of being in $E_{1}$ and columns indicate the cue reliability. Within each panel, the horizontal axis denotes ontogeny. The vertical axis denotes the normalized average phenotypic distance (our plasticity measure) across 10.000 pairs of simulated clones measured at the end of ontogeny. A specific point on any of the curves corresponds to a simulation experiment in which clones have been separated at the time point indicated by the horizontal axis. We show results from a model with incremental deconstruction in teal and with complete deconstruction in yellow. For each mode of deconstruction, we present three types of plasticity curves. First, we show plasticity in construction (dotted lines and stars) as the Euclidean distance between the number of time points spent constructing. Second, we show plasticity in deconstruction (dash-dotted lines and squares) as the Euclidean distance between the number of time points spent deconstruction. Third, we show total plasticity (solid lines and circles) as the Euclidean distance between the number of specialization steps towards either target (after accounting for deconstruction). We normalize phenotypic distance in construction and total phenotypes by dividing by the maximally possible Euclidean distance, corresponding to $2*\sqrt{T=20}$. The normalization constant for deconstruction is $2*\sqrt{T/2=10}$.

*Increasing rewards and increasing penalties*

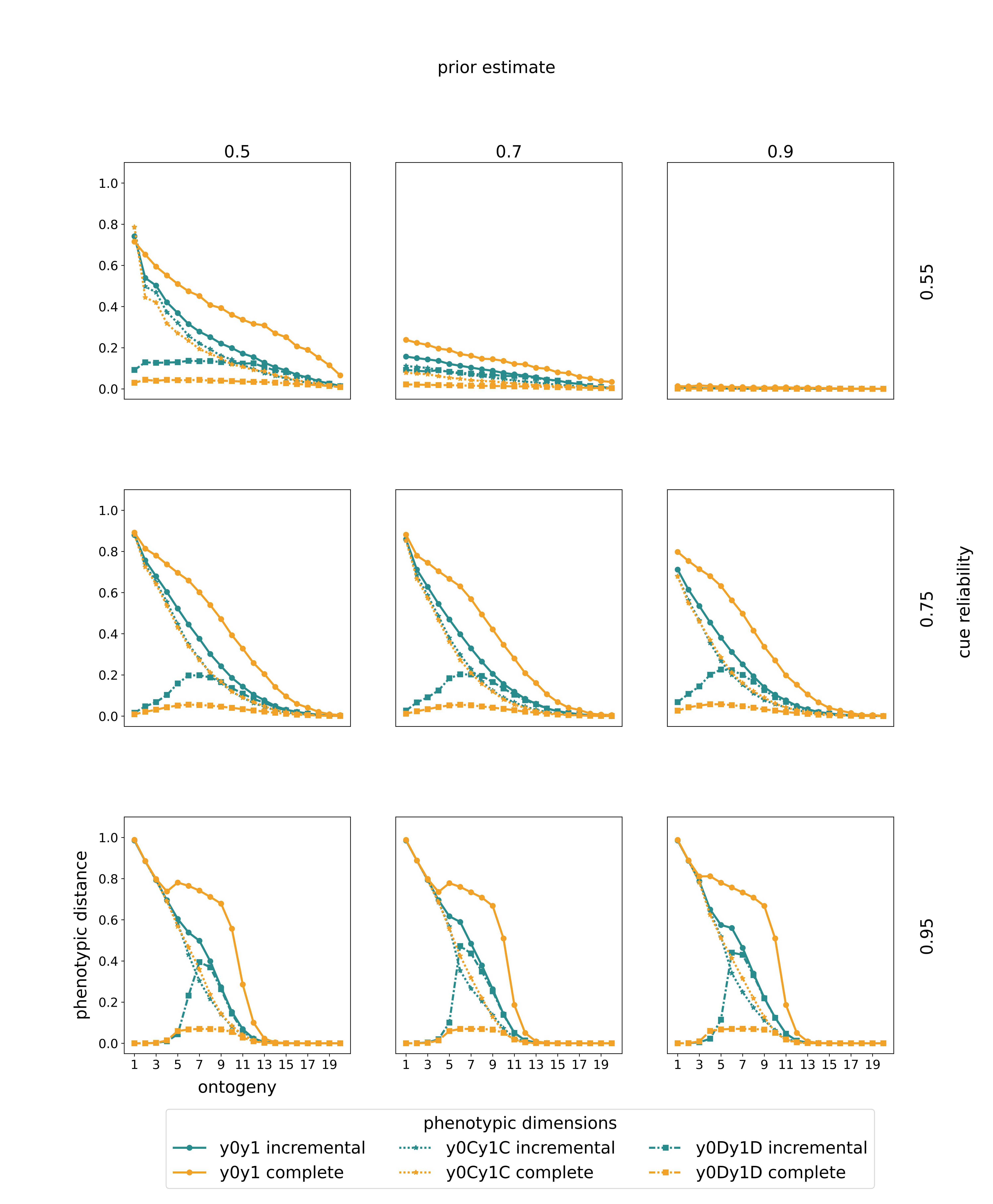

*Figure S2.50:* Changes in plasticity. Changes in plasticity are shown for increasing rewards and increasing penalties. Rows indicate the prior estimate of being in $E_{1}$ and columns indicate the cue reliability. Within each panel, the horizontal axis denotes ontogeny. The vertical axis denotes the normalized average phenotypic distance (our plasticity measure) across 10.000 pairs of simulated clones measured at the end of ontogeny. A specific point on any of the curves corresponds to a simulation experiment in which clones have been separated at the time point indicated by the horizontal axis. We show results from a model with incremental deconstruction in teal and with complete deconstruction in yellow. For each mode of deconstruction, we present three types of plasticity curves. First, we show plasticity in construction (dotted lines and stars) as the Euclidean distance between the number of time points spent constructing. Second, we show plasticity in deconstruction (dash-dotted lines and squares) as the Euclidean distance between the number of time points spent deconstruction. Third, we show total plasticity (solid lines and circles) as the Euclidean distance between the number of specialization steps towards either target (after accounting for deconstruction). We normalize phenotypic distance in construction and total phenotypes by dividing by the maximally possible Euclidean distance, corresponding to $2*\sqrt{T=20}$. The normalization constant for deconstruction is $2*\sqrt{T/2=10}$.

*Increasing rewards and diminishing penalties*

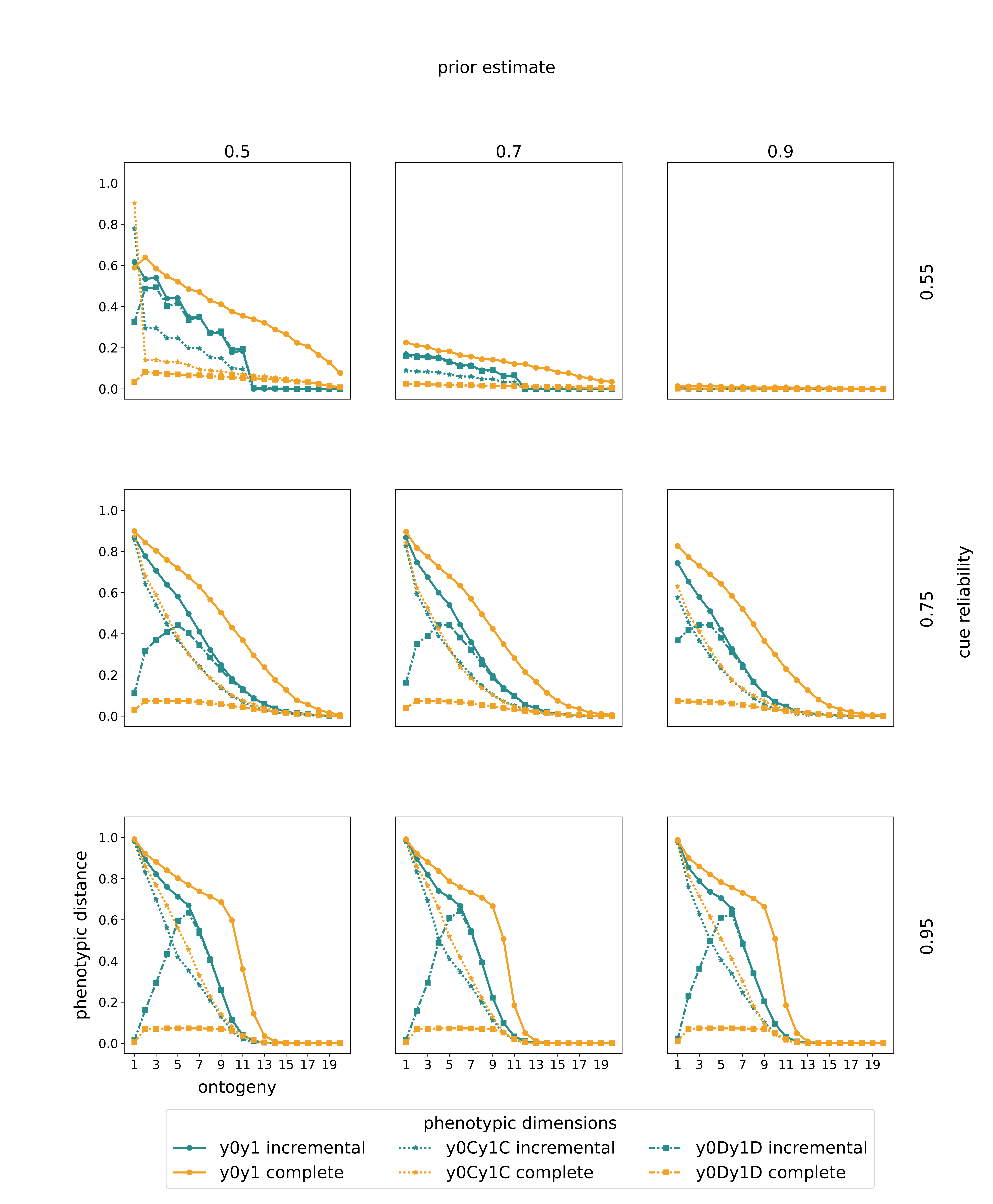

*Figure S2.51:* Changes in plasticity. Changes in plasticity are shown for increasing rewards and diminishing penalties. Rows indicate the prior estimate of being in $E_{1}$ and columns indicate the cue reliability. Within each panel, the horizontal axis denotes ontogeny. The vertical axis denotes the normalized average phenotypic distance (our plasticity measure) across 10.000 pairs of simulated clones measured at the end of ontogeny. A specific point on any of the curves corresponds to a simulation experiment in which clones have been separated at the time point indicated by the horizontal axis. We show results from a model with incremental deconstruction in teal and with complete deconstruction in yellow. For each mode of deconstruction, we present three types of plasticity curves. First, we show plasticity in construction (dotted lines and stars) as the Euclidean distance between the number of time points spent constructing. Second, we show plasticity in deconstruction (dash-dotted lines and squares) as the Euclidean distance between the number of time points spent deconstruction. Third, we show total plasticity (solid lines and circles) as the Euclidean distance between the number of specialization steps towards either target (after accounting for deconstruction). We normalize phenotypic distance in construction and total phenotypes by dividing by the maximally possible Euclidean distance, corresponding to $2*\sqrt{T=20}$. The normalization constant for deconstruction is $2*\sqrt{T/2=10}$.

*Diminishing rewards and linear penalties*

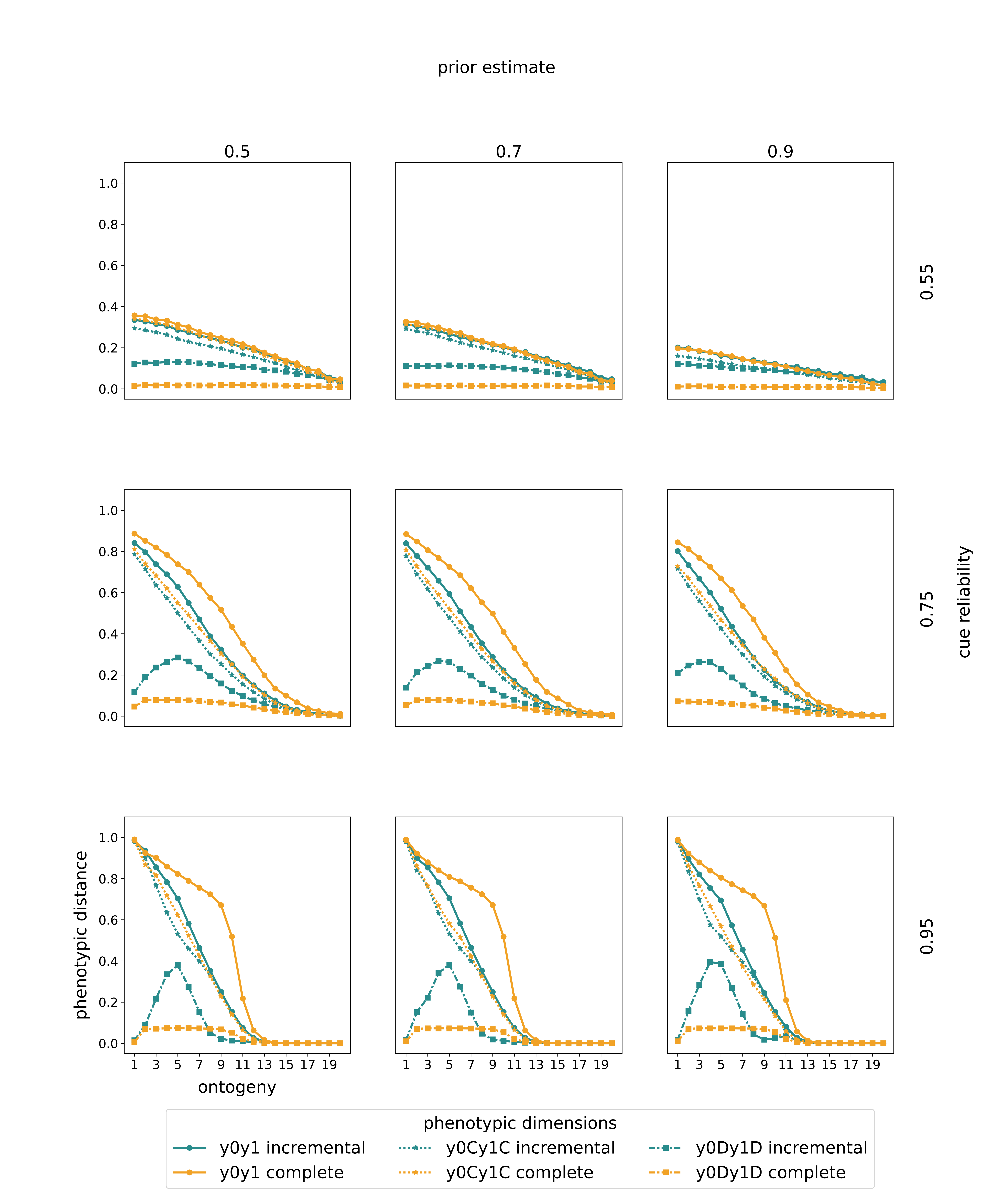

*Figure S2.52:* Changes in plasticity. Changes in plasticity are shown for diminishing rewards and linear penalties. Rows indicate the prior estimate of being in $E_{1}$ and columns indicate the cue reliability. Within each panel, the horizontal axis denotes ontogeny. The vertical axis denotes the normalized average phenotypic distance (our plasticity measure) across 10.000 pairs of simulated clones measured at the end of ontogeny. A specific point on any of the curves corresponds to a simulation experiment in which clones have been separated at the time point indicated by the horizontal axis. We show results from a model with incremental deconstruction in teal and with complete deconstruction in yellow. For each mode of deconstruction, we present three types of plasticity curves. First, we show plasticity in construction (dotted lines and stars) as the Euclidean distance between the number of time points spent constructing. Second, we show plasticity in deconstruction (dash-dotted lines and squares) as the Euclidean distance between the number of time points spent deconstruction. Third, we show total plasticity (solid lines and circles) as the Euclidean distance between the number of specialization steps towards either target (after accounting for deconstruction). We normalize phenotypic distance in construction and total phenotypes by dividing by the maximally possible Euclidean distance, corresponding to $2*\sqrt{T=20}$. The normalization constant for deconstruction is $2*\sqrt{T/2=10}$.

*Diminishing rewards and increasing penalties*

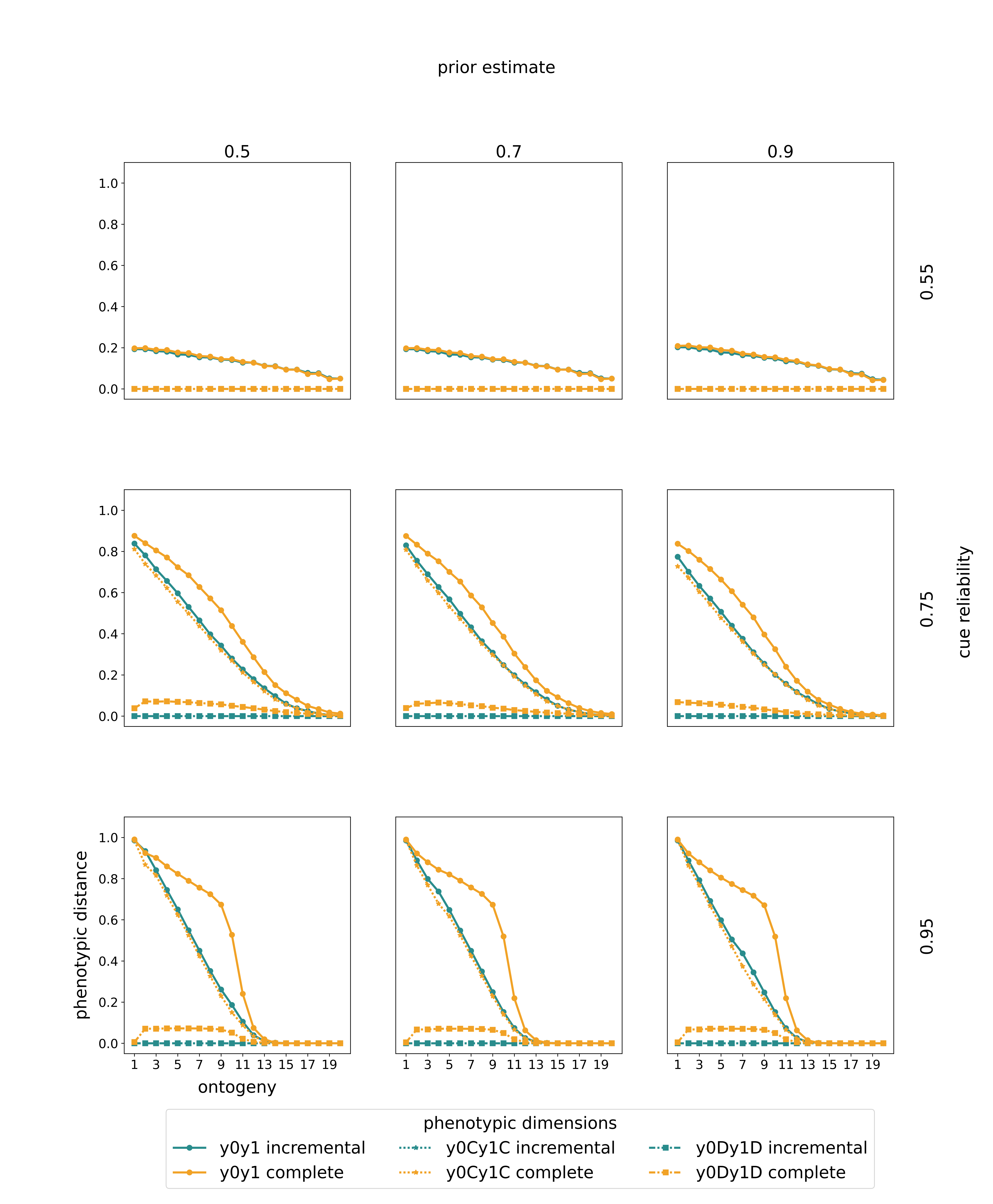

*Figure S2.53:* Changes in plasticity. Changes in plasticity are shown for diminishing rewards and increasing penalties. Rows indicate the prior estimate of being in $E_{1}$ and columns indicate the cue reliability. Within each panel, the horizontal axis denotes ontogeny. The vertical axis denotes the normalized average phenotypic distance (our plasticity measure) across 10.000 pairs of simulated clones measured at the end of ontogeny. A specific point on any of the curves corresponds to a simulation experiment in which clones have been separated at the time point indicated by the horizontal axis. We show results from a model with incremental deconstruction in teal and with complete deconstruction in yellow. For each mode of deconstruction, we present three types of plasticity curves. First, we show plasticity in construction (dotted lines and stars) as the Euclidean distance between the number of time points spent constructing. Second, we show plasticity in deconstruction (dash-dotted lines and squares) as the Euclidean distance between the number of time points spent deconstruction. Third, we show total plasticity (solid lines and circles) as the Euclidean distance between the number of specialization steps towards either target (after accounting for deconstruction). We normalize phenotypic distance in construction and total phenotypes by dividing by the maximally possible Euclidean distance, corresponding to $2*\sqrt{T=20}$. The normalization constant for deconstruction is $2*\sqrt{T/2=10}$.

*Diminishing rewards and diminishing penalties*

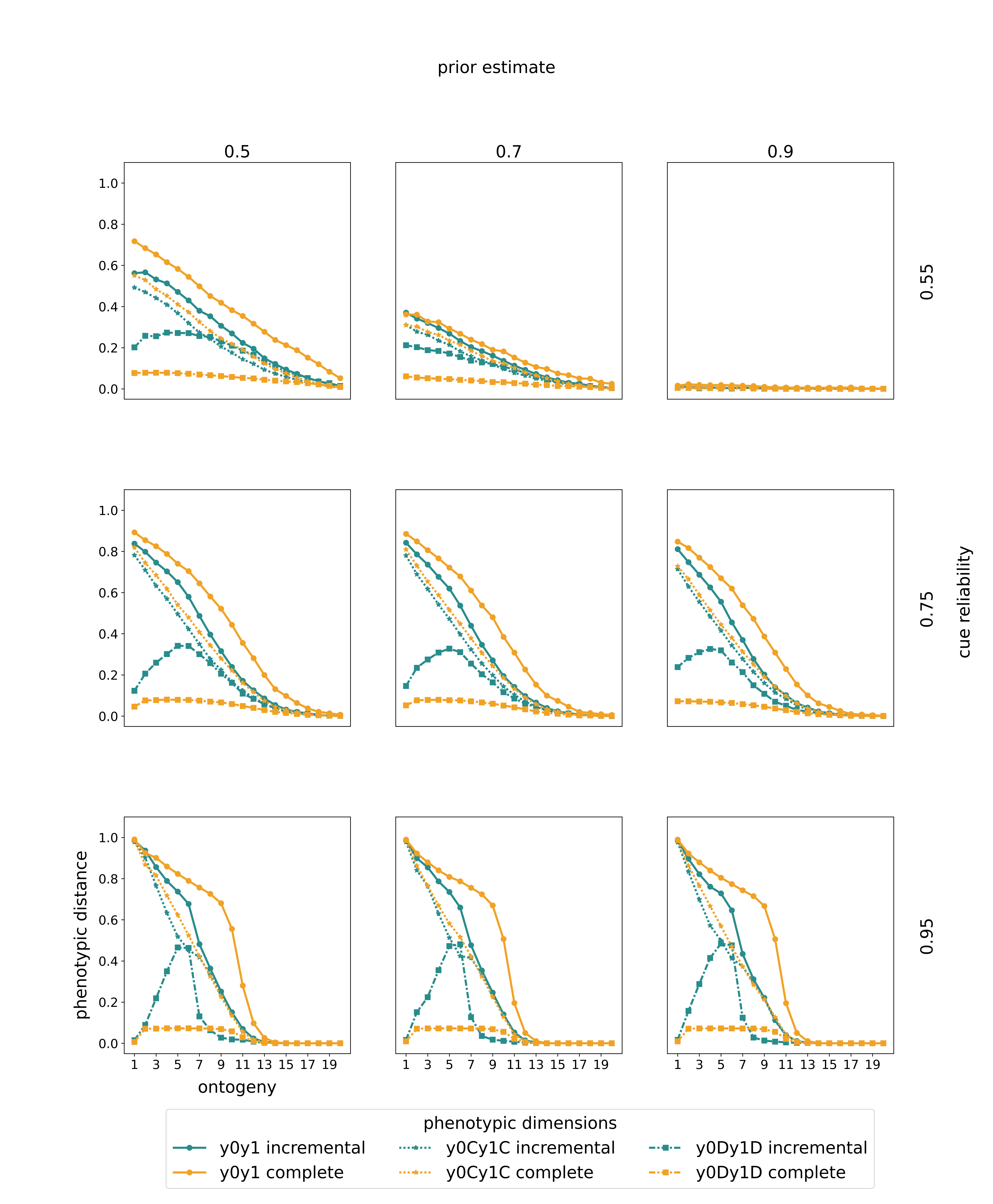

*Figure S2.54:* Changes in plasticity. Changes in plasticity are shown for diminishing rewards and diminishing penalties. Rows indicate the prior estimate of being in $E_{1}$ and columns indicate the cue reliability. Within each panel, the horizontal axis denotes ontogeny. The vertical axis denotes the normalized average phenotypic distance (our plasticity measure) across 10.000 pairs of simulated clones measured at the end of ontogeny. A specific point on any of the curves corresponds to a simulation experiment in which clones have been separated at the time point indicated by the horizontal axis. We show results from a model with incremental deconstruction in teal and with complete deconstruction in yellow. For each mode of deconstruction, we present three types of plasticity curves. First, we show plasticity in construction (dotted lines and stars) as the Euclidean distance between the number of time points spent constructing. Second, we show plasticity in deconstruction (dash-dotted lines and squares) as the Euclidean distance between the number of time points spent deconstruction. Third, we show total plasticity (solid lines and circles) as the Euclidean distance between the number of specialization steps towards either target (after accounting for deconstruction). We normalize phenotypic distance in construction and total phenotypes by dividing by the maximally possible Euclidean distance, corresponding to $2*\sqrt{T=20}$. The normalization constant for deconstruction is $2*\sqrt{T/2=10}$.

ESM 3 - Comparison of 5 and 10 time periods of on ontogeny

*5 time periods of ontogeny*

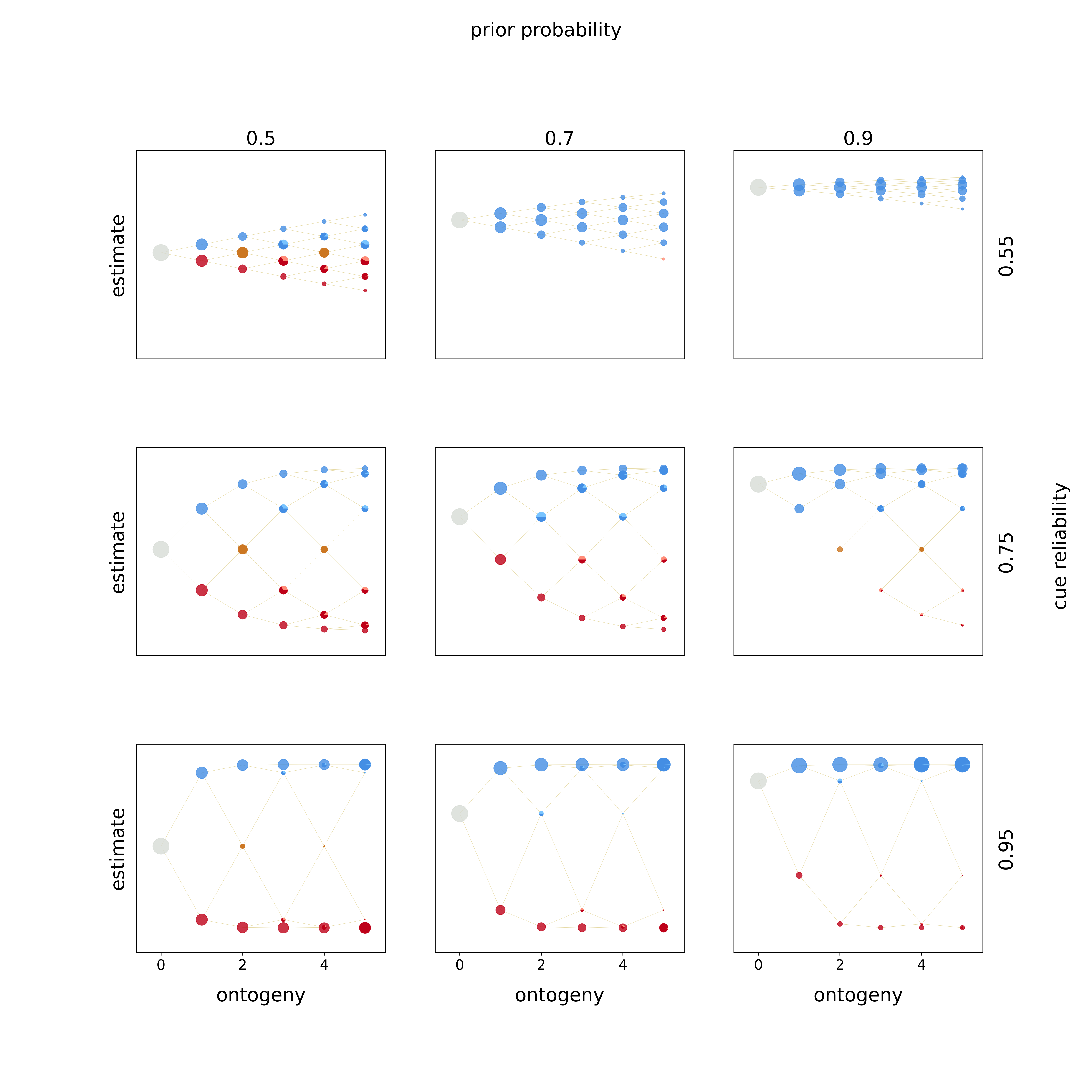

*Figure S3.1:* Optimal policies. Optimal policies are shown for a model with incremental deconstruction and linear rewards and penalties. Rows indicate the prior estimate of being in $E_{1}$ and columns indicate the cue reliability. Within each panel, the horizontal axis denotes ontogeny and the vertical axis the posterior estimates of being in $E_{1}$. The entire population starts ontogeny with zero cues sampled and the prior estimate indicated by the row (indicated by the grey circle). In each time period organisms sample a cue (either $C_{0}$ or $C_{1}$), update their estimate, and make a phenotypic decision (colored circles). Beige lines indicate developmental trajectories through this decision space, with lines branching upwards indicating the sampling of $C_{1}$ and lines branching downwards indicating the sampling of $C_{o}$. Colors denote the optimal, fitness-maximizing phenotypic choice in each state. Pies indicate cases in which organisms with the same posterior estimates make different phenotypic decisions. The area of a circle (pie piece) is proportional the probability of reaching that particular state. Colors indicate the following phenotypic decisions: Black corresponds to waiting, red to constructing $P_{0}$, blue to constructing $P_{1}$, purple to deconstructing $P_{0}$, green to deconstructing $P_{1}$, light red to a tie between constructing $P_{0}$ and deconstructing $P_{1}$, light blue to a tie between constructing $P_{1}$ and deconstructing $P_{0}$, brown to a tie between constructing either phenotypic target, yellow to a tie between deconstructing either target, grey to a tie between construction and waiting, dark grey to a tie between deconstruction and waiting, and lastly ochre to a tie between all options.

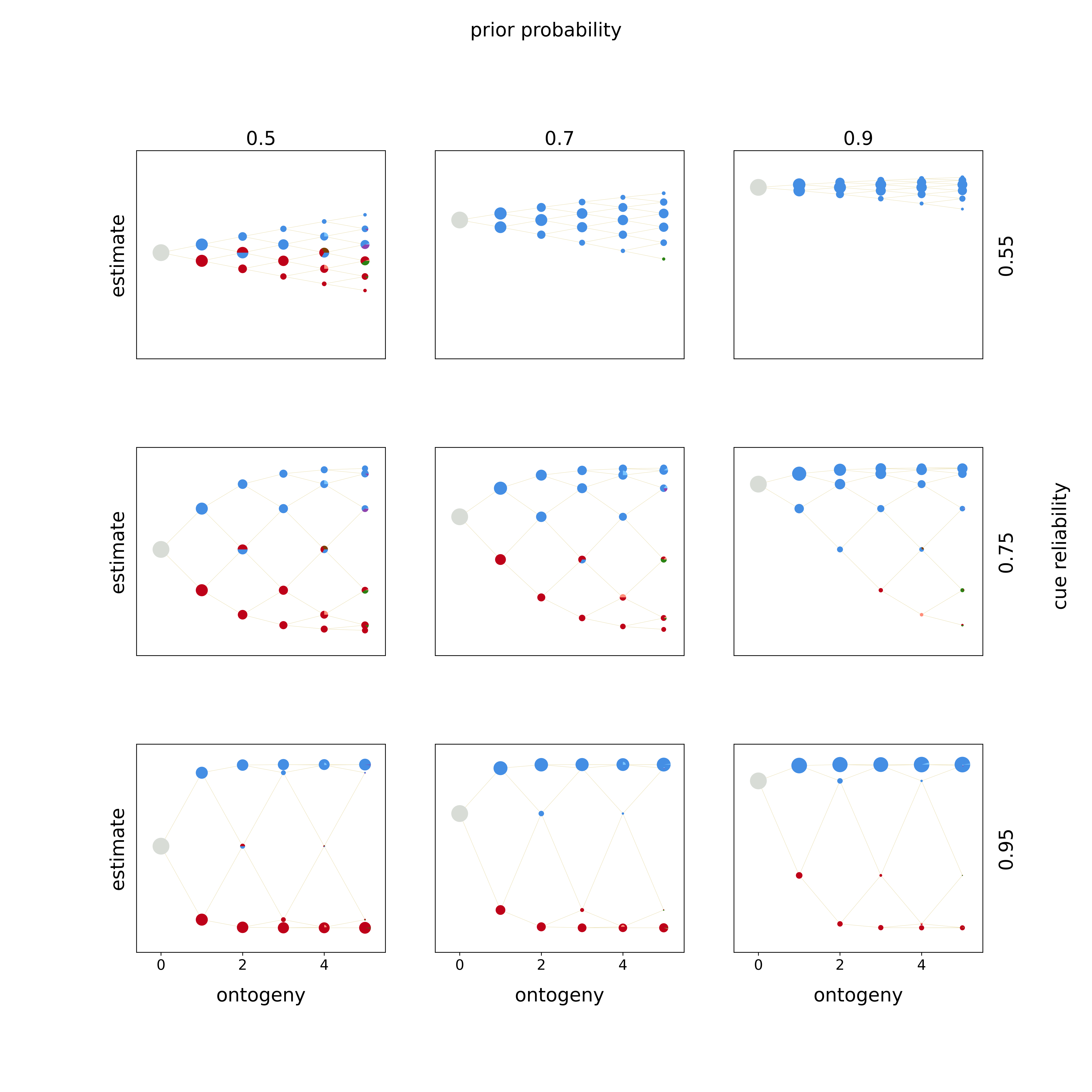

*Figure S3.2:* Optimal policies. Optimal policies are shown for a model with complete deconstruction and linear rewards and penalties. Rows indicate the prior estimate of being in $E_{1}$ and columns indicate the cue reliability. Within each panel, the horizontal axis denotes ontogeny and the vertical axis the posterior estimates of being in $E_{1}$. The entire population starts ontogeny with zero cues sampled and the prior estimate indicated by the row (indicated by the grey circle). In each time period organisms sample a cue (either $C_{0}$ or $C_{1}$), update their estimate, and make a phenotypic decision (colored circles). Beige lines indicate developmental trajectories through this decision space, with lines branching upwards indicating the sampling of $C_{1}$ and lines branching downwards indicating the sampling of $C_{o}$. Colors denote the optimal, fitness-maximizing phenotypic choice in each state. Pies indicate cases in which organisms with the same posterior estimates make different phenotypic decisions. The area of a circle (pie piece) is proportional the probability of reaching that particular state. Colors indicate the following phenotypic decisions: Black corresponds to waiting, red to constructing $P_{0}$, blue to constructing $P_{1}$, purple to deconstructing $P_{0}$, green to deconstructing $P_{1}$, light red to a tie between constructing $P_{0}$ and deconstructing $P_{1}$, light blue to a tie between constructing $P_{1}$ and deconstructing $P_{0}$, brown to a tie between constructing either phenotypic target, yellow to a tie between deconstructing either target, grey to a tie between construction and waiting, dark grey to a tie between deconstruction and waiting, and lastly ochre to a tie between all options.

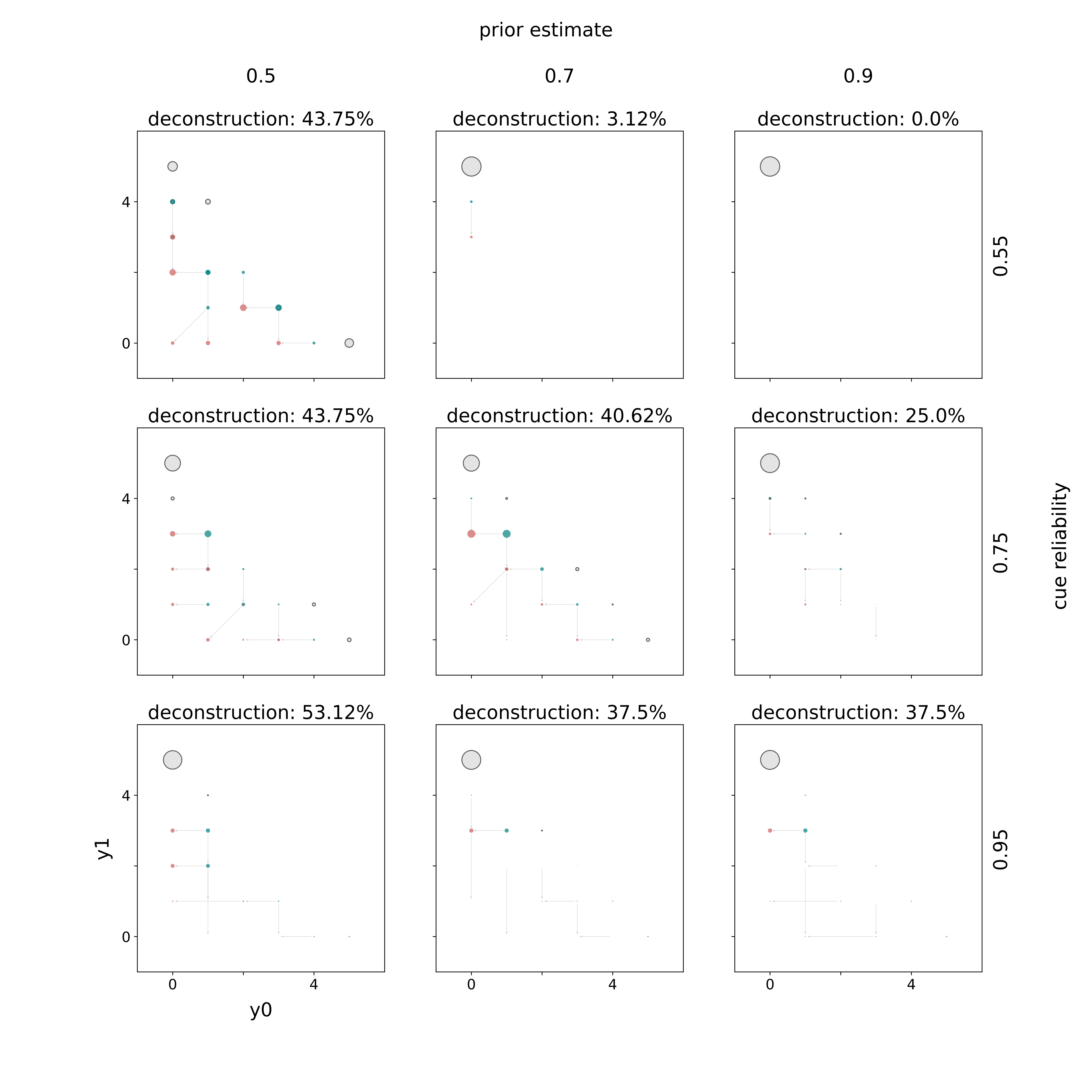

*Figure S3.3:* Distributions of mature phenotypes. Distributions of mature phenotypes are shown for a model with incremental deconstruction and linear rewards and penalties. Rows indicate the prior estimate of being in $E_{1}$ and columns indicate the cue reliability. The populations of mature phenotypes have been simulated in $E_{1}$. The title of each panel indicates the percentage of mature phenotypes that have deconstructed at some point during ontogeny. Within each panel the horizontal axis indicates the number of specializations towards $E_{0}$ and the vertical axis towards $E_{1}$. The lower triangle indicates how much mature phenotypes have constructed (teal circles) and what their phenotype looked like after deconstruction (red circles). Grey arrows connect phenotypes before (teal) and after (red) deconstruction. Grey circles with a black outline belong to mature phenotypes that never deconstructed. The area of a circle is proportional to the number mature organisms with this phenotype. The upper triangle indicates waiting. For each mature phenotype (after deconstruction) below the diagonal the corresponding square above the diagonal highlights the amount of waiting. The color intensity is proportional to the amount of waiting. Black squares indicate phenotypes that waited all of ontogeny (i.e. 20 time periods) and white squares phenotypes that never waited.

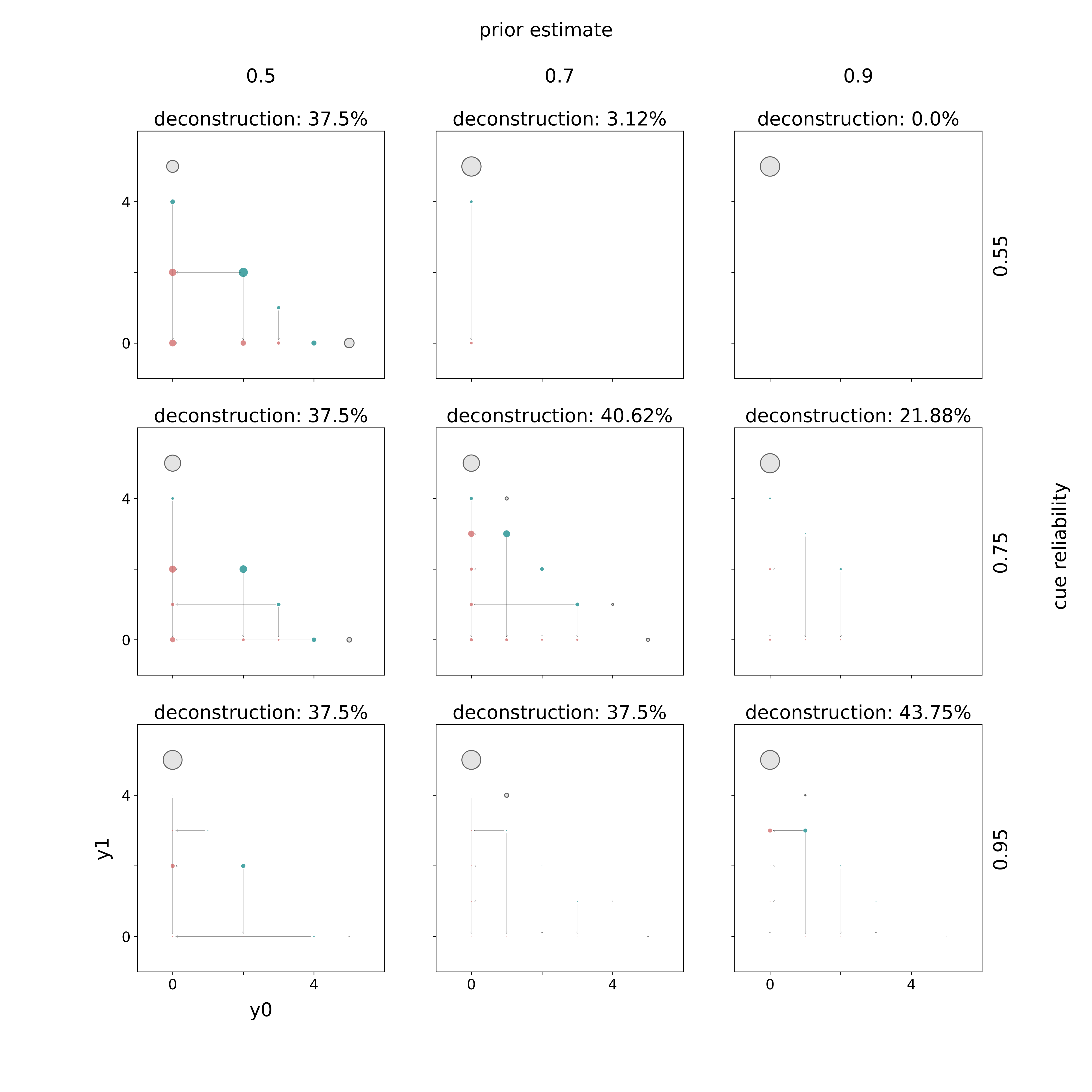

*Figure S3.4:* Distributions of mature phenotypes. Distributions of mature phenotypes are shown for a model with complete deconstruction and linear rewards and penalties. Rows indicate the prior estimate of being in $E_{1}$ and columns indicate the cue reliability. The populations of mature phenotypes have been simulated in $E_{1}$. The title of each panel indicates the percentage of mature phenotypes that have deconstructed at some point during ontogeny. Within each panel the horizontal axis indicates the number of specializations towards $E_{0}$ and the vertical axis towards $E_{1}$. The lower triangle indicates how much mature phenotypes have constructed (teal circles) and what their phenotype looked like after deconstruction (red circles). Grey arrows connect phenotypes before (teal) and after (red) deconstruction. Grey circles with a black outline belong to mature phenotypes that never deconstructed. The area of a circle is proportional to the number mature organisms with this phenotype. The upper triangle indicates waiting. For each mature phenotype (after deconstruction) below the diagonal the corresponding square above the diagonal highlights the amount of waiting. The color intensity is proportional to the amount of waiting. Black squares indicate phenotypes that waited all of ontogeny (i.e. 20 time periods) and white squares phenotypes that never waited.

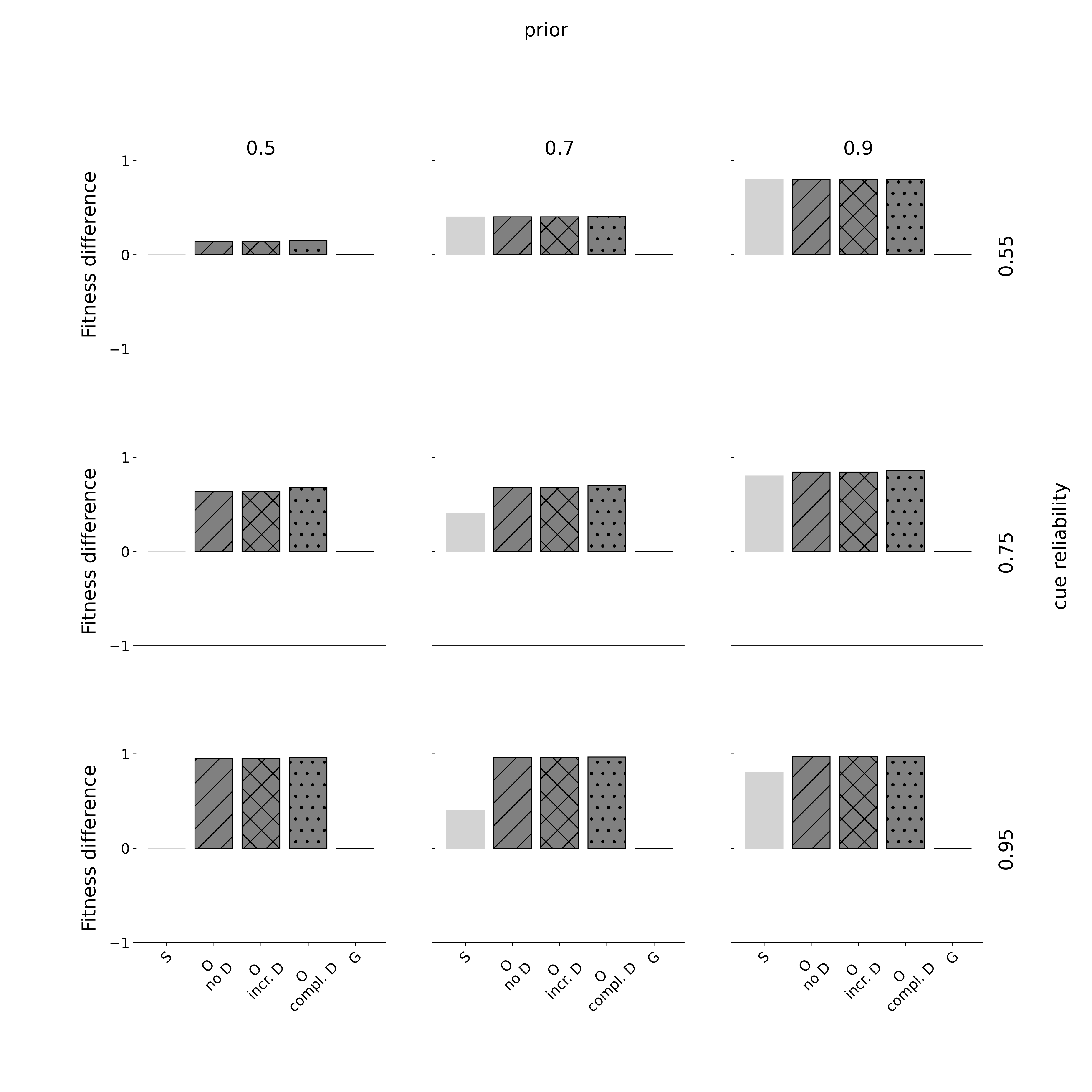

*Figure S3.5:* Fitness differences. Fitness differences are shown for linear rewards and linear penalties. Rows indicate the prior estimate of being in $E_{1}$ and columns indicate the cue reliability. Within each panel, the horizontal axis denotes the type of strategy where ‘S’ corresponds to a pure specialist strategy, ‘O’ to an optimal policy, and ‘G’ to a pure generalist strategy. The horizontal axis denotes fitness differences from baseline (corresponding to 0), normalized to range between -1 and 1. We show fitness of three different optimal policies: without deconstruction (‘no D’), with incremental deconstruction (‘incr. D’), and complete deconstruction (‘compl. D’). Specialists always fully specialize according to the prior distribution. When priors are uninformative (0.5), half the population fully specializes towards $P_{0}$ and the other one towards $P_{1}$. Generalists always specialize halfway towards either phenotypic target.

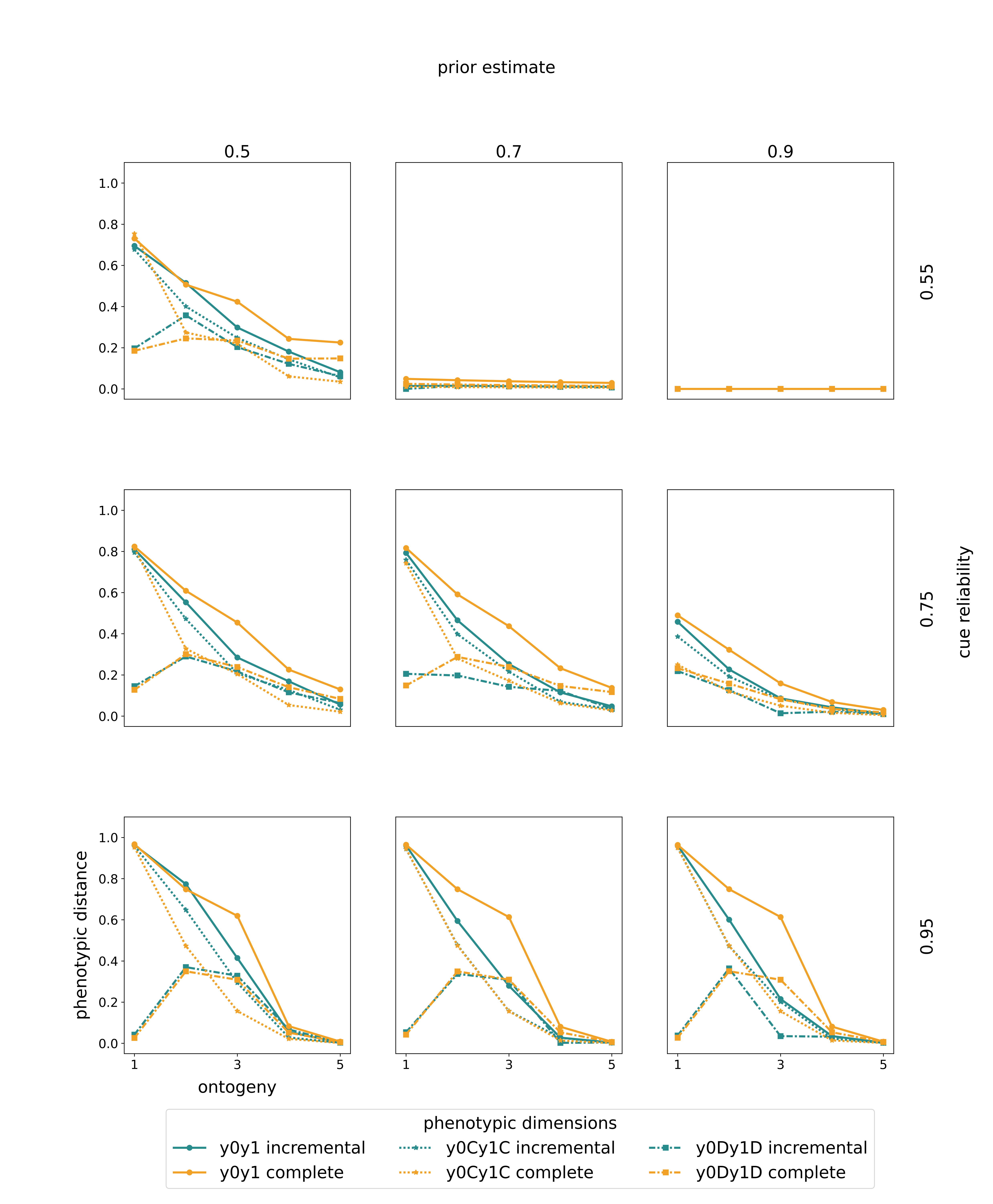

*Figure S3.6:* Changes in plasticity. Changes in plasticity are shown for linear rewards and linear penalties. Rows indicate the prior estimate of being in $E_{1}$ and columns indicate the cue reliability. Within each panel, the horizontal axis denotes ontogeny. The vertical axis denotes the normalized average phenotypic distance (our plasticity measure) across 10.000 pairs of simulated clones measured at the end of ontogeny. A specific point on any of the curves corresponds to a simulation experiment in which clones have been separated at the time point indicated by the horizontal axis. We show results from a model with incremental deconstruction in teal and with complete deconstruction in yellow. For each mode of deconstruction, we present three types of plasticity curves. First, we show plasticity in construction (dotted lines and stars) as the Euclidean distance between the number of time points spent constructing. Second, we show plasticity in deconstruction (dash-dotted lines and squares) as the Euclidean distance between the number of time points spent deconstruction. Third, we show total plasticity (solid lines and circles) as the Euclidean distance between the number of specialization steps towards either target (after accounting for deconstruction). We normalize phenotypic distance in construction and total phenotypes by dividing by the maximally possible Euclidean distance, corresponding to $2*\sqrt{T=5}$. The normalization constant for deconstruction is $2*\sqrt{T/2=2.5}$.

*10 time periods of ontogeny*

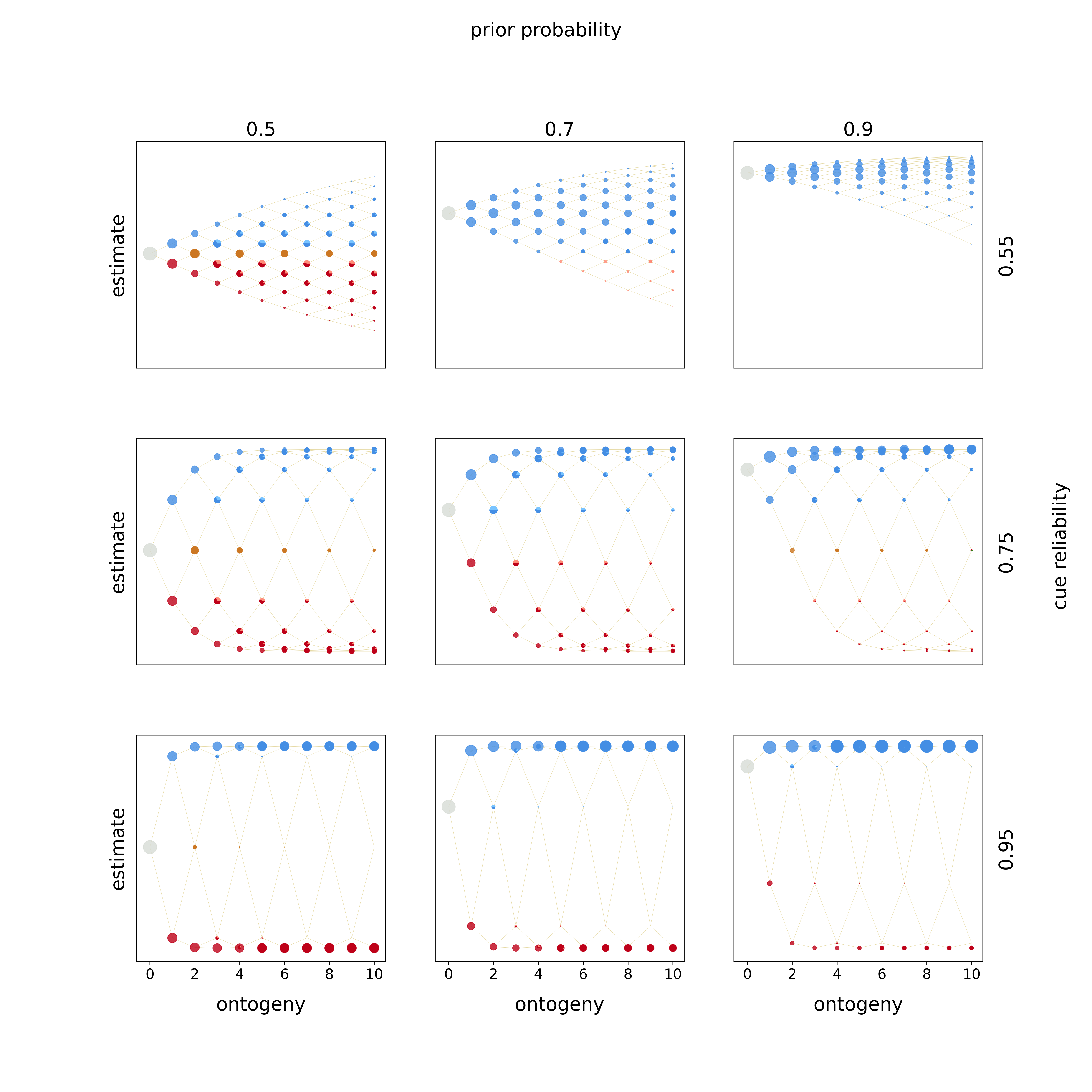

*Figure S3.7:* Optimal policies. Optimal policies are shown for a model with incremental deconstruction and linear rewards and penalties. Rows indicate the prior estimate of being in $E_{1}$ and columns indicate the cue reliability. Within each panel, the horizontal axis denotes ontogeny and the vertical axis the posterior estimates of being in $E_{1}$. The entire population starts ontogeny with zero cues sampled and the prior estimate indicated by the row (indicated by the grey circle). In each time period organisms sample a cue (either $C_{0}$ or $C_{1}$), update their estimate, and make a phenotypic decision (colored circles). Beige lines indicate developmental trajectories through this decision space, with lines branching upwards indicating the sampling of $C_{1}$ and lines branching downwards indicating the sampling of $C_{o}$. Colors denote the optimal, fitness-maximizing phenotypic choice in each state. Pies indicate cases in which organisms with the same posterior estimates make different phenotypic decisions. The area of a circle (pie piece) is proportional the probability of reaching that particular state. Colors indicate the following phenotypic decisions: Black corresponds to waiting, red to constructing $P_{0}$, blue to constructing $P_{1}$, purple to deconstructing $P_{0}$, green to deconstructing $P_{1}$, light red to a tie between constructing $P_{0}$ and deconstructing $P_{1}$, light blue to a tie between constructing $P_{1}$ and deconstructing $P_{0}$, brown to a tie between constructing either phenotypic target, yellow to a tie between deconstructing either target, grey to a tie between construction and waiting, dark grey to a tie between deconstruction and waiting, and lastly ochre to a tie between all options.

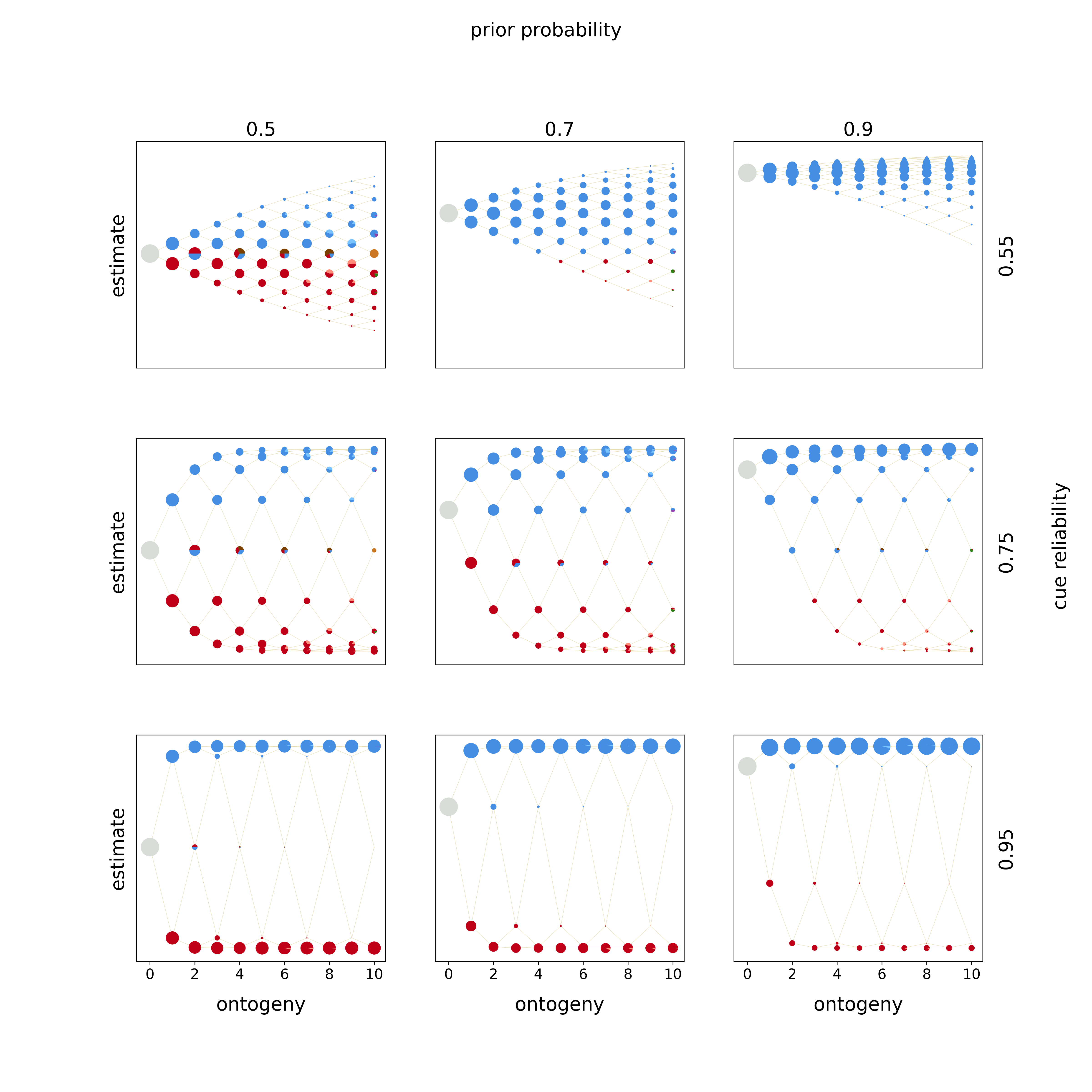

*Figure S3.8:* Optimal policies. Optimal policies are shown for a model with complete deconstruction and linear rewards and penalties. Rows indicate the prior estimate of being in $E_{1}$ and columns indicate the cue reliability. Within each panel, the horizontal axis denotes ontogeny and the vertical axis the posterior estimates of being in $E_{1}$. The entire population starts ontogeny with zero cues sampled and the prior estimate indicated by the row (indicated by the grey circle). In each time period organisms sample a cue (either $C_{0}$ or $C_{1}$), update their estimate, and make a phenotypic decision (colored circles). Beige lines indicate developmental trajectories through this decision space, with lines branching upwards indicating the sampling of $C_{1}$ and lines branching downwards indicating the sampling of $C_{o}$. Colors denote the optimal, fitness-maximizing phenotypic choice in each state. Pies indicate cases in which organisms with the same posterior estimates make different phenotypic decisions. The area of a circle (pie piece) is proportional the probability of reaching that particular state. Colors indicate the following phenotypic decisions: Black corresponds to waiting, red to constructing $P_{0}$, blue to constructing $P_{1}$, purple to deconstructing $P_{0}$, green to deconstructing $P_{1}$, light red to a tie between constructing $P_{0}$ and deconstructing $P_{1}$, light blue to a tie between constructing $P_{1}$ and deconstructing $P_{0}$, brown to a tie between constructing either phenotypic target, yellow to a tie between deconstructing either target, grey to a tie between construction and waiting, dark grey to a tie between deconstruction and waiting, and lastly ochre to a tie between all options.

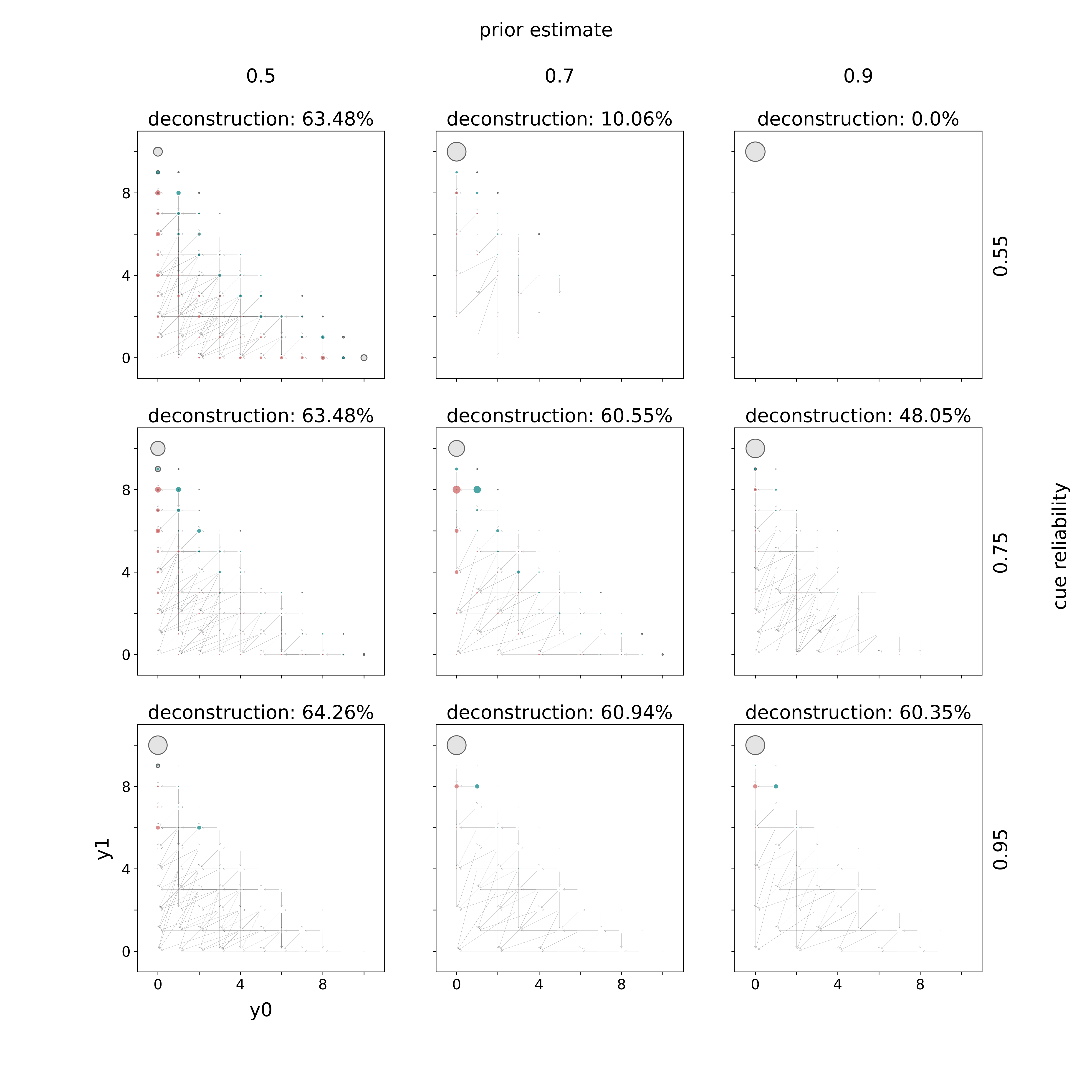

*Figure S3.9:* Distributions of mature phenotypes. Distributions of mature phenotypes are shown for a model with incremental deconstruction and linear rewards and penalties. Rows indicate the prior estimate of being in $E_{1}$ and columns indicate the cue reliability. The populations of mature phenotypes have been simulated in $E_{1}$. The title of each panel indicates the percentage of mature phenotypes that have deconstructed at some point during ontogeny. Within each panel the horizontal axis indicates the number of specializations towards $E_{0}$ and the vertical axis towards $E_{1}$. The lower triangle indicates how much mature phenotypes have constructed (teal circles) and what their phenotype looked like after deconstruction (red circles). Grey arrows connect phenotypes before (teal) and after (red) deconstruction. Grey circles with a black outline belong to mature phenotypes that never deconstructed. The area of a circle is proportional to the number mature organisms with this phenotype. The upper triangle indicates waiting. For each mature phenotype (after deconstruction) below the diagonal the corresponding square above the diagonal highlights the amount of waiting. The color intensity is proportional to the amount of waiting. Black squares indicate phenotypes that waited all of ontogeny (i.e. 20 time periods) and white squares phenotypes that never waited.

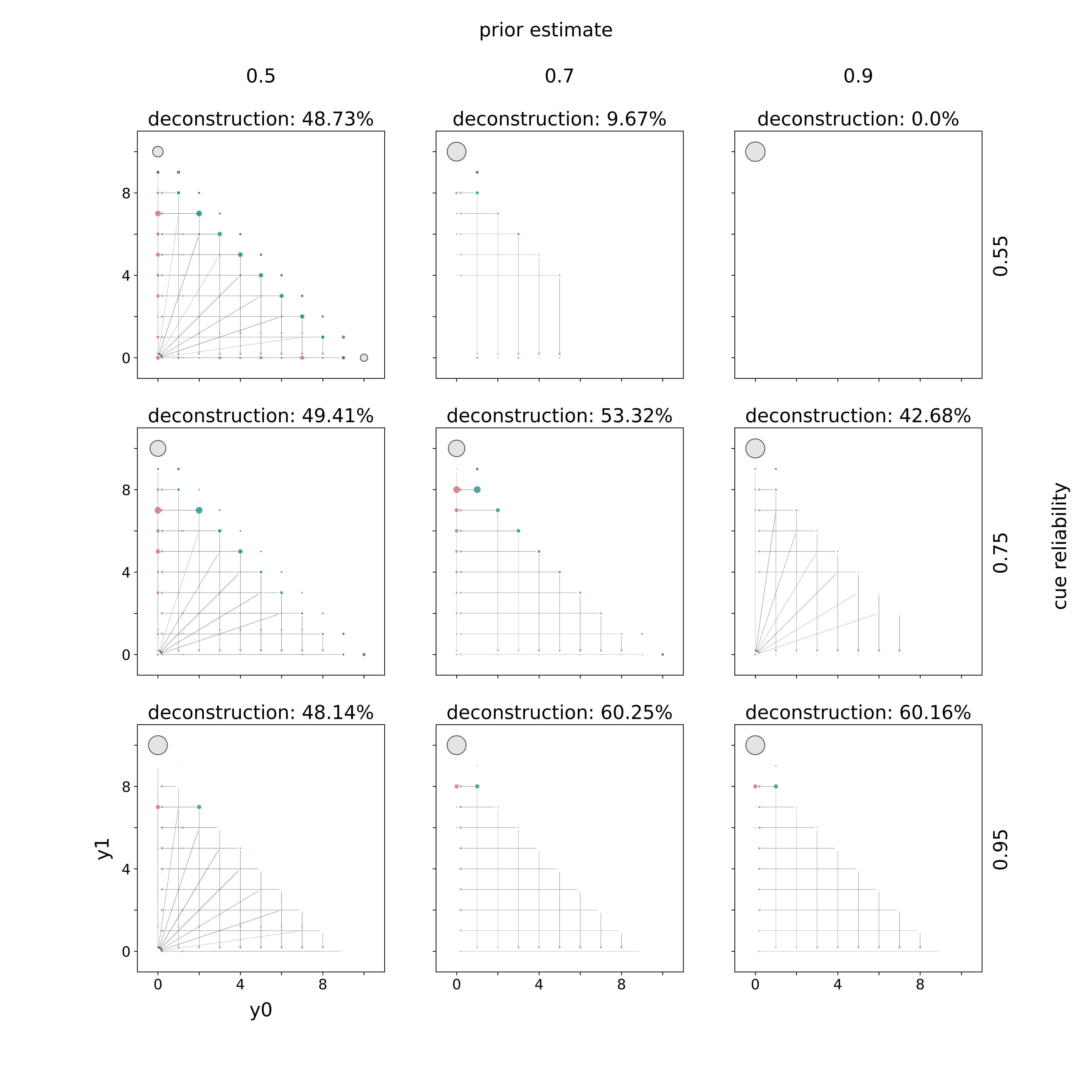

*Figure S3.10:* Distributions of mature phenotypes. Distributions of mature phenotypes are shown for a model with complete deconstruction and linear rewards and penalties. Rows indicate the prior estimate of being in $E_{1}$ and columns indicate the cue reliability. The populations of mature phenotypes have been simulated in $E_{1}$. The title of each panel indicates the percentage of mature phenotypes that have deconstructed at some point during ontogeny. Within each panel the horizontal axis indicates the number of specializations towards $E_{0}$ and the vertical axis towards $E_{1}$. The lower triangle indicates how much mature phenotypes have constructed (teal circles) and what their phenotype looked like after deconstruction (red circles). Grey arrows connect phenotypes before (teal) and after (red) deconstruction. Grey circles with a black outline belong to mature phenotypes that never deconstructed. The area of a circle is proportional to the number mature organisms with this phenotype. The upper triangle indicates waiting. For each mature phenotype (after deconstruction) below the diagonal the corresponding square above the diagonal highlights the amount of waiting. The color intensity is proportional to the amount of waiting. Black squares indicate phenotypes that waited all of ontogeny (i.e. 20 time periods) and white squares phenotypes that never waited.

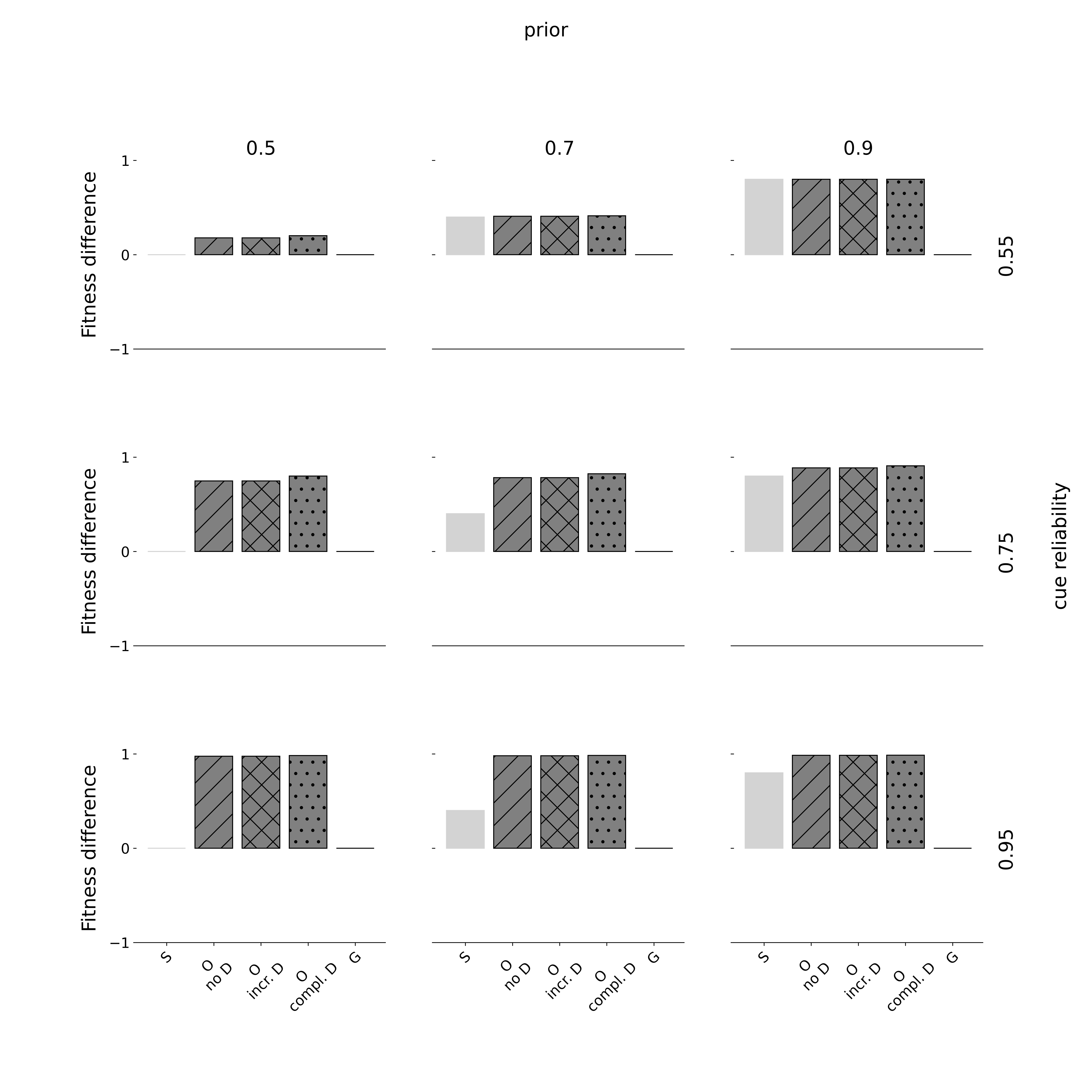

*Figure S3.11:* Fitness differences. Fitness differences are shown for linear rewards and linear penalties. Rows indicate the prior estimate of being in $E_{1}$ and columns indicate the cue reliability. Within each panel, the horizontal axis denotes the type of strategy where ‘S’ corresponds to a pure specialist strategy, ‘O’ to an optimal policy, and ‘G’ to a pure generalist strategy. The horizontal axis denotes fitness differences from baseline (corresponding to 0), normalized to range between -1 and 1. We show fitness of three different optimal policies: without deconstruction (‘no D’), with incremental deconstruction (‘incr. D’), and complete deconstruction (‘compl. D’). Specialists always fully specialize according to the prior distribution. When priors are uninformative (0.5), half the population fully specializes towards $P_{0}$ and the other one towards $P_{1}$. Generalists always specialize halfway towards either phenotypic target.

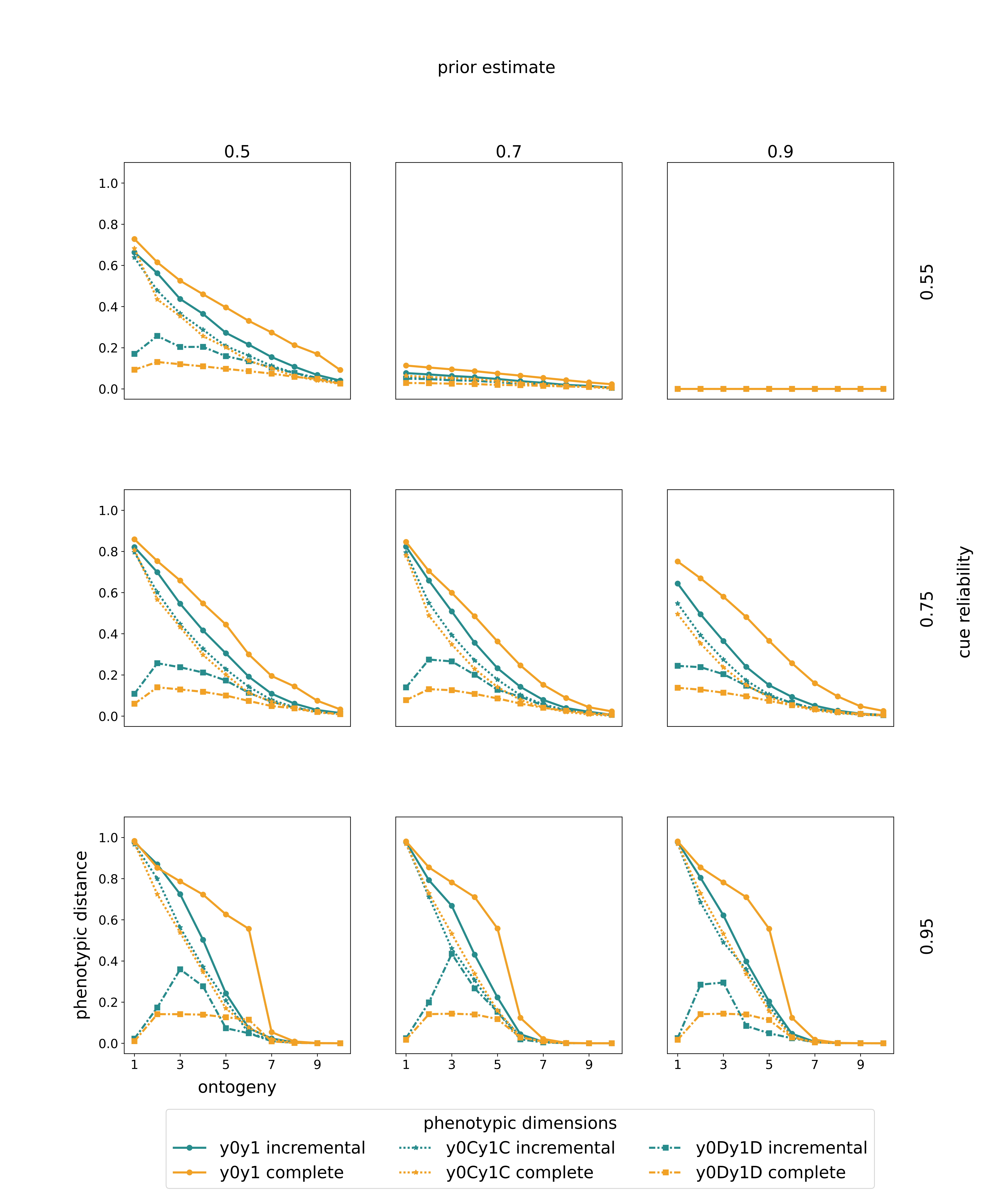

*Figure S3.12:* Changes in plasticity. Changes in plasticity are shown for linear rewards and linear penalties. Rows indicate the prior estimate of being in $E_{1}$ and columns indicate the cue reliability. Within each panel, the horizontal axis denotes ontogeny. The vertical axis denotes the normalized average phenotypic distance (our plasticity measure) across 10.000 pairs of simulated clones measured at the end of ontogeny. A specific point on any of the curves corresponds to a simulation experiment in which clones have been separated at the time point indicated by the horizontal axis. We show results from a model with incremental deconstruction in teal and with complete deconstruction in yellow. For each mode of deconstruction, we present three types of plasticity curves. First, we show plasticity in construction (dotted lines and stars) as the Euclidean distance between the number of time points spent constructing. Second, we show plasticity in deconstruction (dash-dotted lines and squares) as the Euclidean distance between the number of time points spent deconstruction. Third, we show total plasticity (solid lines and circles) as the Euclidean distance between the number of specialization steps towards either target (after accounting for deconstruction). We normalize phenotypic distance in construction and total phenotypes by dividing by the maximally possible Euclidean distance, corresponding to $2*\sqrt{T=10}$. The normalization constant for deconstruction is $2*\sqrt{T/2=5}$.

ESM 4 - Different paradigms for quantifying plasticity (linear rewards and penalties)

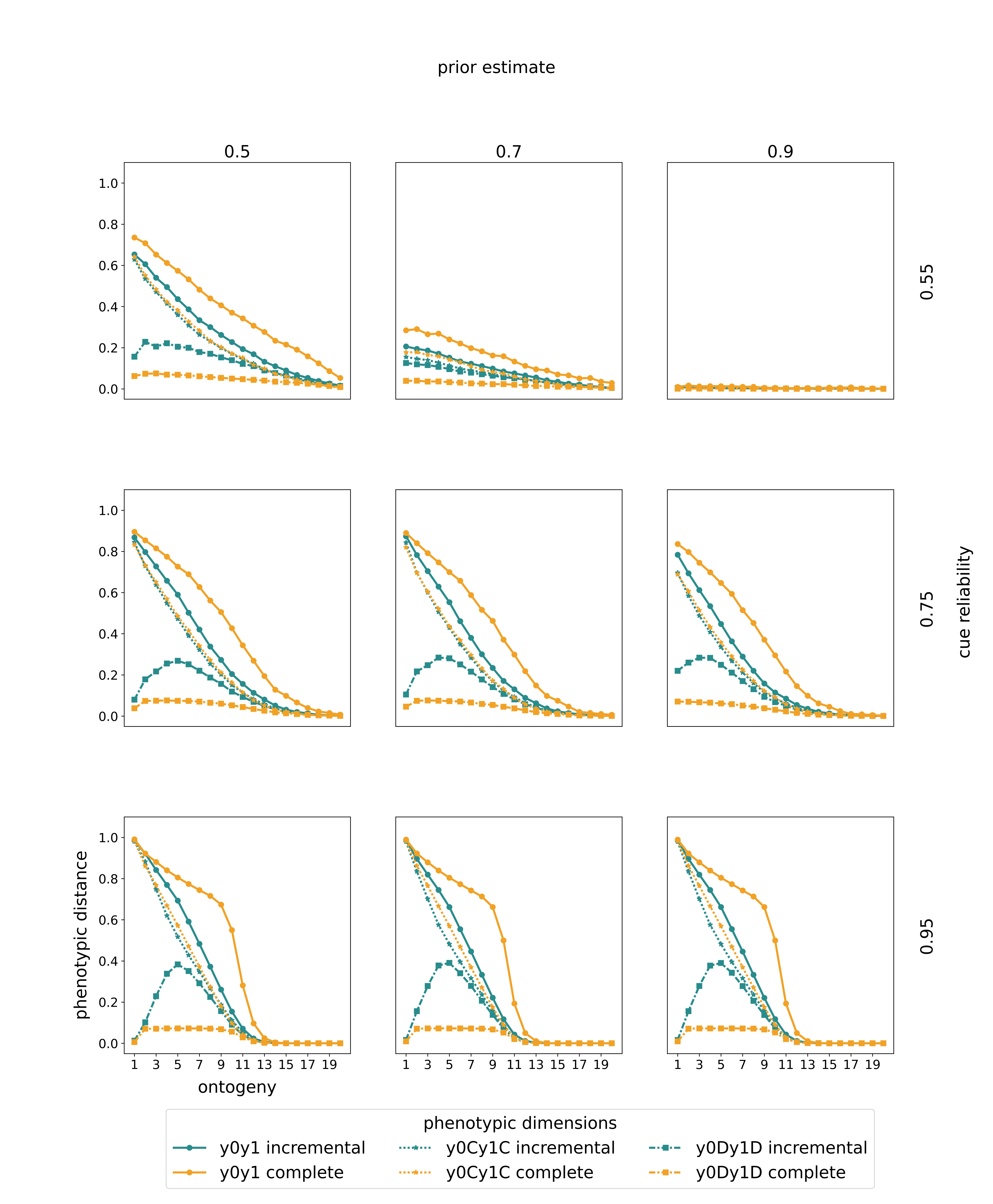

*Figure S4.1:* Changes in plasticity. Changes in plasticity are shown for linear rewards and linear penalties when clones receive reciprocal, opposite cues. Rows indicate the prior estimate of being in $E_{1}$ and columns indicate the cue reliability. Within each panel, the horizontal axis denotes ontogeny. The vertical axis denotes the normalized average phenotypic distance (our plasticity measure) across 10.000 pairs of simulated clones measured at the end of ontogeny. A specific point on any of the curves corresponds to a simulation experiment in which clones have been separated at the time point indicated by the horizontal axis. We show results from a model with incremental deconstruction in teal and with complete deconstruction in yellow. For each mode of deconstruction, we present three types of plasticity curves. First, we show plasticity in construction (dotted lines and stars) as the Euclidean distance between the number of time points spent constructing. Second, we show plasticity in deconstruction (dash-dotted lines and squares) as the Euclidean distance between the number of time points spent deconstruction. Third, we show total plasticity (solid lines and circles) as the Euclidean distance between the number of specialization steps towards either target (after accounting for deconstruction). We normalize phenotypic distance in construction and total phenotypes by dividing by the maximally possible Euclidean distance, corresponding to $2*\sqrt{T=20}$. The normalization constant for deconstruction is $2*\sqrt{T/2=10}$.

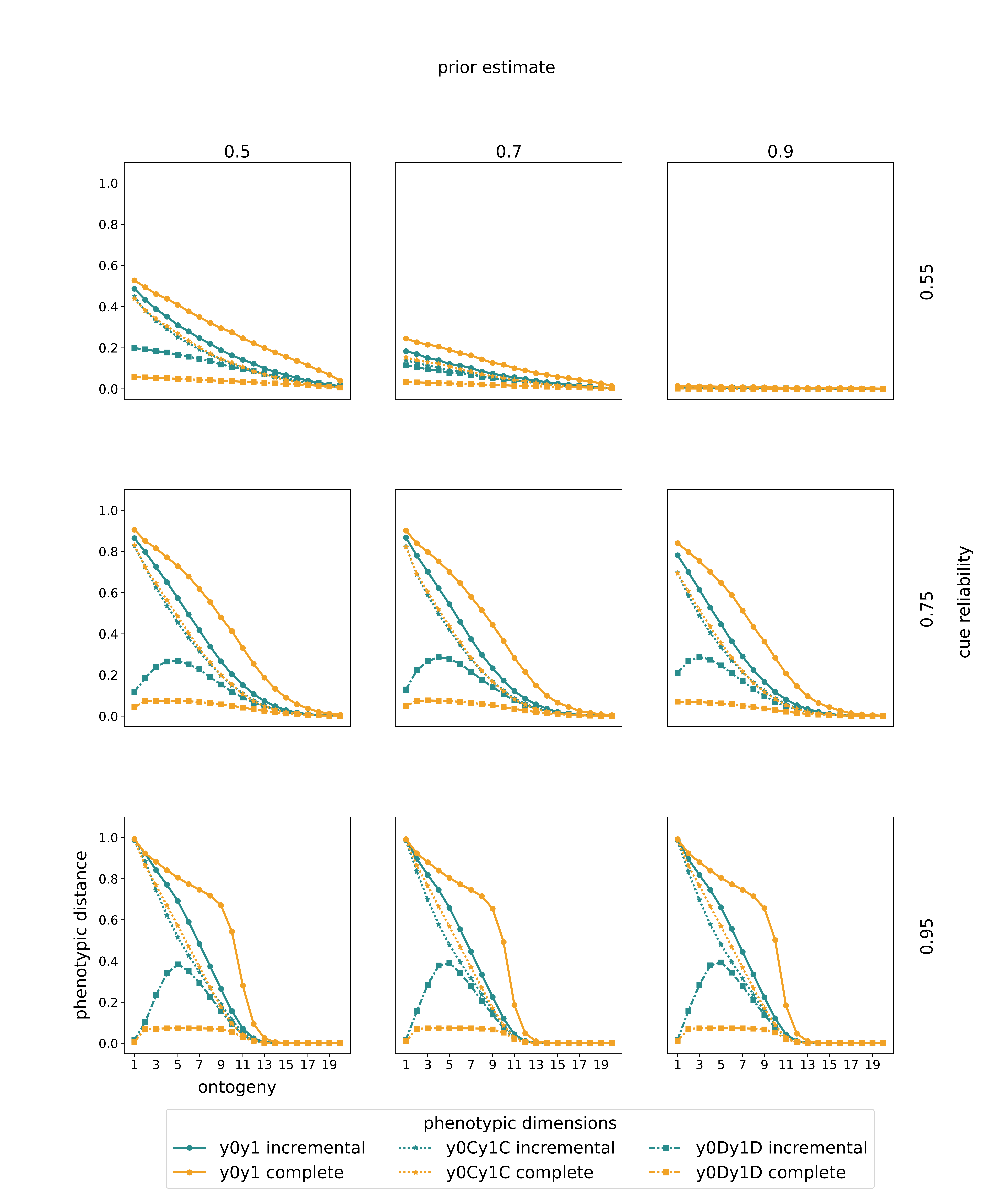

*Figure S4.2:* Changes in plasticity. Changes in plasticity are shown for linear rewards and linear penalties when clones receive cues from the opposite patch. Rows indicate the prior estimate of being in $E_{1}$ and columns indicate the cue reliability. Within each panel, the horizontal axis denotes ontogeny. The vertical axis denotes the normalized average phenotypic distance (our plasticity measure) across 10.000 pairs of simulated clones measured at the end of ontogeny. A specific point on any of the curves corresponds to a simulation experiment in which clones have been separated at the time point indicated by the horizontal axis. We show results from a model with incremental deconstruction in teal and with complete deconstruction in yellow. For each mode of deconstruction, we present three types of plasticity curves. First, we show plasticity in construction (dotted lines and stars) as the Euclidean distance between the number of time points spent constructing. Second, we show plasticity in deconstruction (dash-dotted lines and squares) as the Euclidean distance between the number of time points spent deconstruction. Third, we show total plasticity (solid lines and circles) as the Euclidean distance between the number of specialization steps towards either target (after accounting for deconstruction). We normalize phenotypic distance in construction and total phenotypes by dividing by the maximally possible Euclidean distance, corresponding to $2*\sqrt{T=20}$. The normalization constant for deconstruction is $2*\sqrt{T/2=10}$.

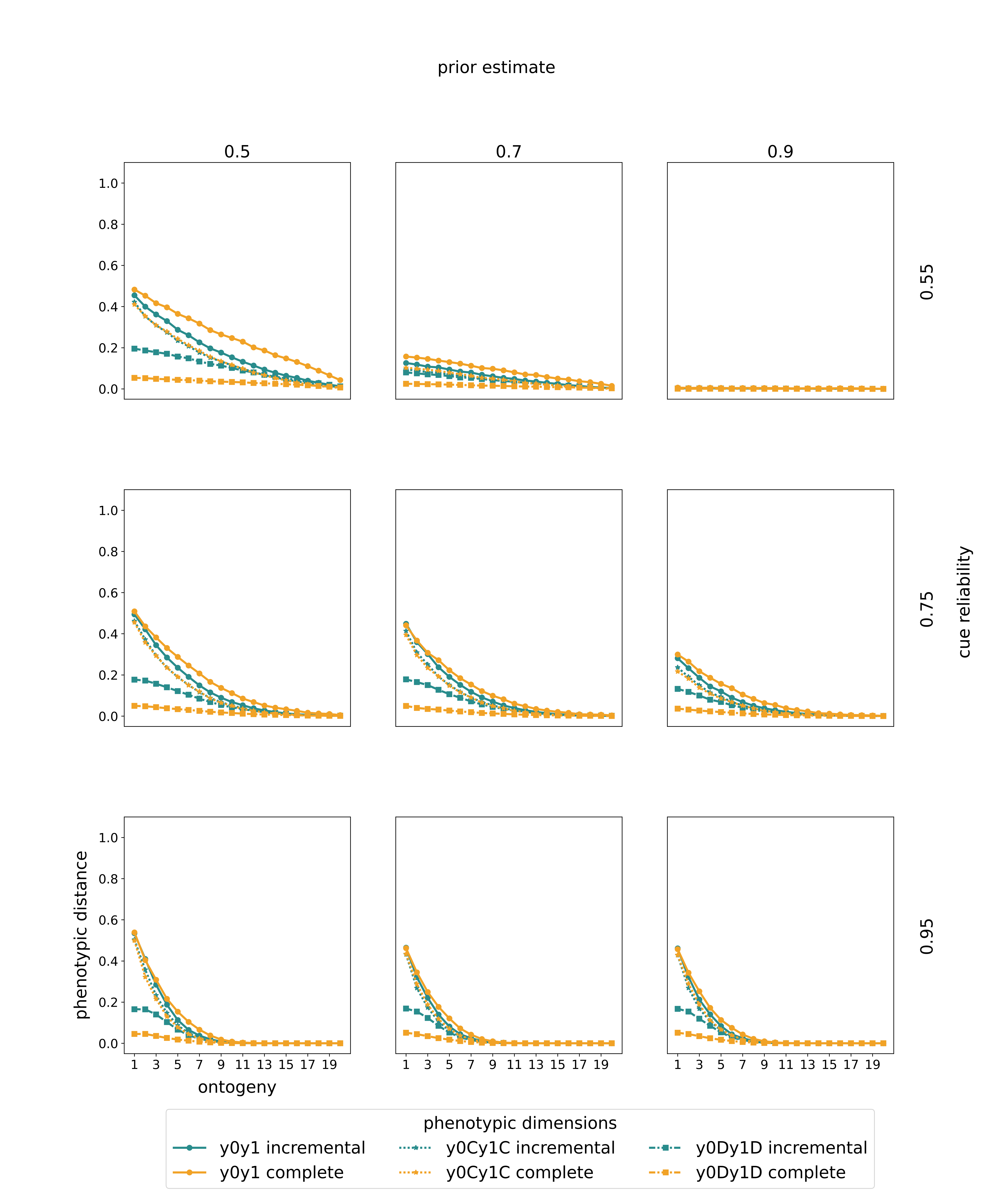

*Figure S4.3:* Changes in plasticity. Changes in plasticity are shown for linear rewards and linear penalties when clones receive uninformative cues (‘deprivation’). Rows indicate the prior estimate of being in $E_{1}$ and columns indicate the cue reliability. Within each panel, the horizontal axis denotes ontogeny. The vertical axis denotes the normalized average phenotypic distance (our plasticity measure) across 10.000 pairs of simulated clones measured at the end of ontogeny. A specific point on any of the curves corresponds to a simulation experiment in which clones have been separated at the time point indicated by the horizontal axis. We show results from a model with incremental deconstruction in teal and with complete deconstruction in yellow. For each mode of deconstruction, we present three types of plasticity curves. First, we show plasticity in construction (dotted lines and stars) as the Euclidean distance between the number of time points spent constructing. Second, we show plasticity in deconstruction (dash-dotted lines and squares) as the Euclidean distance between the number of time points spent deconstruction. Third, we show total plasticity (solid lines and circles) as the Euclidean distance between the number of specialization steps towards either target (after accounting for deconstruction). We normalize phenotypic distance in construction and total phenotypes by dividing by the maximally possible Euclidean distance, corresponding to $2*\sqrt{T=20}$. The normalization constant for deconstruction is $2*\sqrt{T/2=10}$.

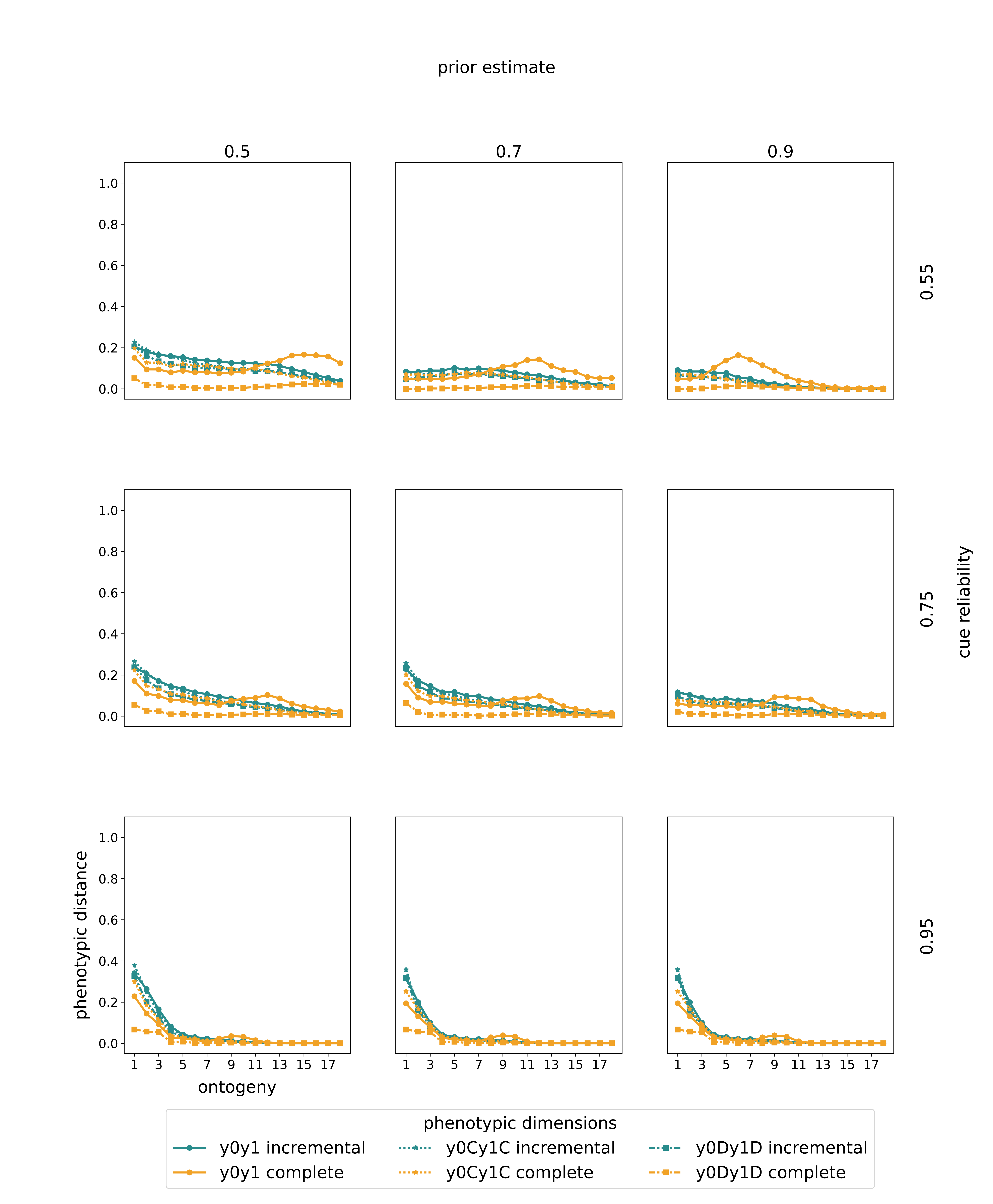

*Figure S4.4:* Changes in plasticity. Changes in plasticity are shown for linear rewards and linear penalties when clones receive reciprocal, opposite cues during a temporary separation period of 3 time periods. Phenotypic differences are measured at the end of ontogeny. Rows indicate the prior estimate of being in $E_{1}$ and columns indicate the cue reliability. Within each panel, the horizontal axis denotes ontogeny. The vertical axis denotes the normalized average phenotypic distance (our plasticity measure) across 10.000 pairs of simulated clones measured at the end of ontogeny. A specific point on any of the curves corresponds to a simulation experiment in which clones have been separated at the time point indicated by the horizontal axis. We show results from a model with incremental deconstruction in teal and with complete deconstruction in yellow. For each mode of deconstruction, we present three types of plasticity curves. First, we show plasticity in construction (dotted lines and stars) as the Euclidean distance between the number of time points spent constructing. Second, we show plasticity in deconstruction (dash-dotted lines and squares) as the Euclidean distance between the number of time points spent deconstruction. Third, we show total plasticity (solid lines and circles) as the Euclidean distance between the number of specialization steps towards either target (after accounting for deconstruction). We normalize phenotypic distance in construction and total phenotypes by dividing by the maximally possible Euclidean distance, corresponding to $2*\sqrt{T=20}$. The normalization constant for deconstruction is $2*\sqrt{T/2=10}$.

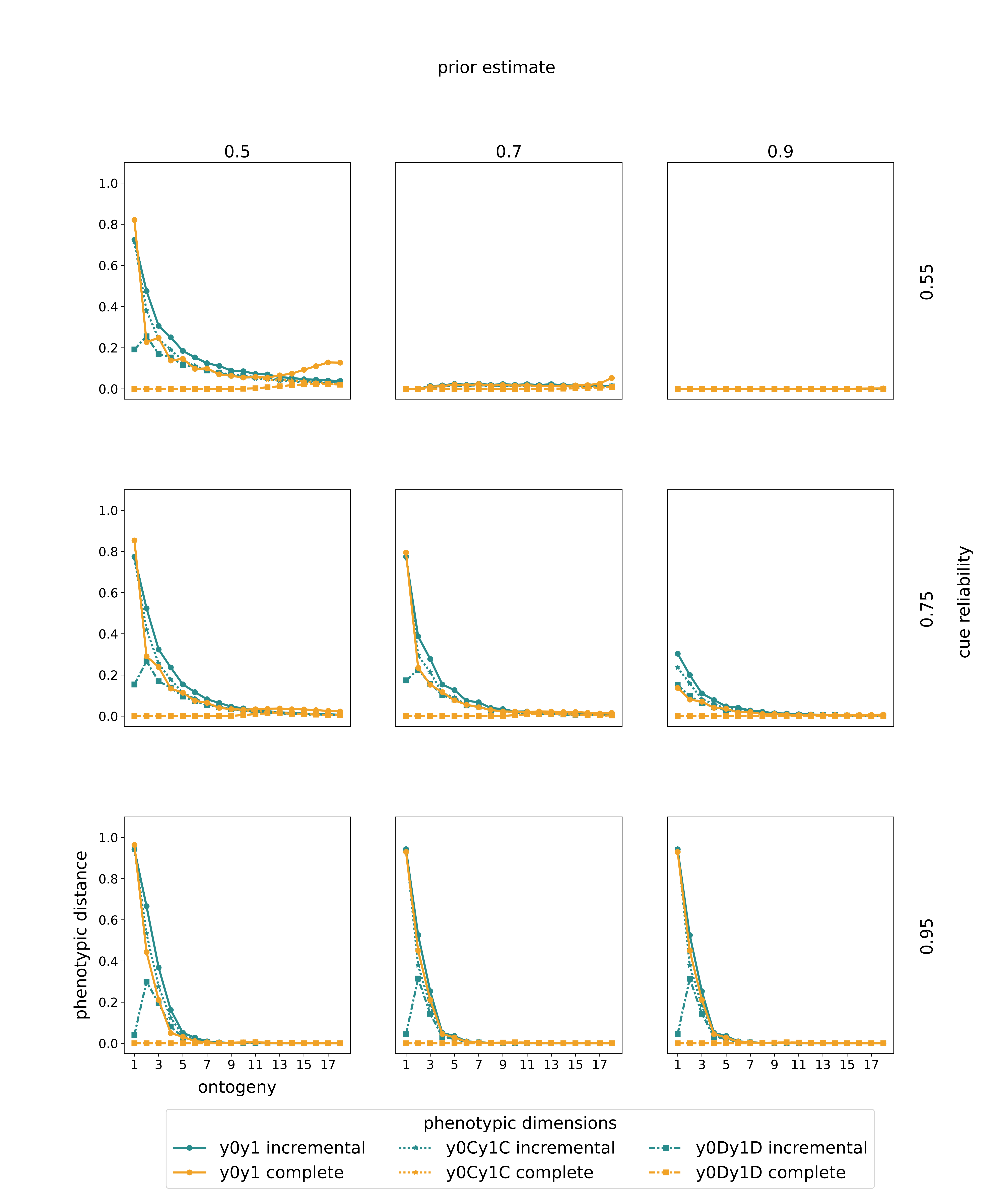

*Figure S4.5:* Changes in plasticity. Changes in plasticity are shown for linear rewards and linear penalties when clones receive reciprocal, opposite cues during a temporary separation period of three time periods. Phenotypic differences are measured right after the separation. Rows indicate the prior estimate of being in $E_{1}$ and columns indicate the cue reliability. Within each panel, the horizontal axis denotes ontogeny. The vertical axis denotes the normalized average phenotypic distance (our plasticity measure) across 10.000 pairs of simulated clones measured at the end of ontogeny. A specific point on any of the curves corresponds to a simulation experiment in which clones have been separated at the time point indicated by the horizontal axis. We show results from a model with incremental deconstruction in teal and with complete deconstruction in yellow. For each mode of deconstruction, we present three types of plasticity curves. First, we show plasticity in construction (dotted lines and stars) as the Euclidean distance between the number of time points spent constructing. Second, we show plasticity in deconstruction (dash-dotted lines and squares) as the Euclidean distance between the number of time points spent deconstruction. Third, we show total plasticity (solid lines and circles) as the Euclidean distance between the number of specialization steps towards either target (after accounting for deconstruction). We normalize phenotypic distance in construction and total phenotypes by dividing by the maximally possible Euclidean distance, corresponding to $2*\sqrt{time period of separation+3}$. The normalization constant for deconstruction is $2*\sqrt{time period of separation+3}$.
